# Adaptive predictor-set linear model: an imputation-free method for linear regression prediction on datasets with missing values

**DOI:** 10.1101/2024.02.19.581027

**Authors:** Benjamin Planterose Jiménez, Manfred Kayser, Athina Vidaki, Amke Caliebe

## Abstract

Linear regression (LR) is vastly used in data analysis for continuous outcomes in biomedicine and epidemiology. Despite its popularity, LR is incompatible with missing data, which frequently occur in health sciences. For parameter estimation, this short-coming is usually resolved by complete-case analysis or imputation. Both workarounds, however, are inadequate for prediction, since they either fail to predict on incomplete records or ignore missingness-induced reduction in prediction accuracy and rely on (unrealistic) assumptions about the missing mechanism. Here, we derive adaptive predictor-set linear model (aps-lm), capable of making predictions for incomplete data without the need for imputation. It is derived by using a predictor-selection operation, the Moore-Penrose pseudoinverse and the reduced QR-decomposition. aps-lm is an LR generalization that inherently handles missing values. It is applied on a reference dataset, where complete predictors and outcome are available, and yields a set of privacy-preserving parameters. In a second stage, these are shared for making predictions of the outcome on external datasets with missing entries for predictors without imputation. Moreover, aps-lm computes prediction errors that account for the pattern of missing values even under extreme missingness. We benchmark aps-lm in a simulation study. aps-lm showed greater prediction accuracy and reduced bias compared to popular imputation strategies under a wide range of scenarios including variation of sample size, goodness-of-fit, missing value type and covariance structure. Finally, as a proof-of-principle, we apply aps-lm in the context of epigenetic aging clocks, linear models that predict a person’s biological age from epigenetic data with promising clinical applications.

## 1 INTRODUCTION

Due to its simplicity and well-known properties, linear regression (LR) is a frequently applied statistical model in biomedical sciences when performing statistical inference and prediction of a continuous outcome given a set of predictors. Even though LR requires strong assumptions such as normally-distributed residuals, violations can in practice often be tolerated without compromising its validity, especially at large sample sizes ^1^. Two different statistical problems have to be differentiated when it comes to LR: On the one hand there is parameter estimation of the regression coefficients. On the other hand there is the aim to predict the outcome of the LR from new data of the predictors given the model with estimated parameters.

Missing values are a common phenomenon in empirical sciences in the Age of Information ^2^, and can be a threat to the integrity of data analysis in life and health sciences ^3,4^. Refined prediction models for disease risk or treatment response often incorporate a large number of clinical influence or risk variables, and thus make incomplete records likely. In the past, research has focused on how LR can be adapted to tolerate missing values in the context of parameter estimation ^5^. Two commonly applied approaches are complete-case analysis (CCA) or imputation, in which incomplete observations are either removed or replaced by complete, partially imputed values, respectively. In contrast to this, adaptations of LR that can deal with missing values under the prediction paradigm have barely been explored ^6,7,8^ which is especially problematic because both CCA and imputation are sub-optimal solutions in this situation ^9^: CCA implies that predictions can only be made for complete observations. Incomplete medical records are common and CCA will result in the inability to draw predictive outcomes for certain patients, an especially heavy loss when the data was collected via invasive or expensive methodologies ^10^. Moreover, CCA can have unintended consequences: for instance, minority groups are more reluctant to give information and thus, more likely to generate incomplete records. Failing to embrace the missingness in our modern datasets may thus lead to a fairness reduction in our models ^2^. Alternatively, aiming to include incomplete observations, imputation is typically employed prior to calculating predictions ^11^. For this, however, additional assumptions are required, namely, missing completely at random (MCAR) or missing at random (MAR) ^12^, which are seldom realistic scenarios in practice. In fact, the missing data mechanism is rarely known and cannot be tested for, with the exception of MCAR ^13^. Imputation may thus introduce sources of bias or increase the computational burden, especially in the case of multiple imputation ^6^. Perhaps even more importantly, quantifying the uncertainty of predictions (so called “pragmatic model performance”) poses a challenge in the presence of imputation ^11^: To what extent can we trust a predicted outcome if certain predictors were missing? Finally, many imputation strategies simply fail under extreme missingness, a scenario often neglected in the literature ^14^.

In this study, we will regard the following set-up consisting of two stages: i) In the parameter estimation stage, one set of researchers measures a complete (reference) dataset with *m* ∈ ℕ predictors and *n* ∈ ℕ observations captured in the *n* × (*m* + 1) design matrix *X*_ref_ (including the intercept) and outcome vector *y*_ref_ ∈ ℝ^*n*^. In this stage, parameter estimation for the LR model is performed. ii) In the prediction stage, another set of researchers has an (application) dataset with an incomplete predictor design matrix *X*_app_ and wants to predict the outcome *y*_app_ by applying the model of stage (i). Let us also assume that only summary statistics from the reference dataset *X*_ref_, *y*_ref_ are allowed to be passed from stage (i) to stage (ii) due to privacy constraints (which is typically the case when medical, genetic or otherwise identifiable patient data are involved). Here, we present a novel method called adaptive predictor-set linear model (aps-lm) that is able to transfer parameters from the reference (i) to the application (ii) stage in order to make predictions for incomplete observations without having to rely on imputation and without compromising privacy in the reference dataset. It is a CPU/storage-efficient analytical solution that does not introduce additional assumptions and was derived by using a predictor selection matrix operation, the Moore-Penrose pseudoinverse and the reduced QR-decomposition. aps-lm is a generalization of LR that can inherently handle missing values. It is fitted to the reference dataset of stage (i) and can be efficiently applied to the external dataset of stage (ii) suffering from missing data by taking into account the pattern of missing values for each individual prediction. We derived closed-form expressions for the coefficient estimates, standard errors, confidence and prediction intervals, and give statistical interpretation of the parameters for both unregularized and *l*^2^-regularized LR models. aps-lm has a very compact representation: The required parameters are an upper triangular matrix *R* ∈ ℝ^(*m*+1)×(*m*+1)^, the standard ordinary least squares (OLS) estimator 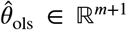 and the scalar *y*^⊤^*y* ∈ ℝ, which preserve the privacy of the reference dataset. To investigate the properties of aps-lm compared to its alternatives, we benchmarked it in a simulation study against two popular imputation methods: mean imputation and MICE multiple imputation ^15^. As a result, aps-lm showed greater prediction accuracy and reduced bias compared to the tested alternatives under a wide range of sample sizes, goodness-of-fit values, missing value types and covariance structures. Moreover, as a proof-of-principle application example, we explored the benefits of aps-lm in the context of epigenetic aging clocks; i.e., statistical models that can estimate a person’s biological age from epigenetic data, with promising prospects in the understanding of aging and assessment of clinical interventions ^16^ or other more specialized applications, such as in forensic genetics ^17^. Ultimately, we envision aps-lm as a simple and flexible method with a wide range of applications that can smooth the transition of linear models from the controlled research setting to the disarray of the real world, with special interest for applications under privacy restrictions such as medical data, data suffering from extreme missingness or requiring a thorough prediction error model.

The manuscript is organized as follows: The next section introduces the set-up, the problem of missing values and the corresponding notation. In section three, our novel aps-lm is developed. Simulations comparing the performance of aps-lm to mean and MICE imputation are given in section four. Finally, an application example is shown using data for epigenetic age prediction.

## 2 THE LINEAR MODEL AND THE PROBLEM OF MISSING DATA

### 2.1 Parameter estimation in standard linear regression

For *n* ∈ ℕ observations and *m* ∈ ℕ predictors with *n* > *m*, we define a standard linear model in the following form:

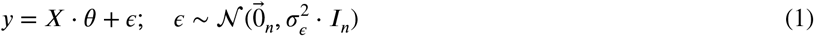

where *y* ∈ ℝ^*n*^ is the vector containing the dependent variable, *X* ∈ ℝ^*n*×(*m*+1)^ is a full rank design matrix with an intercept column such that rank(*X*) = *m* + 1, θ ∈ ℝ^(*m*+1)^ is the vector of linear coefficients, *ϵ* ∈ ℝ^*n*^ is the random error vector, 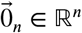 is the vector whose components are all equal to zero, 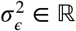 of order *n*. The parameter vector θ is usually estimated by the OLS estimate 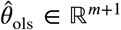 design matrix *X* as which can be written for a full-rank design matrix *X* as

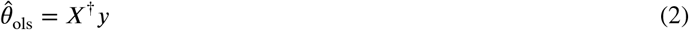

where †denotes the Moore-Penrose pseudoinverse, easily computed from the singular-value decomposition ^18^. We use this expression in our derivation of the aps-lm model (at equation 6) due to its compactness rather than the more popular normal equations 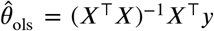, where *X*^⊤^ denotes the transpose of *X*. Note that these two representations are equivalent for full-rank design matrices since then *X*^†^= (*X*^⊤^*X*)^−1^*X*^⊤^.

### 2.2 Set-up and notation for our problem

We will formulate our problem in two stages.

i. In the first stage, we have the well-known parameter estimation set-up of a linear model where a reference dataset with outcome 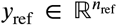 and design matrix 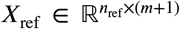 is available and the aim is to estimate the parameter vector *θ* ∈ ℝ^(*m* + 1)^ of the corresponding linear model. We assume no missing values in *X*_ref_ and *y*_ref_ in stage (i).
ii. In the second stage, we have the prediction set-up where an application dataset with design matrix 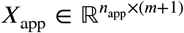 is measured and the outcome 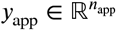 is to be estimated (sometimes the term “predicted” is also used).

Here, *n*_ref_ and *n*_app_ are the number of observations in the reference and application dataset, respectively. We will further assume that the research groups of stages (i) and (ii) are independent from each other and that *X*_ref_ and *X*_app_ are hosted on different servers with access only for the respective group. Only summary statistics can be freely exchanged between research groups as the data are considered sensitive and are hence to be protected because of data privacy ^19^. The objective in stage (ii) is to make predictions of the outcome *y*_app_ from *X*_app_ by employing a linear model without violating privacy restrictions of the individuals registered in the reference dataset. In the absence of missing values in *X*_app_ the problem is trivial and can be solved for example by computing 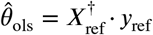 in stage (i), transferring this privacy-preserving summary statistic between servers and computing the linear predictions as

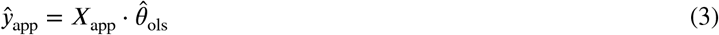

However, the above scheme fails if missing values are present in stage (ii) in *X*_app_, as the direct computation of 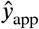 is not possible in presence of missing values. To tackle this issue, one possibility is to pre-train linear models for all combination of available variables to get missing-value dependent estimates of θ in stage (i). The downside is that the number of models and parameters to be stored scales with *m* in an unfavorable fashion ^7^:

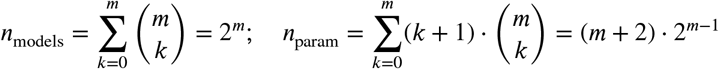

Another approach is to train upon demand: For each observation in the application dataset in stage (ii) note the missing values and exclude the corresponding variables in the reference dataset of stage (i). Then estimate linear coefficients in stage (i), pass the estimates over to stage (ii) and generate predictions using these sample-customized linear coefficients in stage (ii) ^6^. This procedure, however, requires intricate multi-party computation and is thus usually highly impractical. As a result, the most convenient option is to simply perform imputation on *X*_app_ and employ the full model as in 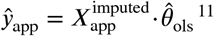 ^11^. Imputation has the drawback that it causes high computational costs, especially for multiple imputation, and introduces assumptions, typically MCAR or MAR, which in practice are often violated. To overcome the aforementioned limitations, we propose aps-lm, an extension to linear models which can inherently handle any combinations of available predictors.

### 2.3 A matrix operation to exclude rows/columns

In aps-lm it is necessary to select subsets of predictors. To transfer this to the language of linear algebra, we introduce a matrix operation. Let *A* ∈ ℝ^*n*×*m*^be a matrix from which to delete certain rows or columns. Let Ω := {*a*_1_, *a*_2_, …, *a*_*r*_} ⊂ {1, …, *n*} be a set of indices of *r* rows to remove, where *r* ∈ {1, …, *n*}. Similarly, Ω can be defined to exclude columns. We denote by 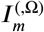 and 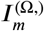 the identity matrix of order *m* excluding column or row *k* iff *k* ∈ Ω, respectively. We can extend the row/column removal operation to the matrix *A* with the following identities:

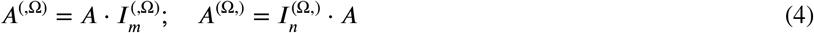

where *A*^(,Ω)^∈ ℝ^*n*×(*m*−|Ω|)^, *A*^(Ω,)^∈ ℝ^(*n*−|Ω|)×*m*^and |Ω| denotes the cardinality of the set Ω. For example:

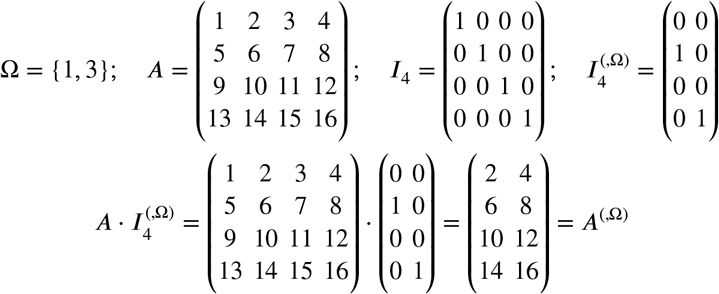

Note that this operation projects an *n* × *m* matrix down onto an *n* × (*m* − |Ω|) or (*n* − |Ω|) × *m* matrix. The operation and its properties are formally defined in Suppl. Methods 1.

## 3 THE ADAPTIVE PREDICTOR-SET LINEAR MODEL (APS-LM)

### 3.1 Definition of aps-lm and parameter estimation

Let Ω be a set of missing variables such that Ω ⊂ {2, …, (*m*+1)} (the first column of *X* is reserved for the intercept). We can now describe the linear model in the presence of missing data in the following central equation for our novel adaptive predictor-set linear model (aps-lm) by combining equations 1 and 4:

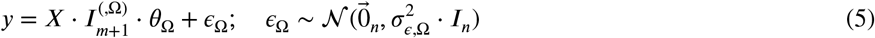

where θ_Ω_ ∈ ℝ^(*m*+1−|Ω|)^is the vector of remaining linear regression coefficients, *ϵ*_Ω_ ∈ ℝ^*n*^is the vector of error terms and 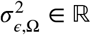 is the variance of the error terms. Note that the new design matrix 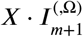 is the original design matrix *X* but having removed the columns corresponding to the set Ω. The model of equation 5 can be seen as a projection of the full saturated model of equation 1 to the submodel on the set of available predictors. For a later application in stage (ii), Ω will represent the missing variables for an individual as recorded in *X*_app_. Thus, every set Ω corresponds to a different standard OLS linear submodel, with number of coefficients equal to *m* + 1 − |Ω| and different 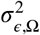. We will first derive an estimate for θ_Ω_. For this, we will assume full rank and apply the normal equation in its pseudoinverse form (equation 2):

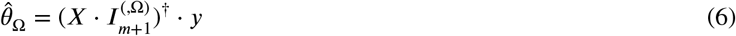

This expression, however, depends on *X* and is thus neither privacy- nor storage-friendly. To solve this issue, we exploit some properties of the pseudoinverse. It has been shown that (*A* ⋅ *B*)^†^= *B*^†^⋅ *A*^†^if any of the following properties applies: i) A has orthonormal columns, *A*^⊤^⋅ *A* = *I*, ii) B has orthonormal rows, *B* ⋅ *B*^⊤^= *I*, iii) A and B have linearly independent rows and columns, respectively, iv) *B* = *A*^⊤^or v) *B* = *A*^†20^. Though none of these conditions necessarily apply for *X* or 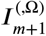, we can achieve condition i) by performing reduced-QR decomposition of *X*. We can write *X* as

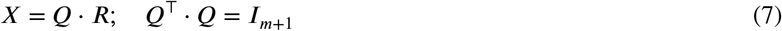

where *Q* ∈ ℝ^*n*×(*m*+1)^has the same dimension as *X*, and *R* ∈ ℝ^(*m*+1)×(*m*+1)^is an upper triangular matrix. This differs from the more popular full QR decomposition in which *Q* ∈ ℝ^*n*×*n*^and *R* ∈ ℝ^*n*×(*m*+1)^. Reduced-QR decomposition was initially developed in numerical linear algebra to improve implementation efficiency, but here we use it to compress the number of parameters stored in *R*. Introducing equation 7 into equation 6, we get:

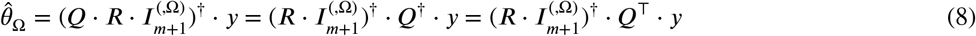

In the last step, we also used the result that the pseudoinverse of a matrix with orthonormal columns is its transpose. To obtain the final expression for 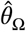, we want to exchange *Q*^⊤^⋅*y* for a more interpretable term. We can do so by setting Ω to the empty set *∅*:

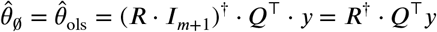

By multiplying by *R* from the left on both sides we get:

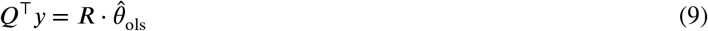

Lastly, by substituting equation 9 into equation 8, we arrive at the final expression for aps-lm coefficient estimates as a function of Ω, *R* and 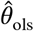 :

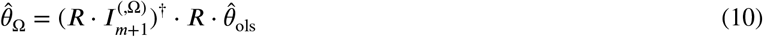

The matrix 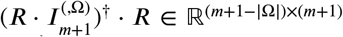 transforms 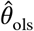, which corresponds to Ω = *∅*, to 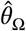 for any set Ω *≠ ∅*. 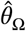 thus corresponds to 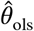 mapped down to the reduced predictor space. The model parameters *R* and 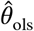 do not depend on Ω. We are here in stage (i) where *X* corresponds to *X*_ref_ and *n* to *n*_ref_. *R* and 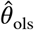 can be readily calculated and transferred to stage (ii). As *R* ∈ ℝ^(*m*+1)×(*m*+1)^is upper triangular and 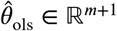, the total number of parameters stored in *R* and 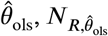, is independent of *n* and equal to:

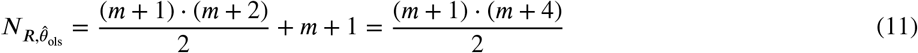

This set-up is satisfactory if *n* ≫ *m*, which is the typical case in linear regression analysis. In addition, neither *X* nor *y* can be retrieved from *R* and 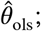 as a result, privacy of the participants in the reference dataset cannot be compromised. With respect to CPU efficiency, although aps-lm is better represented in its pseudoinverse form (equation 10), the computation of the pseudoinverse has an unfavorable algorithmic time complexity (e.g., 𝒪((*m* + 1 − Ω)^2^ ⋅ *n*) with the Golub and Kahan algorithm ^21^) ^22^. For the practical implementation of aps-lm, we can substitute the pseudoinverse with the more efficient sweep operator ^23,10^ (more details in Suppl. Methods 5).

### 3.2 Making predictions with 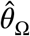 on *X*_app_

Let us now set *X* := *X*_ref_. In stage (i), *R* and 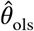 are calculated and can be freely transferred to stage (ii). In stage (ii), for each observation *k* ∈ {1, …, *n*_app_} the set Ω_*k*_ of missing values is determined and a personalized set of linear parameters, 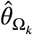, can be estimated by equation 10. An individual prediction is then simply computed by:

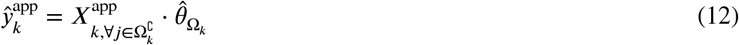

where 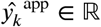 is the prediction for individual *k* and 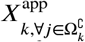 contains all non-missing variables of individual *k*. This step can be trivially parallellized for improved computational performance.

### 3.3 Parameter interpretation

From the parameter matrix *R* it is possible to retrieve the sample size *n* ∈ ℕ, mean vector 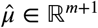 and the covariance matrix 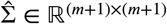, defined as

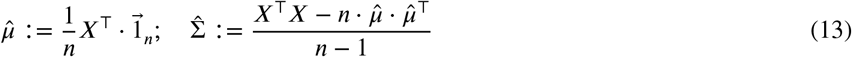

where 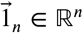 denotes the vector whose components are all equal to 1. 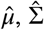 and *n* can be easily recovered from the entries in the matrix *R* (derivation in Suppl. Methods 2.1):

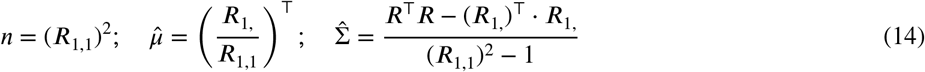

The relationship is bijective, meaning that *R* can also be computed from these same parameters:

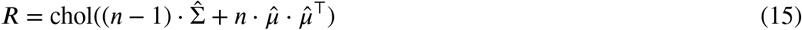

where for a matrix *A* the notation chol(*A*) stands for the matrix *B* from the Cholesky decomposition with *B*^⊤^⋅ *B* = *A*.

### 3.4 aps-lm standard errors

We can obtain the standard errors for the coefficients in aps-lm following somewhat similar steps as in the derivation of the standard errors in the classical linear model (available in Suppl. Methods 3):

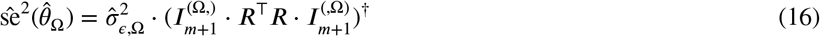

where

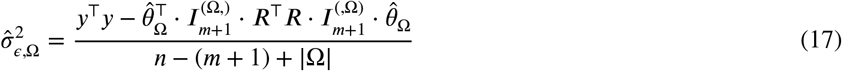

Here, we see that one new parameter is introduced, *y*^⊤^*y* ∈ ℝ. Including *y*^⊤^*y* in our set of aps-lm parameters, we end up with a total of (*m* + 1)(*m* + 4)/2 + 1 parameters in order to compute linear coefficient estimates and standard errors for all missing value sets Ω (see equation 11). Standard errors can be used to compute confidence intervals for the mean response as in:

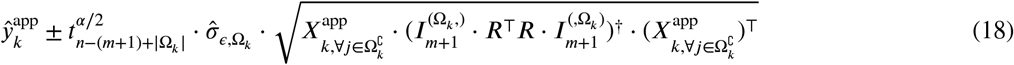

where 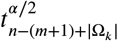 is the α/2-quantile of the *t*-distribution with *n* − (*m* + 1) + |Ω_*k*_| degrees of freedom. We can also obtain prediction intervals as in:

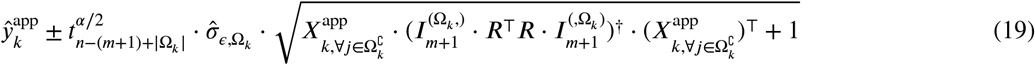

This expression allows the reporting of prediction errors that take Ω_*k*_ into account without the need to access the reference dataset.

### 3.5 Adaptive predictor-set ridge regression

Regularization is a statistical methodology that modifies objective functions to include certain constraints, especially useful to solve ill-posed optimization problems. Ridge regression (aka *l*^2^-regression or Tikhonov regression) is a regularized version of linear regression in which an *l*^2^-penalty for the linear coefficients is introduced to the OLS linear regression objective function. Expressing the objective function in its Lagrangian form, we have:

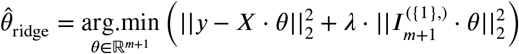

where λ is the regularization parameter (equivalent to a Lagrange multiplier of the *l*^2^-norm constraint of *θ*) and 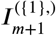 is included in the regularization term to avoid penalizing the intercept variable. Ridge regression manages to reduce the standard errors of 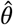, but at the expense of creating bias in the linear coefficients estimates. Conveniently, a closed-form solution is available for this problem ^24^. Thus, we can also derive an adaptive predictor-set ridge regression model aps-ridge. The calculations are similar to aps-lm and given in Suppl. Methods 4. The model parameters, which are derived in stage 1 and are communicated to stage 2, are now *R*_λ_, 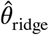, *y*^⊤^*y* and λ. The ridge linear coefficients estimate excluding the set of variables in Ω is given by:

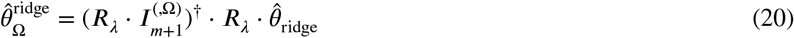

where 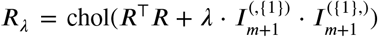 and 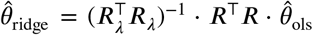. This expression is parallel to that of aps-lm (equation 10), but substituting *R* and 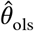 by *R*_λ_ and 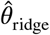, respectively. Alternatively, 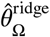 can be efficiently calculated with the sweep operator (Suppl. Methods 5). We also derived standard errors for aps-ridge (Suppl. Methods 4):

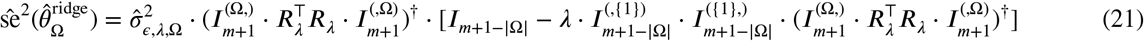

where

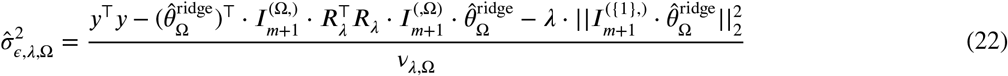

and where *v*_λ,Ω_ denotes the effective degrees of freedom ^25^, computed as

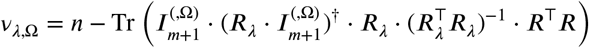

with Tr being the trace operator. Similar to aps-lm, we approximate confidence intervals for the mean response by

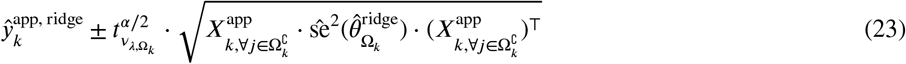

and prediction intervals by

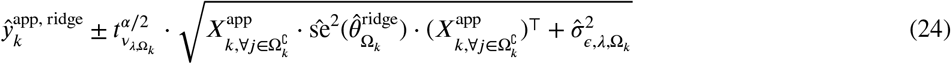

This is only an approximation since the distribution of the ridge estimator is unknown ^26^. Warnings against the use of confidence intervals in penalized regression are common due to the fact that the estimation of the linear coefficients is biased ^24^ (and thus of the aps-ridge coefficients). However, this does not mean that the predictions are biased. In fact, 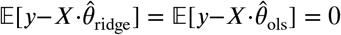 (Suppl. Methods 4.1); therefore, our reported confidence and prediction intervals for the response are unbiased.

### 3.6 An approximation to adaptive predictor-set generalized linear models

In the study of Marshall *et al* ^10^, a similar approach to aps-lm is employed in the context of logistic regression. This is only one possibility of extending the aps-lm to generalized linear models. Other possibilities and related properties have to be investigated in future studies. We will give here only a rough description of their method and compare it to aps-lm. Written in aps-lm notation, the following generalized linear model is proposed:

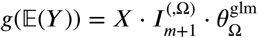

where *g* is a link function. Although *g* is originally defined on ℝ, we can extend this easily to ℝ^*n*^by applying *g* to each component (in order to conform to our matrix notation). In ^10^ they particularly focus on the logistic regression. Let 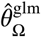 and 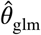 be the maximum likelihood estimators of 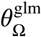 and 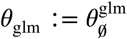. We then have:

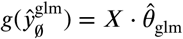

and

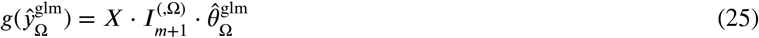

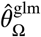 is computationally intensive and difficult to calculate. Therefore, Marshall *et al* ^10^ propose an estimator which is equivalent to the following approximation:

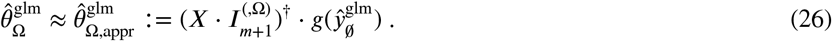

As can be seen from equation 25, the prediction based on the missing dataset, 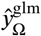, is replaced by the prediction based on the whole dataset, 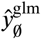. In our notation, 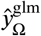 corresponds to the desired stage two prediction whereas 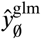 refers to the stage 1 prediction of the full model.

To compare with aps-lm, let us set the link function to the identity function:

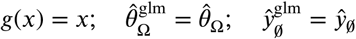

where 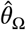 is the aps-lm estimate of the linear coefficient vector. Curiously, in this scenario, the approximation of ^10^ becomes an equality. Employing equation 7 we get:

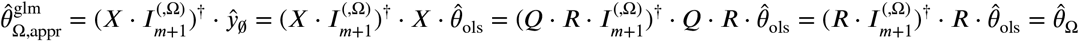

Thus, with identity link function, Marshall’s approximation is equal to aps-lm’s 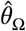. For a general link function *g*, note that equation 26 needs access to the full stage 1 data. We can also extend Marshall’s approximation in a way that protects the privacy of the individuals in the reference dataset by expanding with equation 7:

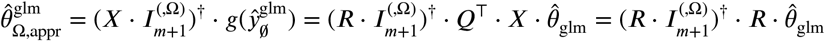

This last expression is equivalent to the standard aps-lm coefficient estimate expression (equation 10) but replacing 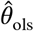 by 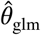. However, this approximation will worsen with increasing |Ω|. More research in this area is clearly necessary.

## 4 SIMULATION STUDY

To test the benefits of aps-lm over its alternatives, we performed an extensive simulation study, which was set-up according to the ADEMP guidelines ^27^ covering aims, data generation mechanism, simulation estimands, methods and performance measures. Four rounds of simulations were performed with different aims (Table 1): In rounds 1 and 2, we compared our novel aps-lm to (unconditinal) mean imputation and to MICE multiple imputation ^15^. For this, we first simulated the reference and application datasets *X*_ref_, *y*_ref_, *X*_app_ and *y*_app_ of stage (i) and (ii) assuming that they follow the same linear model and without missing values. To create missing values for stage (ii), we removed entries from *X*_app_. Then we supposed that the true simulated outcome *y* = *y*_app_ of stage (ii) is unknown and we estimated it with three methods: (unconditional) mean imputation, MICE multiple imputation and aps-lm. For the two imputation methods, the estimated linear model from stage (i) was applied to the imputed dataset *X*_app,imputed_ of stage (ii). For aps-lm, the parameters *R* and 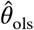 were derived in stage (i) from *X*_ref_ and *y*_ref_ and applied to the unimputed *X*_app_ in stage (ii). Application of the estimated linear model from stage (i) to the full application dataset without missing values served as a benchmark and gold standard. In round 1 a wide range of conditions were simulated (21 simulations) and in round 2 the influence of missing value type was examined (6 simulations). Round 3 compared the performance of aps-ridge to aps-lm and finally in round 4 the simulated coverage of the confidence and prediction intervals was compared to the predefined coverage. To judge the performance of the approaches in rounds 1 to 3, we compared the estimated 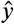 with the true simulated value *y*. For this, we applied four measures: the Pearson correlation squared 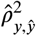 (for convenience since meaningful negative correlations are not expected) (rounds 1-2), the bias between *y* and 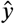 (rounds 2-3), the concordance correlation (round 2) and the mean squared prediction error (MSE) (rounds 2-3). For the ridge models, the regularization parameter λ had to be determined by cross-validation which was based on the MSE in the software we applied. Therefore, using the Pearson correlation in the comparison of ridge to other models would be detrimental to those models. We consequently used only the MSE in round Details on the implementation are available in Suppl. Methods 6.

**Table 1.** Simulation study overview.

| Round | Simulation | Characteristics | Missing value type |
| --- | --- | --- | --- |
| 1 | 1 | Independence | MCAR |
| 1 | 2 | Weak dependence | MCAR |
| 1 | 3 | Strong dependence | MCAR |
| 1 | 4 | Independence | MAR-L |
| 1 | 5 | Weak dependence | MAR-L |
| 1 | 6 | Strong dependence | MAR-L |
| 1 | 7 | Independence | MAR-LR |
| 1 | 8 | Weak dependence | MAR-LR |
| 1 | 9 | Strong dependence | MAR-LR |
| 1 | 10 | Independence | MAR-C |
| 1 | 11 | Weak dependence | MAR-C |
| 1 | 12 | Strong dependence | MAR-C |
| 1 | 13 | Independence | MNAR-L |
| 1 | 14 | Weak dependence | MNAR-L |
| 1 | 15 | Strong dependence | MNAR-L |
| 1 | 16 | Independence | MNAR-LR |
| 1 | 17 | Weak dependence | MNAR-LR |
| 1 | 18 | Strong dependence | MNAR-LR |
| 1 | 19 | Independence | MNAR-C |
| 1 | 20 | Weak dependence | MNAR-C |
| 1 | 21 | Strong dependence | MNAR-C |
| 2 | 1 | Independence, $\theta_1$ | All types |
| 2 | 2 | Weak dependence, $\theta_1$ | All types |
| 2 | 3 | Strong dependence, $\theta_1$ | All types |
| 2 | 4 | Ultra-strong dependence, $\theta_1$ | All types |
| 2 | 5 | Independence, $\theta_2$ | All types |
| 2 | 6 | Weak dependence, $\theta_2$ | All types |
| 2 | 7 | Strong dependence, $\theta_2$ | All types |
| 3 | 1 | Independence, $\theta_1$ | All types |
| 3 | 2 | Weak dependence, $\theta_1$ | All types |
| 3 | 3 | Strong dependence, $\theta_1$ | All types |
| 4 | 1 | Independence, $\theta_1$ | All types |
| 4 | 2 | Weak dependence, $\theta_1$ | All types |
| 4 | 3 | Strong dependence, $\theta_1$ | All types |

Each round pursued different aims: (round 1) comparing aps-lm to mean and MICE imputation under a wide range of conditions; (round 2) comparing aps-lm to mean and MICE imputation simultaneously assessing all types of missing values; (round 3) benchmarking aps-lm against aps-ridge; (round 4) testing the coverage of confidence and prediction intervals. Three types of covariance matrices of predictors are used with either no dependence structure (independence), weak or strong dependence. In round 2, additionally a covariance matrix with ultra-strong dependence was simulated. $\theta_1$ and $\theta_2$ are two different linear coefficient vectors to assess model-dependent effects. MCAR: missing completely at random; MAR: missing at random; MAR-L, -R, -LR, -C: MAR with an *L*, *R*, *LR* or *C* link function that makes the left, right, both tails or the center of the distribution more likely to suffer from missing values (for details see Suppl. Methods 6); MNAR: missing not at random

Briefly, we first simulated *X*_ref_ and *X*_app_ from a multivariate normal distribution by fixing mean vector *μ* and covariance matrix Σ for *m* = 10 predictors and appending an intercept column. Secondly, to generate *y*_ref_ and *y*_app_, we fixed the parameter vector θ, computed *X*_ref_ ⋅ θ and *X*_app_ ⋅ θ and added white noise (*ϵ*) whose variance 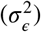 was tuned to adjust the goodness-of-fit (GOF) of the linear model. Three types of covariance matrices of predictors were used: independent, weakly dependent and strongly dependent; their corresponding correlation matrices are visualized in Figure S1A-C. For round 2, an additional a covariance matrix with ultra-strong dependence was investigated (Figure S1D). Three types of missing value schemes were employed ^28^. We provide extensive definitions and descriptions on how the missing values types were implemented in this study in Suppl. Methods 6.1. Briefly, missing completely at random (MCAR) occurs when the probability of becoming missing does not depend on observed or unobserved variables but solely on a single parameter, the proportion of missing values in *X*_app_, here denoted by *p*_NA_, which we set equal for all variables in our simulations. For missing at random (MAR) the probability of becoming missing does not depend on the variable in question but on other observed variables and an additional tuning parameter. For our simulations, this relationship is modelled with the use of link functions; four types of link functions are used in this study, all of which are based on the logistic function, such that a higher probability of becoming missing is given to extreme points to the left (L), to the right (R), both sides (LR) or values closer to the center of the distribution (C). Finally, the term missing not-at-random (MNAR) is used in all other cases, i.e., when the probability of a variable becoming missing depends additionally on the value of the variable itself or on unobserved variables. We model this relationship again with the aforementioned link functions.

### 4.1 Round 1: aps-lm under a wide range of conditions

In this round, we aimed to explore a wide simulation parameter space comprising the proportion of missing values (*p*_NA_), linear model GOF (*R*^2^), sample size (*n*), missing value type and covariance structure of predictors. We focused on the squared Pearson correlation measure in this section for convenience. The results of all 21 simulations are available in Figures 1, 2 and 3 and Figures S2 to S19. By definition, aps-lm tends towards unconditional mean imputation when a diagonal covariance matrix is used. In general, this is apparent in the simulations (Figures S2, S4, S7, S9, S12, S14 and S17); however, at low *n* and high *p*_NA_ or under MNAR, aps-lm outperforms unconditional mean imputation in terms of mean squared error in spite of independence. The cause is probably that mean vector estimation is less reliable on the relatively small application dataset when the sample size is low or many missing values occur, whilst aps-lm has access to the robust estimates from the reference dataset without missing values (stored in the *R* matrix). When dependence is included in the covariance matrix, aps-lm systematically outperforms both unconditional mean and MICE imputation in the explored parameter space (1, 2 and 3). In general, despite the much higher computational cost of MICE, this method performs worse than aps-lm or even unconditional mean imputation. Only at high sample size, low *p*_NA_ and some degree of dependence, MICE performs slightly better than unconditional mean imputation but never better than aps-lm.

**Figure 1.**
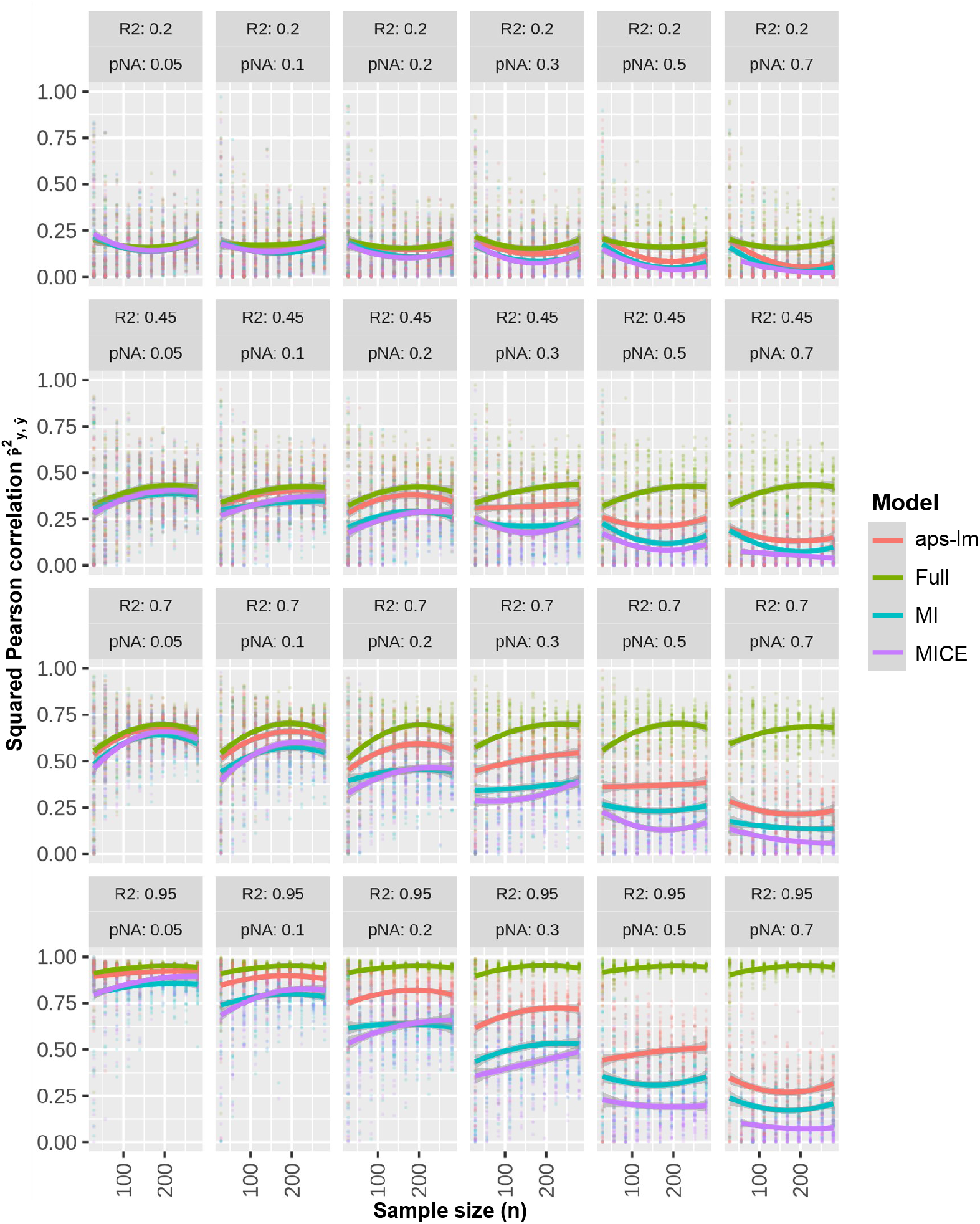
Round 1-Simulation 2: MCAR under weak dependence. Missing values were injected on the application dataset following an MCAR scheme with variable proportion of missing values (*p*_NA_). Squared Pearson correlation between real and predicted dependent variable is plotted at varying sample size (*n*), *R*^2^and *p*_NA_ for: i) the estimated linear regression model of stage (i) applied on the application dataset *X*_app_ without missing data (full), applied on *X*_app_ with missing data after performing ii) un-conditional mean (MI) or iii) MICE imputation (MICE) and iv) aps-lm without applying any imputation. Simulation results are visualized as jitter plots while overall tendencies in the mean are represented as LOESS curves.

**Figure 2.**
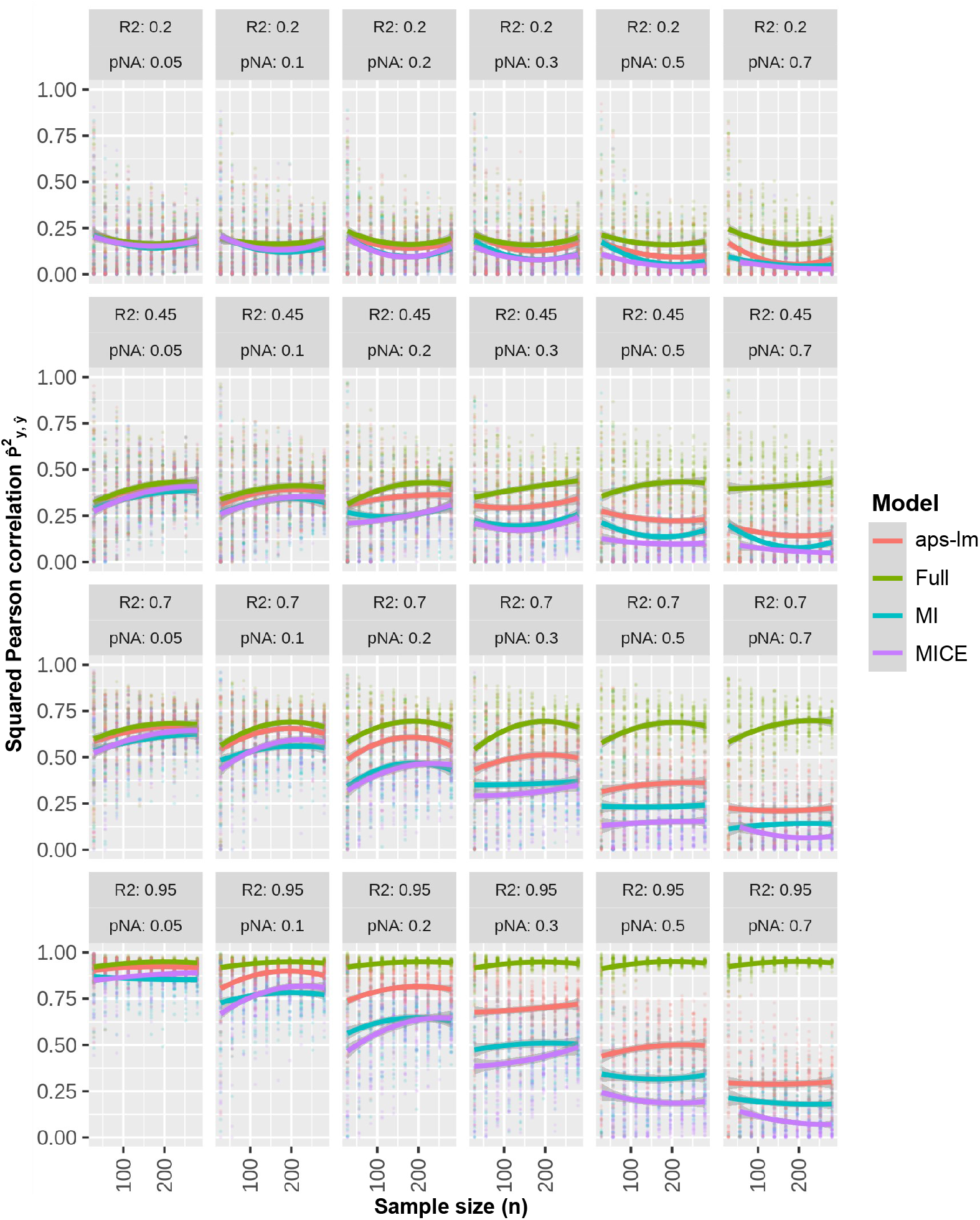
Round 1-Simulation 8: MAR-LR under weak dependence. Missing values were injected on the application dataset following a MAR-LR scheme that makes missing values more likely on both the left (L) and the right tails (R). For further details see legend to Fig. 1.

**Figure 3.**
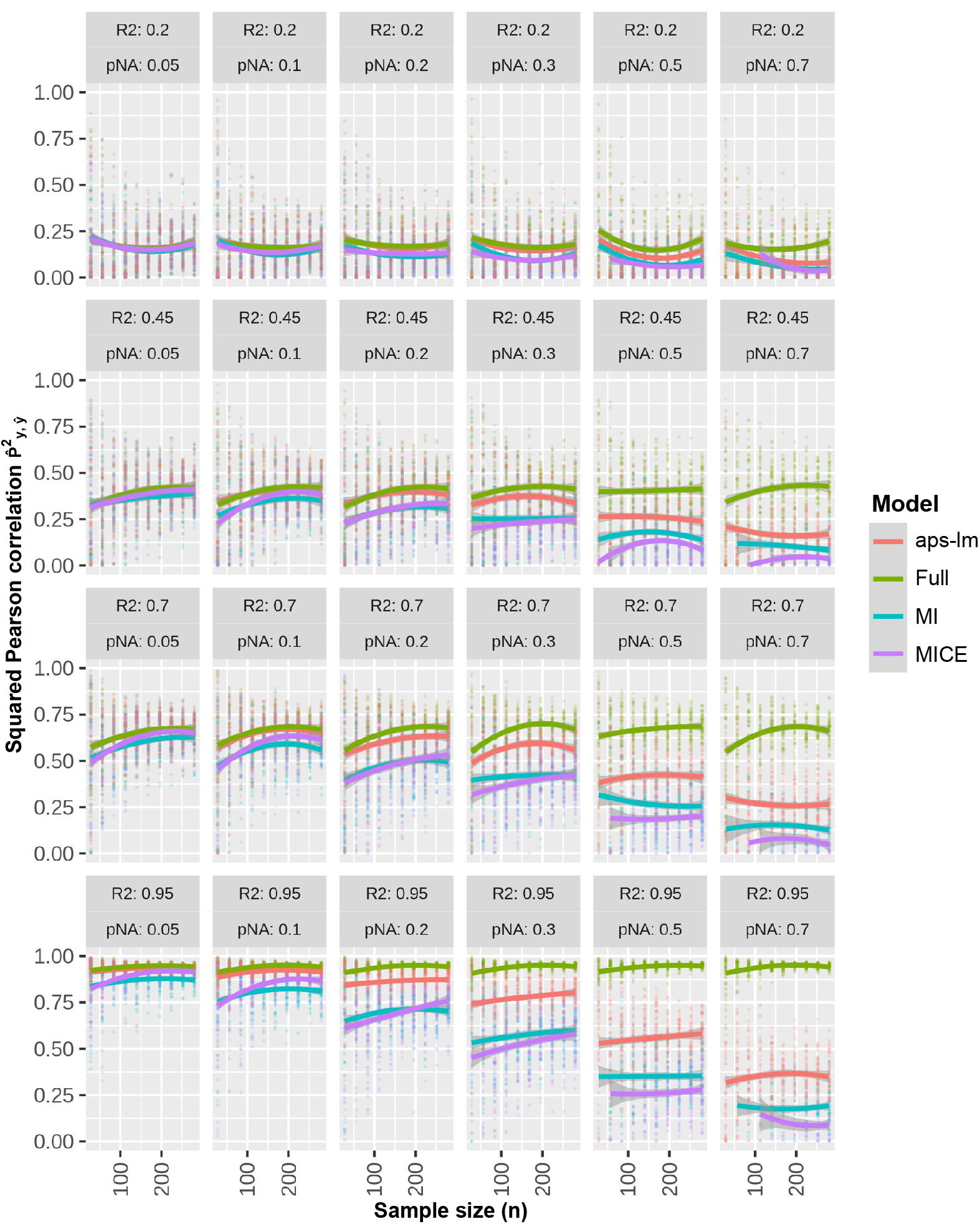
Round 1-Simulation 14: MNAR-L under weak dependence. Missing values were injected on the application dataset following a MNAR-L scheme that makes missing values more likely on the left tail (L). For further details see legend to Fig. 1.

### 4.2 Round 2: The influence of different types of missing values

In this round, to draw comparisons between different missing value schemes, we tested all missing value schemes in parallel on the same simulated datasets instead of injecting missing values on independent datasets, (Figure 4, Figures S20-21). Additionally, we modified θ to see whether any of the observed tendencies were θ-dependent (Figures S22-24). In general, MNAR is more detrimental to performance than MCAR or MAR. In particular, the decrease of 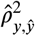 with higher *p*_NA_ values is more pronounced for MNAR and considerable bias is generated, the direction of which is θ- and missing value subtype-dependent. Though MICE and mean imputation seem to be very sensitive to MNAR-induced bias, aps-lm manages to quench its influence, especially in the presence of dependence. In this round we also computed the MSE and the concordance correlation in addition to the Pearson correlation and the bias. These two measures showed very similar results (Figures S25-27). Additionally, we simulated a covariance matrix with ultra-strong dependence (Figure S1D). The differnces between the methods is even more pronounced in this case (Figure S28). This round further highlights the potential of aps-lm to tolerate missing values, even under the MNAR mechanism.

**Figure 4.**
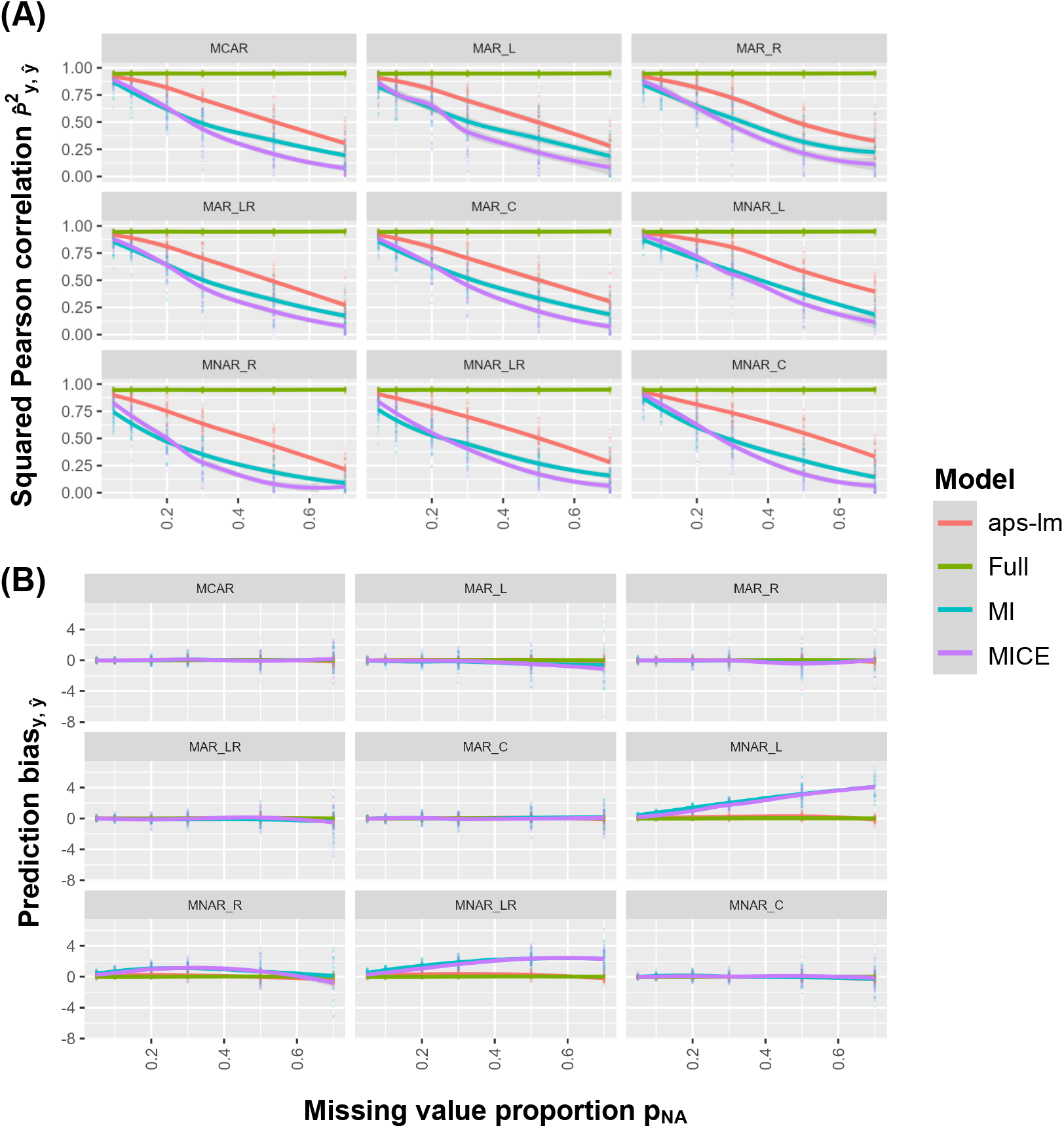
Round 2-Simulation 2: all missing value types under weak dependence. Missing values were injected on the application dataset following all implemented missing value schemes (MCAR, MAR-L,R, LR, C and MNAR-L, R, LR, C). (A) Squared Pearson correlation between real and predicted dependent variable or (B) prediction bias is plotted at varying missing value type, *R*^2^and *p*_NA_. For further details see legend to Fig. 1.

### 4.3 Round 3: aps-lm vs aps-ridge

With the simulations in round 3 we aimed to compare the performance of ordinary linear regression versus ridge regression for our adapted predictor-set methods (Figure 5, Figures S29-30). Similarly to round 2, we simulated all missing value scenarios on the same datasets. Because ridge regression is used here, we used MSE instead of the Pearson correlation as performance measure in addition to the bias. For our simulations we see very similar performance in bias and MSE for the full ordinary linear and ridge models as well as for aps-ridge and aps-lm. Note, however, that we only used ten predictors. Ridge is expected to perform better for higher numbers of variables. An overlay of the results of round 2 and round 3 can be found in Figures S31-33. Here again one can see that the full models perform best while aps-lm and aps-ridge give similar results which are superior to mean and MICE imputation.

**Figure 5.**
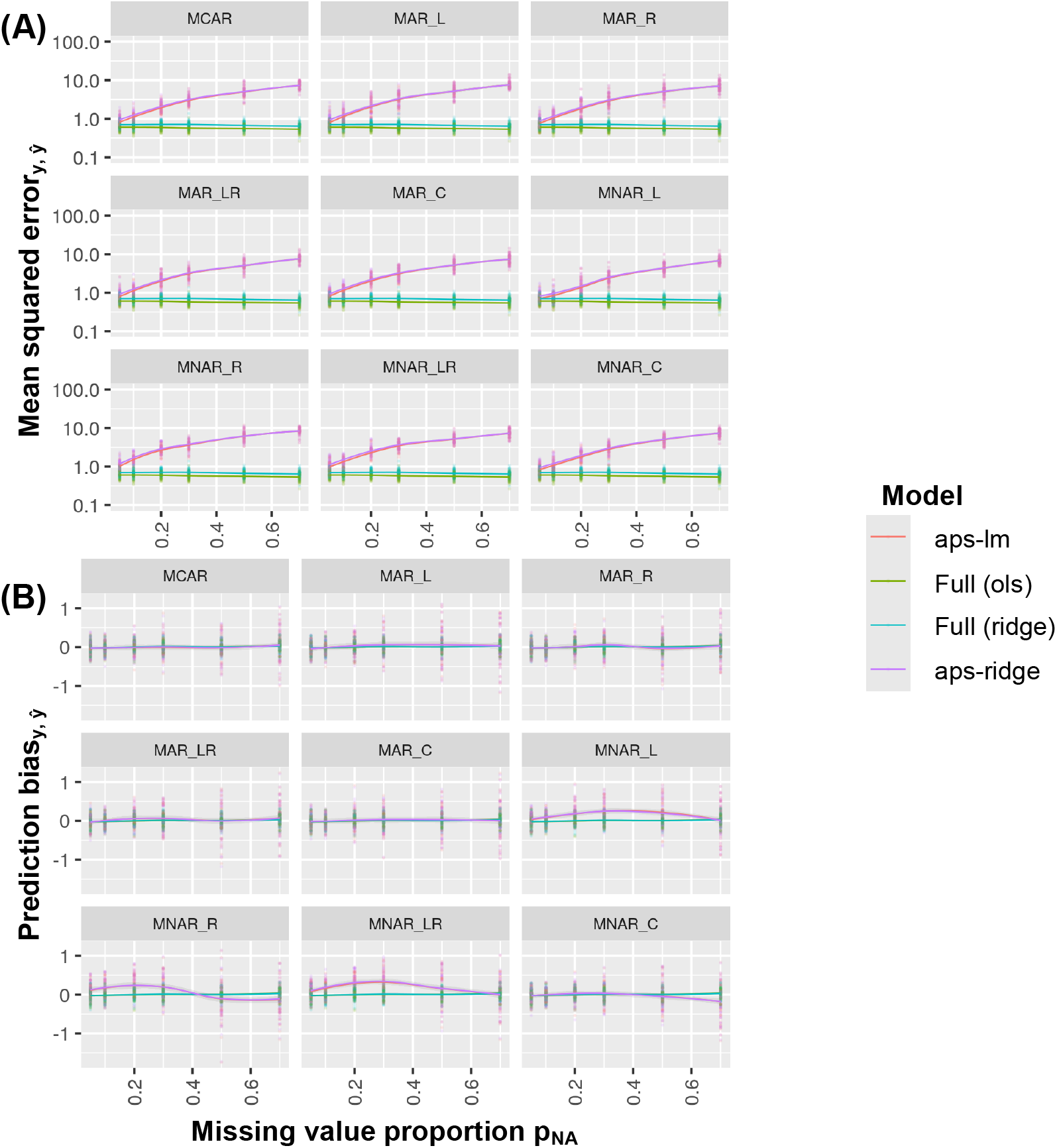
Round 3-Simulation 2: all missing value types under weak dependence employing θ_1_. Missing values were injected on the application dataset following all implemented missing value schemes (MCAR, MAR-L,R, LR, C and MNAR-L, R, LR, C). (A) Prediction mean squared error between real and predicted dependent variable or (B) prediction bias is plotted at varying missing value type, *R*^2^and *p*_NA_. This is shown for the estimated ordinary least squares and ridge linear regression model of stage 1 applied on the application dataset *X*_app_ without missing data (i) full (ols), ii) full (ridge), respectively) and iii) aps-lm and iv) aps-ridge without applying any imputation. The regularization parameter for full (ridge) and aps-ridge was set equally and obtained via cross-validation on the reference dataset based on the mean squared prediction error metric. Note that the mean squared error axis is in log-scale. Curves aps-ridge and aps-lm overlay in panels A and B; curves full (ols) and full (ridge) overlay in panel B.

### 4.4 Round 4: Coverage of confidence and prediction intervals

In this round it was investigated whether the predefined coverage of the confidence and prediction intervals was achieved for our aps-lm model. We compared the coverage for a given 95 % coverage level. The simulation procedure needed to be adapted because now simulations had to be made from the submodel 5. A detailed description can be found in Suppl. Methods 6.6. The simulation was performed for four different sets of missing values Ω (with 0, 1, 3 and 6 missing predictors out of 10), three linear model GOF (*R*^2^=0.25, 0.50, 0.95) and three sample sizes (*n*=50, 100, 200). Figure 6 and Figures S34-35 show the results of the simulations. The predefined coverage is nicely reached for all the different scenarios. In addition to the GOF of the full model *R*^2^, the GOF of the submodel 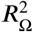 is also displayed. For no missing values, i.e., Ω_0_, those two of course agree. For Ω_1_ with only one missing predictor, the value of *R*^2^is only slightly reduced. In contrast, one can see massively lower values of 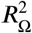 compared to *R*^2^for Ω_2_ and Ω_3_. Note that 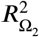 is smaller than 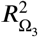 even though Ω_2_ contains three missing predictors and Ω_3_ six. This is due to the fact that predictor seven is included in Ω_2_ but not in Ω_3_. Predictor seven is the most relevant influence variable in our simulated model because it has by far the largest absolute regression coefficient (compare equation (45) in Suppl. Methods).

**Figure 6.**
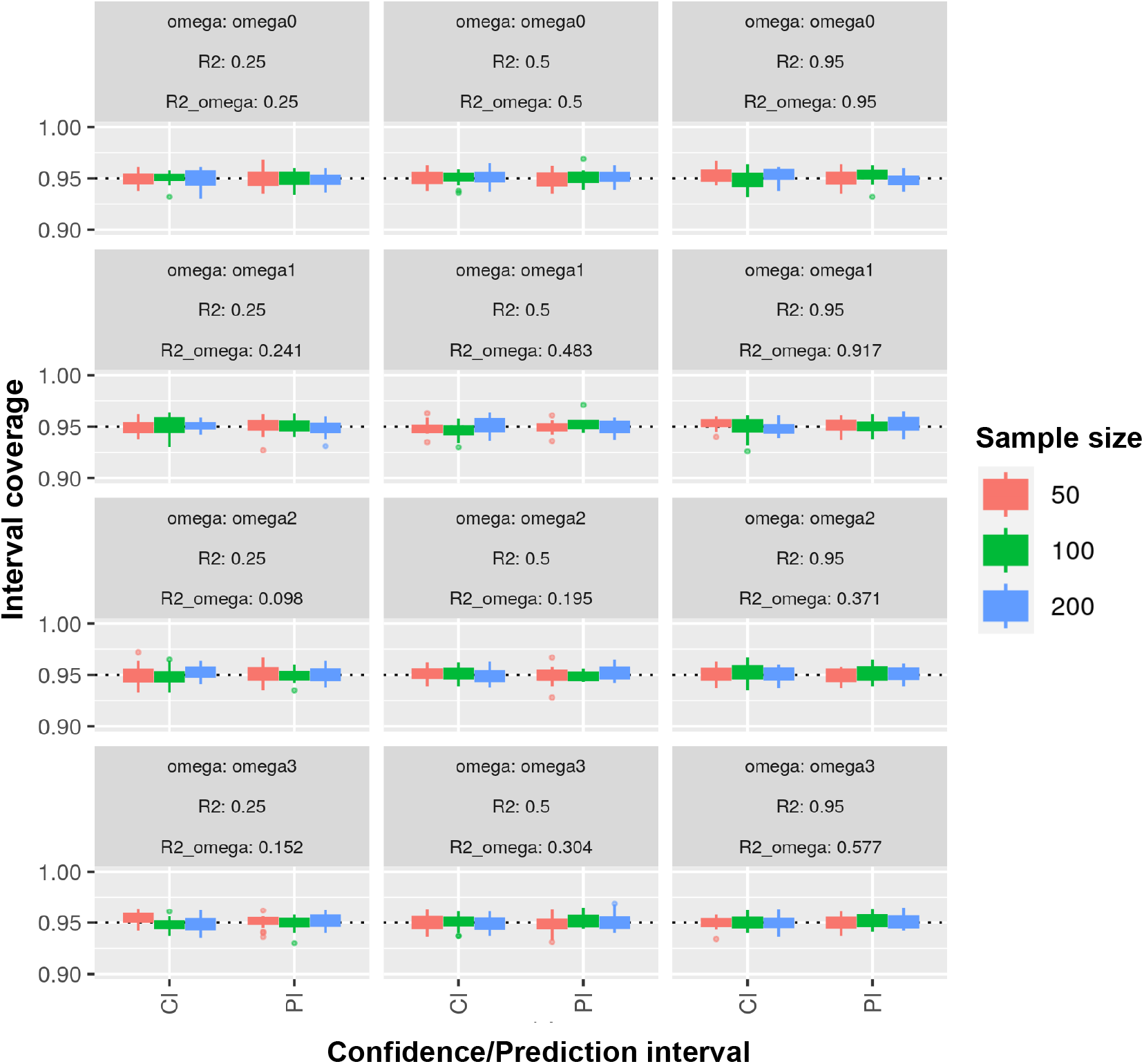
Round 4-Simulation 2: aps-lm coverage confidence and prediction intervals under weak dependence for different *R*^2^and sample sizes *n*. Four sets of missing value sets are tested: Ω_0_ = *∅*(omega0), Ω_1_ = {2} (omega1), Ω_2_ = {2, 4, 7} (omega2) and Ω_3_ = {1, 2, 3, 4, 8, 9} (omega3). 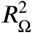 (R2_omega) denotes the corresponding *R*^2^of the submodel after excluding the predictors from in Ω. Boxplots represent the distribution of the interval coverage across replicates (*N*_rep_ = 20), estimated as the number of iterations at which 𝔼[*Y X*] (confidence interval) or *Y*| *X* (prediction interval) lies within the bounds of the estimated intervals; the total number of iterations employed for each of the 20 replicates was *N*_cov_ = 1000. Note that the y-axis begins at 0.90. CI: confidence interval; PI: prediction interval.

## 5 APPLICATION TO EPIGENETIC AGING CLOCKS

Epigenetics is the branch of molecular biology that aims to understand how genes are regulated (i.e., if DNA is viewed as the hardware of genes, the epigenome can be considered as the software). The best characterized epigenetic mechanism is DNA methylation, which serves as a key signal in genomic regulation. Epigenetic aging clocks are statistical models that can predict the chronological age of healthy individuals from DNA methylation, notorious for their unprecedented accuracy unmet by any other biomarker so far. Namely, these statistical clocks employ specific sites in the human genome whose DNA methylation ratios change monotonically with age in a synchronized fashion across human individuals for reasons not well understood yet. For example, the Horvath’s clock ^29^ is based on an elastic net regression model and includes methylation ratios of 353 sites. In an independent dataset it was able to predict age with a mean absolute error of 3.6 years within a wide range of tissues. Interestingly, when applied on samples from non-healthy individuals (Down Syndrome, Parkinson’s and Alzheimer’s disease and certain types of cancer), the predicted age is often higher than the true chronological age (epigenetic age acceleration) ^30^. On the other hand, when applied to centenarians and their offspring, predicted age is significantly lower than chronological age (epigenetic age deceleration; i.e., association with extreme longevity) ^31^. The prospect that epigenetic aging clocks may serve as a window to the aging process has raised strong biomedical interest since it may result in promising avenues towards the reduction of chronic diseases in the world’s ever-aging population and a potential relief from the health care burden ^16^. Missing values are common for epigenetic aging clocks ^32^, especially in single-cell data ^16^, formalin-fixation and paraffin-embedding (FFPE) samples ^33^, cell-free DNA (cfDNA) from liquid biopsies ^34^ and low-quality and quantity trace samples in forensics ^17^ or paleoepigenetics ^35^. The current state-of-the-art is to perform imputation prior to linear model prediction ^36^, hence, justifying aps-lm’s potential in this application.

To assess the applicability of aps-lm in a real world setting, we made use of the DNA methylation microarray data freely available at the EWAS Data Hub ^37^, targeting 485,512 cytosines and consisting of 8,374 samples from a wide range of tissues and ages. We restricted our analysis to the 353 markers included in the Horvath’s model and we split the dataset in a 9:1 ratio into a reference dataset *X*_ref_, *y*_ref_ (n = 7,536) belonging to stage (i) and an application dataset *X*_app_, *y*_app_ (n = 838) belonging to stage (ii) where *X* refers to the methylation values and *y* to the chronological age. Like in the simulations, *y*_app_ is supposed to be not known but to be estimated by the prediction models. Unlike the set-up for aps-lm, missing values were now present for both *X*_ref_ and *X*_app_ with a proportion of 4.2 %. This required imputation for *X*_ref_. We tested aps-lm and additionally aps-ridge, since the original Horvath’s epigenetic aging model was developed by elastic net regression. Thus, we had a total of four scenarios (more details in Suppl. Methods 7): *scenario 1*: mean imputation on *X*_ref_ followed by a standard linear model, *scenario 2*: MICE imputation on *X*_ref_ followed by a standard linear model, *scenario 3*: mean imputation on *X*_ref_ followed by a ridge model, and *scenario 4*: MICE imputation on *X*_ref_ followed by a ridge model. For scenarios 3 and 4, the regularization parameter λ was based on 10-fold cross-validation (with the glmnet R-package ^38^). For all models, the response variable *y* (chronological age) was transformed by the piecewise Horvath transformation (Suppl. Methods 7). We now had a similar set-up as in the simulations. As the computational costs of MICE for imputation of *X*_app_ in stage (ii) were prohibitive, we solely benchmarked our adaptive-predictor-set models against unconditional mean imputation. We shall in the following use the term “training configuration” to denote the imputation of *X*_ref_ in stage (i) (mean or MICE imputation). “Testing configuration” refers to the two tested prediction methods applied on *X*_app_ in stage (ii) (mean imputation or aps-lm/aps-ridge).

Model performance at the original background level of missing values (4.2 %) was roughly similar across training and testing configurations (Figure 7). To test model performance beyond the original proportion of missing values, we artificially injected missing values to *X*_app_ following the MCAR criterion, solely for simplicity. For an extended range of missing value proportions the training configuration (mean or MICE imputation on *X*_ref_) had little influence on 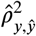. On the other hand, the adaptive predictor-set testing configuration was significantly more resilient to missing value injection than the mean imputation testing configuration in terms of squared correlation and bias; in view of our simulation results, this was to be expected because of the known dependence structure in DNA methylation data. Interestingly, *l*^2^-regularization had a significantly negative impact on the adaptive predictor-set testing configuration. It is well known that ridge performs well in comparison to OLS with respect to the MSE of the parameter vector θ: There is always a λ such that 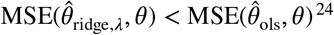 ^24^. In our set-up, however, the MSE of the response *y* is relevant. Here, the relation between ridge and OLS is 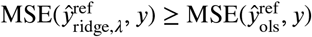 with equality for λ = 0 because OLS minimizes the mean squared error function of *y*. Importantly, this is only true for the reference dataset. In the application dataset there is always a λ such that 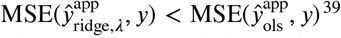 ^39^. We have performed additional simulations with different values of the ratio of the number of observations (*n*) and the number of predictors (*m*) and found that a meaningful lower value of 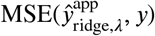 with respect to 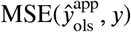 is only reached for a small *n*/*m* (data not shown). For large *n*/*m*, the minimal absolute difference between 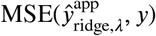 and the respective value for OLS is practically zero and 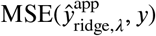 is considerably larger for a wide range of λ. Although a relatively high number of predictors is present in our example dataset (*m* = 353), the number of observations is so high (*n* = 7, 536 in the reference dataset) that *n*/*m* is still large enough such that the advantages of ridge regression do not come into effect. Lastly, we observe that in this application, the adaptive predictor-set testing configuration gives rise to a non-zero bias (though lower than the imputation testing configuration). There are two sources of bias: minor differences in means between reference and application dataset and the use of a non-linear transformation (i.e., Horvath’s piecewise function, *F*) on the dependent variable: 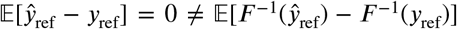. The latter is relevant as we quantify performance on the original scale.

**Figure 7.**
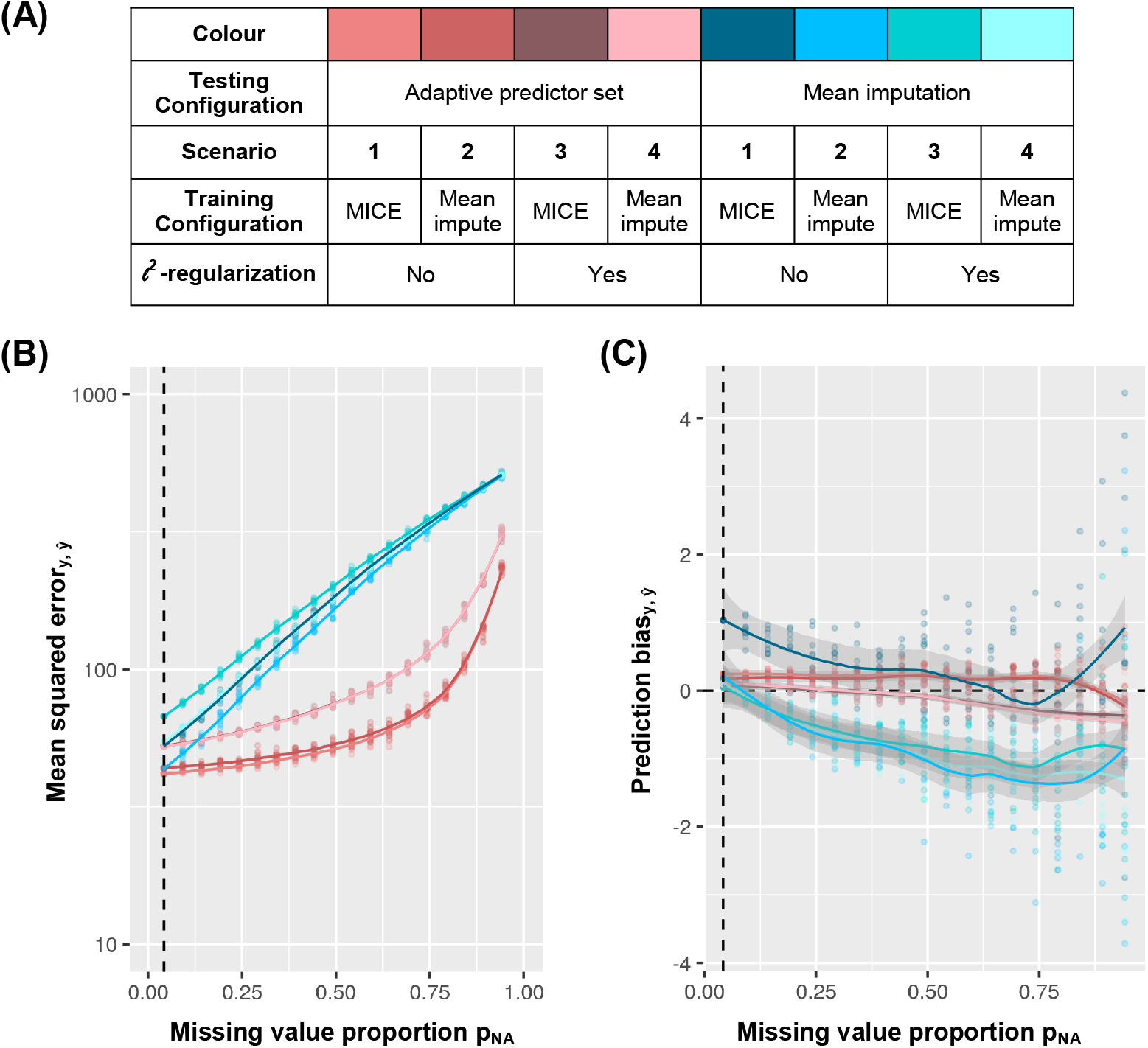
Application of aps-lm on epigenetic aging clocks. The dependent variable consisted of chronological age (in years) on a transformed scale. To test model performance beyond the original proportion of missing values, *p*_0_, we artificially injected missing values following the MCAR scheme. A total of 10 Monte-Carlo replicates were performed for each *p*_NA_ = *p*_0_ +*p*_extra_. (A) Legend with color codes. Mean squared prediction error (B) or prediction bias (C) between real and predicted dependent variable is plotted for varying *p*_NA_, scenarios or testing configurations (mean imputation or aps-lm/aps-ridge). Results are visualized as jitter plots while overall tendencies in the mean are represented as LOESS curves. Dashed vertical line represents *p*_0_.

## 6 DISCUSSION

LR models are omnipresent in statistics. They are, however, incompatible with missing data, a colossal challenge in modern life and health sciences. Complete-case analysis and imputation can aid in parameter estimation of LR models, but not in prediction on an independent dataset, since they either fail to predict on incomplete records or are unable to model how prediction intervals change as a function of the set of missing predictors Ω. In this study, we derived aps-lm as a solution for this problem. It is a generalization of LR that inherently takes the possibility of missing data into account. aps-lm is applied on a reference dataset *X*_ref_, *y*_ref_ in stage (i). Its parameters are then shared in stage (ii) and applied on an incomplete application dataset *X*_app_ to predict the outcome 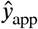 without requiring imputation. In our approach, the full model is adapted to the available predictor sets via equation 5. This submodel is assumed to be the true underlying model and the corresponding OLS-estimator 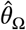 is investigated, which depends on the set of missing variables Ω (equation 6). We have shown that certain parameters *R* and 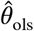 can be calculated from the full model of equation 1 and used to obtain 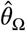. The full model is, however, never assumed to be the true underlying model but only used on a computational basis. The parameters from the full model are projected down to our submodel in order to obtain a good representation of 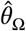. We also developed a version that is applicable to ridge regression, aps-ridge. We have shown that Ω-adapted prediction errors can be easily calculated for aps-lm. Second, aps-lm is CPU-efficient (especially if computed with the sweep operator, Suppl. Methods 5): The parameters of aps-lm are the result of a closed-form solution and there is no need for numerical optimizers and their corresponding inaccuracies and computational burden. Third, aps-lm is storage-efficient: 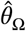 can be written as function of *R* and 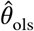, with a total of (*m*+1) ⋅ (*m*+4)/2 parameters, where *m* is the number of predictors (equation 11). This is a substantial compression since it does not depend on the number of observations *n*. With the additional parameter *y*^⊤^*y*, we can also compute custom standard errors that incorporate the degree of missingness in a sample (equation 16). Finally, aps-lm preserves privacy of the reference dataset, since neither *X*_ref_ nor *y*_ref_ can be retrieved from our parametrization *R*,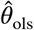 and *y*^⊤^*y*. aps-lm and aps-ridge could only be derived by using the normal equations for the OLS estimator (for aps-lm) and the ridge objective function (for aps-ridge). The adaptive predictor-set model cannot trivially be extended to other regression models such as least absolute deviation, total least squares, lasso regression, elastic nets, robust regression, quantile regression, generalized linear models and Cox regression. Nonetheless, there are possibilities for approximate solutions: As outlined in section 3.6, in a previous study of Marshall *et al* ^10^ a somewhat similar concept to aps-lm was employed in an approximation of parameter estimation in generalized linear models. A reduction of predictors is also considered by Piironen and Vehtari in their “projection predictive method” ^40^. Their equations for the projection of predictors to subsets of predictors are similar to the usage of our matrix operators 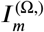 and 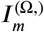. They use this projection to obtain a set Ω that is as close as possible to the reference model (Ω = *∅*) with respect to the Kullback-Leibler divergence.

Our simulations highlighted the power of aps-lm in presence of correlation between predictors. The fact that our aps-lm model outperforms imputation methods might be surprising at first. The underlying reason for that is probably the following: Although multiple imputation strategies like MICE consider multiple predictors as influence variables in their models, MICE performs predictor imputation one at a time using only the predictor to be imputed as an outcome in the modelling equations. Our approach on the other hand takes the full multivariate distribution of all predictors into account via the *R* matrix from the QR decomposition of the design matrix *X*. Another advantage of aps-lm is that information from the whole reference dataset can be utilized via the *R* matrix which is computed from *X*_ref_. So even for independent predictors aps-lm performs better for a high number of missing variables and low sample size than imputation methods. The performance of our method does not depend upon the missingness mechanism and it even gives good results for MNAR. For imputed data, MNAR can introduce large bias and can be worse than CCA because imputation methods like multiple imputation or maximum likelihood usually rely on distributional assumptions on the missingness procedure ^41^. For aps-lm, we assume the existence of a complete reference dataset. By fitting our parameters there, we bypass potential missingness biases occurring at the application stage. Although we attempted to evaluate aps-lm as comprehensively as possible including extreme missingness in our simulations - a scenario often neglected in the literature - there are some limitations to be acknowledged. First, we solely tested aps-lm for a limited number of continuous predictors, covariance structures and missing value types. Applying aps-lm on more datasets in the future will provide the ultimate evidence for its efficacy. Throughout this study, MICE imputation displayed the least favorable performance in spite of its computationally expensive routines. These results are similar to those obtained in the context of Gaussian mixture models by McCaw *et al* ^9^ in which an imputation-free method displayed improved performance over MICE imputation. However, it is important to realize that MICE was not developed in the context of prediction, though many researchers use it for this purpose. Thus, no data shown here shall serve as evidence to discredit MICE’s potential in the context of parameter estimation, for which it was originally derived. In addition, MICE is a very flexible method with a large number of possible settings, which makes it hard to benchmark fairly. For example, we initially observed that MICE was unable to impute when a high proportion of missing values occurred. This was because by default MICE drops collinear variables, which can only be changed by setting the argument `remove.collinear` to `FALSE` (which is not described anywhere on the software’s documentation in version “November 19, 2022”). We used the default for the maximum number of iterations (maxit=5) which may not be sufficient for optimal convergence and also applied the default predictive mean matching (pmm) method with its known danger of over-duplication at a high proportion of missing values. Thus, it is possible that MICE performs more favorable under untested settings.

Furthermore, we applied aps-lm to epigenetic aging clock models as a proof-of-principle. Since our employed dataset contained only 4.2 % missing values, a proportion at which the investigated imputation strategies performed almost indistinguishably from aps-lm, it might seem that there is no need for more refined methods, such as aps-lm. However, it has to be stressed that the dataset employed has been curated and solely includes high-quality data. aps-lm, on the other hand, was derived to obtain predictions even when the data quality is sub-optimal (e.g., for extreme missingness), which enables applications in biological age prediction in FFPE, cfDNA, single-cell or even forensic and paleoepigenetic samples.

The assumption of the existence of completely observed reference data in stage 1 is sometimes not satisfied such that the data of both stage 1 and stage 2 have missing values. Indeed, we had to tackle this very same scenario in our application on epigenetic aging clocks. Nonetheless, having curated (almost) complete reference datasets is a typical situation in biomedical data analysis. A reference dataset is often especially created for model building and thus includes no or only limited missing data. A way around would be to impute missing values in the reference dataset, see our example. Imputation in the reference dataset of stage 1 for model estimation is already well addressed in the literature but not in the case of prediction (e.g., on the application dataset of stage 2).

In summary, the novelty and advantages of our new method are as follows: (i) aps-lm enables the predictive analysis of datasets with missing values without the need for imputation. (ii) aps-lm can be applied regardless of the missing value distribution including MNAR. (iii) aps-lm outperforms imputation approaches. (iv) aps-lm is an analytical method without the need for approximations. (v) aps-lm is computationally more efficient than multiple imputation methods. (vi) aps-lm preserves the privacy of sensitive reference datasets.

Finally, even though we chose for convenience an application whose data were publicly available, we envision our method’s true potential for samples with data protection. Researchers that analyze datasets with privacy restrictions can now make their linear models openly available by sharing the easily computable parameters 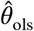 and *R*. Subsequently, these models can be readily applied for any combinations of missing variables by other research groups or for clinical applications.

## ACKNOWLEDGMENTS

We thank Professor Michael Krawczak (Kiel University) for his helpful suggestions. We thank two anonymous reviewers and the associate editor for valuable comments for the improvement of the manuscript.

## DATA AVAILABILITY STATEMENT

R-scripts, raw simulated data and aps-lm models for epigenetic aging prediction with example datasets are available under an MIT license at Github (https://github.com/BenjaminPlanterose/cmblm) and Zenodo (https://doi.org/10.5281/zenodo.6699977). The complete application dataset and the subset of biomarkers employed are publicly available at EWAS Data Hub ^37^ (https://download.cncb.ac.cn/ewas/datahub/download/) and https://github.com/BenjaminPlanterose/cmblm/tree/main/3_application_epigenetic_ageing/data/horvath_age_methylation_v1.txt.zip, respectively.

