## Supplementary Methods for "Adaptive predictor-set linear model: an imputation-free method for linear regression prediction on datasets with missing values"

Amke Caliebe

Institute of Medical Informatics and Statistics, Kiel University, Kiel, Germany
University Medical Centre Schleswig-Holstein, Kiel, Germany

**7 Epigenetic aging clocks** **27**

### 1 Exclude rows/columns operation

In aps-lm models, predictors are removed from the design matrix $X$ depending on the pattern of missing values. To do so in a way compatible with linear algebra, we require a representation of row and column exclusion operations as matrix products. We begin defining a submatrix.

Let $A \in \mathbb{R}^{n \times m}$. A submatrix of $A$ is a matrix formed by selecting rows and columns from $A$ (Clapham and Nicholson (2014)).
Let $\{a_1, a_2, ..., a_r\} \subset \{1, \ldots, n\}$ be the indices of the $r$ rows, where $r \in \{1, \ldots, n\}$.
Let $\{b_1, b_2, ..., b_s\} \subset \{1, \ldots, m\}$ be the indices of the $s$ columns, where $s \in \{1, \ldots, m\}$.

Then the submatrix formed from rows $\{a_1, a_2, ..., a_r\}$ and columns $\{b_1, b_2, ..., b_s\}$ is denoted as follows:

$$A[a_1, a_2, ..., a_r; b_1, b_2, ..., b_s]$$

For example:

$$A = \begin{pmatrix} 1 & 2 & 3 \\ 4 & 5 & 6 \end{pmatrix}; \quad A[1; 1, 3] = \begin{pmatrix} 1 & 3 \end{pmatrix}$$

Let $I_n$ denote the identity matrix of order $n$. We define the following sets: $\Omega_1 = \{a_1, a_2, ..., a_r\}^{\complement}$ and $\Omega_2 = \{b_1, b_2, \ldots, b_s\}^{\complement}$, where $^{\complement}$ denotes the complement set with respect to their universe sets, $U_1 = \{1, \ldots, n\}$ and $U_2 = \{1, \ldots, m\}$, respectively. We define then the following matrices:

$$I_n^{(\Omega_1,)} := I_n[a_1, a_2, ..., a_r; 1, \ldots, n]$$
$$I_m^{(,\Omega_2)} := I_m[1, \ldots, m; b_1, b_2, ..., b_s]$$

It follows that:

$$A^{(\Omega_1,)} := A[a_1, a_2, ..., a_r; 1, \ldots, n] = I_n^{(\Omega_1,)} \cdot A \quad (1)$$
$$A^{(,\Omega_2)} := A[1, \ldots, m; b_1, b_2, ..., b_s] = A \cdot I_m^{(,\Omega_2)} \quad (2)$$

Equations 1 and 2 allow to extend the row/column removal operations to any matrix $X$ by defining the base case on identity matrices and thus, achieving a representation as a matrix product in the language of linear algebra. We here enumerate some useful properties:

$$A \cdot I_m^{(,\emptyset)} = A \cdot I_m = A; \quad I_n^{(\emptyset,)} \cdot A = I_n \cdot A = A$$

where $\emptyset$ denotes the empty set.

$$I_m^{(\Omega,)} \cdot I_m^{(,\Omega)} = I_{m-|\Omega|}; \quad I_m^{(,\Omega)} \cdot I_m^{(\Omega,)} = \text{diag}\left(\mathbf{1}_{\Omega^{\complement}}(1) \;\; \ldots \;\; \mathbf{1}_{\Omega^{\complement}}(m)\right);$$

where $P = \text{diag}(v)$ denotes the diagonal matrix whose components are $P_{i,i} = v_i$ and $P_{i,j \neq i} = 0$ and $\mathbf{1}_{\Phi}(x)$ denotes an indicator function that is equal to 1 if $x \in \Phi$ and equal to 0 when not.

We can also substitute rows and columns for zeros with this operation:

$$(A \cdot I_m^{(,\Omega)} \cdot I_m^{(\Omega,)})_{i,j} = \begin{cases} 0 & j \in \Omega \\ A_{i,j} & j \notin \Omega \end{cases}; \quad (I_n^{(\Omega,)} \cdot I_n^{(,\Omega)} \cdot A)_{i,j} = \begin{cases} 0 & i \in \Omega \\ A_{i,j} & i \notin \Omega \end{cases}$$

### 2 aps-lm model overview

For $n \in \mathbb{N}$ number of observations and $m \in \mathbb{N}$ predictors and given a set missing variables $\Omega$ such that $\Omega \subset \{2, \ldots, (m+1)\}$, we define a aps-lm model as follows:

$$y = X \cdot I_{m+1}^{(,\Omega)} \cdot \theta_\Omega + \epsilon_\Omega; \quad \epsilon_\Omega \sim \mathcal{N}(\vec{0}_n, \sigma^2_{\epsilon,\Omega} \cdot I_n) \tag{3}$$

$$\hat{\theta}_\Omega = (R \cdot I_{m+1}^{(,\Omega)})^\dagger \cdot R \cdot \hat{\theta}_{\text{ols}} \tag{4}$$

where $y \in \mathbb{R}^n$ is a vector containing the dependent variable, $X \in \mathbb{R}^{n \times (m+1)}$ is a full-rank design matrix containing an intercept column, $\theta_\Omega \in \mathbb{R}^{(m+1-|\Omega|)}$ is a submodel linear regression coefficient vector and $\hat{\theta}_\Omega$ its estimator, $\hat{\theta}_{\text{ols}} = (X^\top X)^{-1} X^\top y = X^\dagger y$ (Planitz (1979)) is the vector of coefficients for the full OLS regression model, $R \in \mathbb{R}^{(m+1)\times(m+1)}$ is an upper triangular matrix obtained from the reduced QR decomposition of $X$, $\dagger$ denotes the Moore-Penrose pseudoinverse, $\epsilon_\Omega \in \mathbb{R}^n$ is the vector of error terms, $\vec{0}_n \in \mathbb{R}^n$ is the vector whose components are all equal to zero and $\sigma^2_{\epsilon,\Omega} \in \mathbb{R}$ is the variance of the error terms as a function of $\Omega$. Alternatively, the sweep operator can substitute the pseudoinverse for a more efficient implementation (Suppl. Methods 5)

#### 2.1 Interpreting $R$

$R$ is an upper triangular matrix with $\frac{(m+1)(m+2)}{2}$ non-zero entries that encodes for $n \in \mathbb{N}$, $\hat{\mu} \in \mathbb{R}^m$ (excluding intercept column) and $\hat{\Sigma} \in \mathbb{R}^{m \times m}$ (excluding intercept column) with $\frac{m\cdot(m+1)}{2}$ different parameters; in total, $1 + m + \frac{m\cdot(m+1)}{2} = \frac{(m+1)\cdot(m+2)}{2}$. The proof of the equivalence in parametrization between $R$ and $\{n, \hat{\mu}, \hat{\Sigma}\}$ is shown below.

##### 2.1.1 Obtaining $n$ from $R$

Firstly, given the intercept column:

$$(X^\top X)_{1,1} = (X_{,1})^\top \cdot X_{,1} = \vec{1}_n^\top \cdot \vec{1}_n = n$$

where $X_{,1}$ denotes the first column of $X$ and $\vec{1}_n \in \mathbb{R}^n$ the vector whose components are all equal to one. We also know that:

$$(X^\top X)_{1,1} = ((Q \cdot R)^\top (Q \cdot R))_{1,1} = (R^\top \cdot Q^\top Q \cdot R)_{1,1} = (R_{,1})^\top \cdot R_{,1}$$

As $R$ is upper triangular, thus $(R_{,1})^\top \cdot R_{,1} = R_{1,1}^2 = n$, where $R_{1,1}$ denotes the submatrix $R[1;1]$.

###### 2.1.2 Obtaining $\hat{\mu}$ from $R$

Similarly:

$$(X^\top X)_{1,} = \vec{1}_n^\top \cdot X = n \cdot \hat{\mu}^\top = R_{1,1}^2 \cdot \hat{\mu}^\top$$

$$(X^\top X)_{1,} = \Big((Q \cdot R)^\top (Q \cdot R)\Big)_{1,} = (R_{,1})^\top \cdot R = \begin{pmatrix} R_{1,1} \\ \vec{0}_m \end{pmatrix} \cdot R = R_{1,1} \cdot R_{1,}$$

This means that $(X^\top X)_{1,} = R_{1,1}^2 \cdot \hat{\mu}^\top = R_{1,1} \cdot R_{1,}$ and thus:

$$\hat{\mu} = \left(\frac{R_{1,}}{R_{1,1}}\right)^\top \tag{5}$$

###### 2.1.3 Obtaining $\hat{\Sigma}$ from $R$

Equivalently to $\mathrm{Cov}(X_1, X_2) = \mathbb{E}[X_1 \cdot X_2] - \mathbb{E}[X_1] \cdot \mathbb{E}[X_2]$, we can write:

$$\hat{\Sigma} = \frac{1}{(n-1)} \cdot (X^\top X - n \cdot \hat{\mu} \cdot \hat{\mu}^\top)$$

But we know that $X^\top X = (QR)^\top QR = R^\top R$, that $\hat{\mu} = \left(\frac{R_{1,}}{R_{1,1}}\right)^\top$ and that $n = (R_{1,1})^2$. Thus:

$$\hat{\Sigma} = \frac{R^\top R - (R_{1,})^\top \cdot R_{1,}}{(R_{1,1})^2 - 1} \tag{6}$$

###### 2.1.4 Defining $R$ from the Cholesky decomposition

From the expression for $\hat{\Sigma}$, we can isolate $R^\top R$:

$$R^\top R = (n-1) \cdot \hat{\Sigma} + n \cdot \hat{\mu} \cdot \hat{\mu}^\top$$

Thus:

$$R = \mathrm{chol}((n-1) \cdot \hat{\Sigma} + n \cdot \hat{\mu} \cdot \hat{\mu}^\top) \tag{7}$$

where $\mathrm{chol}(A)$ denotes the matrix $B$ from the Cholesky decomposition with $B^\top \cdot B = A$.

#### 3 Standard errors

In the derivation of the standard errors, we consider the following two linear models, one based on the non-random and unobserved $\theta$ and another based on the random and observed estimate $\hat{\theta}_{\text{ols}}$:

$$y = X \cdot \theta + \epsilon \tag{8}$$

$$y = X \cdot \hat{\theta}_{\text{ols}} + e \tag{9}$$

In these expressions $y$, $\epsilon$, $\hat{\theta}_{\text{ols}}$ and $e$ are random while $X$ and $\theta$ are non-random. Additionally, we assume that $\forall i : \mathbb{E}[\epsilon_i] = 0$ (zero bias in all error random variables) and $\mathbb{E}[\epsilon \cdot \epsilon^\top] = \sigma_\epsilon^2 \cdot I_n$ (homoscedasticity and independence between error random variables). Given all the above, the following relations apply (Greene (2003)):

$$X^\top e = \vec{0}_{m+1} \tag{10}$$

$$\text{Var}(\hat{\theta}_{\text{ols}}) = \sigma_\epsilon^2 \cdot (X^\top X)^{-1} \tag{11}$$

$$\hat{\sigma}_\epsilon^2 = \frac{1}{\nu} e^\top e = \frac{1}{n - (m+1)} e^\top e \tag{12}$$

where $\nu$ is the degrees of freedom. Equivalently, we propose two aps-lm models, one based on the non-random and unobserved $\theta_\Omega$ and another based on the random and observed estimate $\hat{\theta}_\Omega$:

$$y = X \cdot I_{m+1}^{(,\Omega)} \cdot \theta_\Omega + \epsilon_\Omega \tag{13}$$

$$y = X \cdot I_{m+1}^{(,\Omega)} \cdot \hat{\theta}_\Omega + e_\Omega \tag{14}$$

In these expressions $y$, $\epsilon_\Omega$, $\hat{\theta}_\Omega$ and $e_\Omega$ are random while $\Omega$, $X \cdot I_{m+1}^{(,\Omega)}$ and $\theta_\Omega$ are non-random. For the sake of clarity, all variables belong to the reference dataset unless the opposite is specified. Additionally, we assume that $\forall i : \mathbb{E}[\epsilon_{i,\Omega}] = 0$ and $\mathbb{E}[\epsilon_{i,\Omega} \cdot (\epsilon_{i,\Omega})^\top] = \sigma_{\epsilon,\Omega}^2 \cdot I_n$. With these assumptions, the aps-lm estimator is simply the OLS estimator for any combination of predictors (i.e. submodel) and thus OLS properties also apply for aps-lm models. These can be re-written as a function of $\Omega$ as follows:

$$(X \cdot I_{m+1}^{(,\Omega)})^\top \cdot e_\Omega = \vec{0}_{m+1-|\Omega|} \tag{15}$$

$$\text{Var}(\hat{\theta}_\Omega) = \sigma_{\epsilon,\Omega}^2 \cdot ((X \cdot I_{m+1}^{(,\Omega)})^\top \cdot X \cdot I_{m+1}^{(,\Omega)})^{-1} = \sigma_{\epsilon,\Omega}^2 \cdot (I_{m+1}^{(\Omega,)} \cdot R^\top R \cdot I_{m+1}^{(,\Omega)})^\dagger \tag{16}$$

$$\hat{\sigma}_{\epsilon,\Omega}^2 = \frac{1}{\nu_\Omega} \cdot e_\Omega^\top e_\Omega = \frac{1}{n - (m+1) + |\Omega|} \cdot e_\Omega^\top e_\Omega \tag{17}$$

where $\nu_\Omega$ is the degrees of freedom corrected by $\Omega$. To finalize, we need to write $\hat{\sigma}_{\epsilon,\Omega}^2$ as a function of available aps-lm parameters $R$ and $\hat{\theta}_{\text{ols}}$. We first expand equation 17 to get:

$$\hat{\sigma}_{\epsilon,\Omega}^2 = \frac{(y - \hat{y}_\Omega)^\top \cdot (y - \hat{y}_\Omega)}{n - (m+1) + |\Omega|} = \frac{y^\top y + \hat{y}_\Omega^\top \hat{y}_\Omega - 2\hat{y}_\Omega^\top y}{n - (m+1) + |\Omega|} \tag{18}$$

where $\hat{y}_\Omega = X \cdot I_{m+1}^{(,\Omega)} \cdot \hat{\theta}_\Omega$. To further simplify, we require the following result $\hat{y}_\Omega^\top y = \hat{y}_\Omega^\top \hat{y}_\Omega$

which is derived below by reordering, incorporating equation 14 and using equation 15:

$$\hat{y}_\Omega^\top y - \hat{y}_\Omega^\top \hat{y}_\Omega = \hat{y}_\Omega^\top \cdot (y - \hat{y}_\Omega) = (X \cdot I_{m+1}^{(,\Omega)} \cdot \hat{\theta}_\Omega)^\top \cdot e_\Omega = (\hat{\theta}_\Omega)^\top \cdot (X \cdot I_{m+1}^{(,\Omega)})^\top \cdot e_\Omega = 0$$

Therefore, $\hat{y}_\Omega^\top y = \hat{y}_\Omega^\top \hat{y}_\Omega$. Applying this identity on equation 18 yields:

$$\hat{\sigma}_{\epsilon,\Omega}^2 = \frac{y^\top y - \hat{y}_\Omega^\top \hat{y}_\Omega}{n - (m+1) + |\Omega|}$$

We can now expand $\hat{y}_\Omega^\top \hat{y}_\Omega$ as follows:

$$\hat{y}_\Omega^\top \hat{y}_\Omega = (\hat{\theta}_\Omega)^\top \cdot I_{m+1}^{(\Omega,)} \cdot X^\top X \cdot I_{m+1}^{(,\Omega)} \cdot \hat{\theta}_\Omega = (\hat{\theta}_\Omega)^\top \cdot I_{m+1}^{(\Omega,)} \cdot R^\top R \cdot I_{m+1}^{(,\Omega)} \cdot \hat{\theta}_\Omega$$

And thus:

$$\hat{\sigma}_{\epsilon,\Omega}^2 = \frac{y^\top y - (\hat{\theta}_\Omega)^\top \cdot I_{m+1}^{(\Omega,)} \cdot R^\top R \cdot I_{m+1}^{(,\Omega)} \cdot \hat{\theta}_\Omega}{n - (m+1) + |\Omega|} \tag{19}$$

An additional parameter, $y^\top y \in \mathbb{R}$ is required. We can also obtain aps-lm confidence and prediction intervals. Let us assume that we have an observation $k$ belonging to the application set $X_{k,\forall j \in \Omega_k^\complement}^{\text{app}}$. We would like to build confidence intervals for the mean. We assume that the same underlying linear model applies for both the reference and the application dataset. To do so, we derive:

$$\hat{y}_k^{\text{app}} = X_{k,\forall j \in \Omega_k^\complement}^{\text{app}} \cdot \hat{\theta}_{\Omega_k}$$

$$\hat{\text{se}}^2(\hat{y}_k^{\text{app}}) = X_{k,\forall j \in \Omega_k^\complement}^{\text{app}} \cdot \hat{\text{se}}^2(\hat{\theta}_\Omega) \cdot (X_{k,\forall j \in \Omega_k^\complement}^{\text{app}})^\top \tag{20}$$

Via asymptotic theory, we can write the confidence intervals for the mean as

$$\hat{y}_k^{\text{app}} \pm t_{\nu_\Omega}^{\alpha/2} \cdot \sqrt{\hat{\text{se}}^2(\hat{y}_k^{\text{app}})} \tag{21}$$

or

$$X_{k,\forall j \in \Omega_k^\complement}^{\text{app}} \cdot \hat{\theta}_{\Omega_k} \pm t_{n-(m+1)+|\Omega_k|}^{\alpha/2} \cdot \hat{\sigma}_{\epsilon,\Omega} \cdot \sqrt{X_{k,\forall j \in \Omega_k^\complement}^{\text{app}} \cdot (I_{m+1}^{(\Omega_k,)} \cdot R^\top R \cdot I_{m+1}^{(,\Omega_k)})^\dagger \cdot (X_{k,\forall j \in \Omega_k^\complement}^{\text{app}})^\top} \tag{22}$$

To obtain the prediction intervals, we write:

$$y_k^{\text{app}} = X_{k,\forall j \in \Omega_k^\complement}^{\text{app}} \cdot \hat{\theta}_{\Omega_k} + e_{\Omega_k}$$

Assuming that $e_{\Omega_k} \perp (X_{k,\forall j \in \Omega_k^\complement}^{\text{app}})^\top \cdot \hat{\theta}_{\Omega_k}$ (which is at least true for the reference dataset; equation 15), then the variance of the sum is the sum of variances:

$$\hat{\text{se}}^2(y_k^{\text{app}}) = X_{k,\forall j \in \Omega_k^\complement}^{\text{app}} \cdot \hat{\text{se}}^2(\hat{\theta}_\Omega) \cdot (X_{k,\forall j \in \Omega_k^\complement}^{\text{app}})^\top + \hat{\sigma}_{\epsilon,\Omega_k}^2 \tag{23}$$

Similar to the confidence intervals, we now write:

$$y_k^{\text{app}} \pm t_{\nu_\Omega}^{\alpha/2} \cdot \sqrt{\hat{\text{se}}^2(y_k^{\text{app}})} \tag{24}$$

or

$$X^{\text{app}}_{k,\forall j \in \Omega_k^{\complement}} \cdot \hat{\theta}_{\Omega_k} \pm t^{\alpha/2}_{n-(m+1)+|\Omega_k|} \cdot \hat{\sigma}_{\epsilon,\Omega} \cdot \sqrt{X^{\text{app}}_{k,\forall j \in \Omega_k^{\complement}} \cdot (I^{(\Omega_k,)}_{m+1} \cdot R^\top R \cdot I^{(,\Omega_k)}_{m+1})^\dagger \cdot (X^{\text{app}}_{k,\forall j \in \Omega_k^{\complement}})^\top + 1} \tag{25}$$

### 4 Adaptive predictor-set ridge regression

Ridge regression (aka $l^2$-regression or Tikhonov regression) is a regularized version of linear regression with an $l^2$-penalty in the objective function:

$$\hat{\theta}_{\text{ridge}} = \underset{\theta \in \mathbb{R}^{m+1}}{\text{arg.min}} \left( ||y - X \cdot \theta||_2^2 + \lambda \cdot ||I^{(\{1\},)}_{m+1} \cdot \theta||_2^2 \right)$$

where $\lambda$ is the regularization parameter (equivalent to a Lagrange multiplier) and where $I^{(\{1\},)}_{m+1}$ is included in the regularization term to avoid penalizing the intercept variable. Conveniently, a closed-form solution is also available for this problem:

$$\hat{\theta}_{\text{ridge}} = (X^\top X + \lambda \cdot I^{(,\{1\})}_{m+1} \cdot I^{(\{1\},)}_{m+1})^{-1} X^\top y$$

where $I^{(,\{1\})}_{m+1} \cdot I^{(\{1\},)}_{m+1}$ is simply:

$$I^{(,\{1\})}_{m+1} \cdot I^{(\{1\},)}_{m+1} = \begin{pmatrix} 0 & 0 & \dots & 0 \\ 0 & 1 & \dots & 0 \\ \vdots & \vdots & \ddots & 0 \\ 0 & 0 & 0 & 1 \end{pmatrix}$$

To write $\hat{\theta}_{\text{ridge}}$ as a function of the parameters in aps-lm, we apply $X^\top X = (QR)^\top (QR) = R^\top R$ (main manuscript, equation 7) and $Q^\top y = R \cdot \hat{\theta}_{\text{ols}}$ (main manuscript, equation 9), obtaining:

$$\hat{\theta}_{\text{ridge}} = (R^\top R + \lambda \cdot I^{(,\{1\})}_{m+1} \cdot I^{(\{1\},)}_{m+1})^{-1} R^\top R \cdot \hat{\theta}_{\text{ols}}$$

To extend $\hat{\theta}_{\text{ridge}}$ to any combination of available predictors, we simply replace $X$ by $(X \cdot I^{(,\Omega)}_{m+1})$ in the initial expression:

$$\hat{\theta}^{\text{ridge}}_\Omega = \left( (X \cdot I^{(,\Omega)}_{m+1})^\top (X \cdot I^{(,\Omega)}_{m+1}) + \lambda \cdot I^{(,\{1\})}_{m+1-|\Omega|} \cdot I^{(\{1\},)}_{m+1-|\Omega|} \right)^{-1} (X \cdot I^{(,\Omega)}_{m+1})^\top y$$

Which simplifies to:

$$\hat{\theta}^{\text{ridge}}_\Omega = \left( I^{(\Omega,)}_{m+1} \cdot R^\top R \cdot I^{(,\Omega)}_{m+1} + \lambda \cdot I^{(,\{1\})}_{m+1-|\Omega|} \cdot I^{(\{1\},)}_{m+1-|\Omega|} \right)^{-1} \cdot I^{(\Omega,)}_{m+1} \cdot R^\top R \cdot \hat{\theta}_{\text{ols}}$$

To continue, we will assume that $1 \notin \Omega$ (i.e. the model has an intercept) to use the following expression:

$$I_{m+1-|\Omega|}^{(,\{1\})} \cdot I_{m+1-|\Omega|}^{(\{1\},)} = I_{m+1}^{(\Omega,)} \cdot I_{m+1}^{(,\{1\})} \cdot I_{m+1}^{(\{1\},)} \cdot I_{m+1}^{(,\Omega)}$$

Then:

$$\hat{\theta}_{\Omega}^{\text{ridge}} = \left(I_{m+1}^{(\Omega,)} \cdot R^{\top}R \cdot I_{m+1}^{(,\Omega)} + \lambda \cdot I_{m+1}^{(\Omega,)} \cdot I_{m+1}^{(,\{1\})} \cdot I_{m+1}^{(\{1\},)} \cdot I_{m+1}^{(,\Omega)}\right)^{-1} \cdot I_{m+1}^{(\Omega,)} \cdot R^{\top}R \cdot \hat{\theta}_{\text{ols}}$$

$$\hat{\theta}_{\Omega}^{\text{ridge}} = \left(I_{m+1}^{(\Omega,)} \cdot (R^{\top}R + \lambda \cdot I_{m+1}^{(,\{1\})} \cdot I_{m+1}^{(\{1\},)}) \cdot I_{m+1}^{(,\Omega)}\right)^{-1} \cdot I_{m+1}^{(\Omega,)} \cdot R^{\top}R \cdot \hat{\theta}_{\text{ols}}$$

We define $R_{\lambda}$ as

$$R_{\lambda} = \text{chol}(R^{\top}R + \lambda \cdot I_{m+1}^{(,\{1\})} \cdot I_{m+1}^{(\{1\},)})$$

such that $R_{\lambda}^{\top}R_{\lambda} = (R^{\top}R + \lambda \cdot I_{m+1}^{(,\{1\})} \cdot I_{m+1}^{(\{1\},)})$. Thus:

$$I_{m+1}^{(\Omega,)} \cdot (R^{\top}R \cdot +\lambda \cdot I_{m+1}^{(,\{1\})} \cdot I_{m+1}^{(\{1\},)}) \cdot I_{m+1}^{(,\Omega)} = I_{m+1}^{(\Omega,)} \cdot R_{\lambda}^{\top}R_{\lambda} \cdot I_{m+1}^{(,\Omega)}$$

Writing both the expressions for $\hat{\theta}_{\text{ridge}}$ and $\hat{\theta}_{\Omega}^{\text{ridge}}$ as a function of $R_{\lambda}$:

$$\hat{\theta}_{\text{ridge}} = (R_{\lambda}^{\top}R_{\lambda})^{-1} \cdot R^{\top}R \cdot \hat{\theta}_{\text{ols}} \tag{26}$$

$$\hat{\theta}_{\Omega}^{\text{ridge}} = (I_{m+1}^{(\Omega,)} \cdot R_{\lambda}^{\top}R_{\lambda} \cdot I_{m+1}^{(,\Omega)})^{\dagger} \cdot I_{m+1}^{(\Omega,)} \cdot R^{\top}R \cdot \hat{\theta}_{\text{ols}} \tag{27}$$

where for $\hat{\theta}_{\Omega}^{\text{ridge}}$ we additionally replace the inverse for the Moore-Penrose pseudoinverse. Taking advantage of the property $(A^{\top}A)^{\dagger} = A^{\dagger} \cdot (A^{\top})^{\dagger}$, equation 27 becomes:

$$\hat{\theta}_{\Omega}^{\text{ridge}} = (R_{\lambda} \cdot I_{m+1}^{(,\Omega)})^{\dagger} \cdot (I_{m+1}^{(\Omega,)} \cdot R_{\lambda}^{\top})^{\dagger} \cdot I_{m+1}^{(\Omega,)} \cdot R^{\top}R \cdot \hat{\theta}_{\text{ols}} \tag{28}$$

Rearranging expression 26, we get $R_{\lambda}^{\top}R_{\lambda} \cdot \hat{\theta}_{\text{ridge}} = R^{\top}R \cdot \hat{\theta}_{\text{ols}}$. Plugging it into expression 28, we reach the following:

$$\hat{\theta}_{\Omega}^{\text{ridge}} = (R_{\lambda} \cdot I_{m+1}^{(,\Omega)})^{\dagger} \cdot (I_{m+1}^{(\Omega,)} \cdot R_{\lambda}^{\top})^{\dagger} \cdot (I_{m+1}^{(\Omega,)} \cdot R_{\lambda}^{\top}) \cdot R_{\lambda} \cdot \hat{\theta}_{\text{ridge}}$$

Or simply

$$\hat{\theta}_{\Omega}^{\text{ridge}} = (R_{\lambda} \cdot I_{m+1}^{(,\Omega)})^{\dagger} \cdot R_{\lambda} \cdot \hat{\theta}_{\text{ridge}} \tag{29}$$

This last expression is parallel to that of aps-lm (equation 4) but replacing $R$ and $\hat{\theta}_{\text{ols}}$ for $R_{\lambda}$ and $\hat{\theta}_{\text{ridge}}$. Similarly, we can substitute the pseudoinverse for the more efficient sweep operator (Suppl. Methods 5).

In addition, we can derive aps-ridge standard errors. We first begin from the ridge standard errors (Taboga (2021)):

$$\text{Var}(\hat{\theta}_{\text{ridge}}) = \sigma_{\epsilon}^{2} \cdot (X^{\top}X + \lambda \cdot I_{m+1}^{(,\{1\})} \cdot I_{m+1}^{(\{1\},)})^{-1} \cdot X^{\top}X \cdot (X^{\top}X + \lambda \cdot I_{m+1}^{(,\{1\})} \cdot I_{m+1}^{(\{1\},)})^{-1}$$

whose associated estimator is the following:

$$\hat{\text{se}}^2(\hat{\theta}_{\text{ridge}}) = \hat{\sigma}^2_{\epsilon,\lambda} \cdot (X^\top X + \lambda \cdot I^{(,\{1\})}_{m+1} \cdot I^{(\{1\},)}_{m+1})^{-1} \cdot X^\top X \cdot (X^\top X + \lambda \cdot I^{(,\{1\})}_{m+1} \cdot I^{(\{1\},)}_{m+1})^{-1}$$

To adapt this expression to aps-ridge, we exchange $X$ for $(X \cdot I^{(,\Omega)}_{m+1})$ and simplify:

$$\hat{\text{se}}^2(\hat{\theta}^{\text{ridge}}_{\Omega}) = \hat{\sigma}^2_{\epsilon,\lambda,\Omega} \cdot [I^{(\Omega,)}_{m+1} \cdot (R^\top R + \lambda \cdot I^{(,\{1\})}_{m+1} \cdot I^{(\{1\},)}_{m+1}) \cdot I^{(,\Omega)}_{m+1}]^{-1} \cdot I^{(\Omega,)}_{m+1} \cdot R^\top R \cdot I^{(,\Omega)}_{m+1} \cdot [I^{(\Omega,)}_{m+1} \cdot (R^\top R + \lambda \cdot I^{(,\{1\})}_{m+1} \cdot I^{(\{1\},)}_{m+1}) \cdot I^{(,\Omega)}_{m+1}]^{-1}$$

Such expression contain hybrids of $R$ and $R_\lambda$. We can conveniently transform ones into the others as

$$R_\lambda = \text{chol}(R^\top R + \lambda \cdot I^{(,\{1\})}_{m+1} \cdot I^{(\{1\},)}_{m+1}) \tag{30}$$

$$R = \text{chol}(R_\lambda^\top R_\lambda - \lambda \cdot I^{(,\{1\})}_{m+1} \cdot I^{(\{1\},)}_{m+1}) \tag{31}$$

To further simplify:

$$\hat{\text{se}}^2(\hat{\theta}^{\text{ridge}}_{\Omega}) = \hat{\sigma}^2_{\epsilon,\lambda,\Omega} \cdot [I^{(\Omega,)}_{m+1} \cdot R_\lambda^\top R_\lambda \cdot I^{(,\Omega)}_{m+1}]^{-1} \cdot I^{(\Omega,)}_{m+1} \cdot (R_\lambda^\top R_\lambda - \lambda \cdot I^{(,\{1\})}_{m+1} \cdot I^{(\{1\},)}_{m+1}) \cdot I^{(\Omega,)}_{m+1} \cdot [I^{(\Omega,)}_{m+1} \cdot R_\lambda^\top R_\lambda \cdot I^{(,\Omega)}_{m+1}]^{-1}$$

$$\hat{\text{se}}^2(\hat{\theta}^{\text{ridge}}_{\Omega}) = \hat{\sigma}^2_{\epsilon,\lambda,\Omega} \cdot (I^{(\Omega,)}_{m+1} \cdot R_\lambda^\top R_\lambda \cdot I^{(,\Omega)}_{m+1})^{-1} \cdot [I_{m+1-|\Omega|} - \lambda \cdot (I^{(,\{1\})}_{m+1-|\Omega|} \cdot I^{(\{1\},)}_{m+1-|\Omega|}) \cdot (I^{(\Omega,)}_{m+1} \cdot R_\lambda^\top R_\lambda \cdot I^{(,\Omega)}_{m+1})^{-1}]$$

To compute $\hat{\sigma}^2_{\epsilon,\lambda,\Omega}$:

$$\hat{\sigma}^2_{\epsilon,\lambda,\Omega} = \frac{1}{\nu_{\lambda,\Omega}} \cdot (y - X \cdot I^{(,\Omega)}_{m+1} \cdot \hat{\theta}^{\text{ridge}}_{\Omega})^\top (y - X \cdot I^{(,\Omega)}_{m+1} \cdot \hat{\theta}^{\text{ridge}}_{\Omega})$$

where $\nu_{\lambda,\Omega}$ are the effective degrees of freedom. We can expand and simplify as

$$\hat{\sigma}^2_{\epsilon,\lambda,\Omega} = \frac{1}{\nu_{\lambda,\Omega}} \cdot (y^\top y + (\hat{\theta}^{\text{ridge}}_{\Omega})^\top \cdot I^{(\Omega,)}_{m+1} \cdot X^\top X \cdot I^{(,\Omega)}_{m+1} \cdot \hat{\theta}^{\text{ridge}}_{\Omega} - 2 \cdot (\hat{\theta}^{\text{ridge}}_{\Omega})^\top \cdot I^{(\Omega,)}_{m+1} \cdot X^\top \cdot y)$$

Using $X^\top X = R^\top R$ (main manuscript, equation 7) and $Q^\top y = R \cdot \hat{\theta}_{\text{ols}}$ (main manuscript, equation 9), we get:

$$\hat{\sigma}^2_{\epsilon,\lambda,\Omega} = \frac{1}{\nu_{\lambda,\Omega}} \cdot (y^\top y + (\hat{\theta}^{\text{ridge}}_{\Omega})^\top \cdot I^{(\Omega,)}_{m+1} \cdot R^\top R \cdot I^{(,\Omega)}_{m+1} \cdot \hat{\theta}^{\text{ridge}}_{\Omega} - 2 \cdot (\hat{\theta}^{\text{ridge}}_{\Omega})^\top \cdot I^{(\Omega,)}_{m+1} \cdot R^\top R \cdot \hat{\theta}_{\text{ols}})$$

Using $R^\top R \cdot \hat{\theta}_{\text{ols}} = R_\lambda^\top R_\lambda \cdot \hat{\theta}_{\text{ridge}}$ (rearranging equation 26) and $R^\top R = R_\lambda^\top R_\lambda - \lambda \cdot I^{(,\{1\})}_{m+1} \cdot I^{(\{1\},)}_{m+1}$ (from equation 31):

$$\hat{\sigma}^2_{\epsilon,\lambda,\Omega} = \frac{1}{\nu_{\lambda,\Omega}} \cdot (y^\top y + (\hat{\theta}^{\text{ridge}}_{\Omega})^\top \cdot I^{(\Omega,)}_{m+1} \cdot (R_\lambda^\top R_\lambda - \lambda \cdot I^{(,\{1\})}_{m+1} \cdot I^{(\{1\},)}_{m+1}) \cdot I^{(,\Omega)}_{m+1} \cdot \hat{\theta}^{\text{ridge}}_{\Omega} - 2 \cdot (\hat{\theta}^{\text{ridge}}_{\Omega})^\top \cdot I^{(\Omega,)}_{m+1} \cdot R_\lambda^\top R_\lambda \cdot \hat{\theta}_{\text{ridge}})$$

This expression expands into the following large equation:

$$\hat{\sigma}^2_{\epsilon,\lambda,\Omega} = \frac{1}{\nu_{\lambda,\Omega}} \cdot [y^\top y + (\hat{\theta}^{\text{ridge}}_\Omega)^\top \cdot I^{(\Omega,)}_{m+1} \cdot R^\top_\lambda R_\lambda \cdot I^{(,\Omega)}_{m+1} \cdot \hat{\theta}^{\text{ridge}}_\Omega$$
$$- \lambda(\hat{\theta}^{\text{ridge}}_\Omega)^\top \cdot I^{(\Omega,)}_{m+1} \cdot I^{(,\{1\})}_{m+1} \cdot I^{(\{1\},)}_{m+1} \cdot I^{(,\Omega)}_{m+1} \cdot \hat{\theta}^{\text{ridge}}_\Omega - 2 \cdot (\hat{\theta}^{\text{ridge}}_\Omega)^\top \cdot I^{(\Omega,)}_{m+1} \cdot R^\top_\lambda R_\lambda \cdot \hat{\theta}_{\text{ridge}}]$$

To simplify, we need to use the following results:

$$\lambda\cdot(\hat{\theta}^{\text{ridge}}_\Omega)^\top\cdot I^{(\Omega,)}_{m+1}\cdot I^{(,\{1\})}_{m+1}\cdot I^{(\{1\},)}_{m+1}\cdot I^{(,\Omega)}_{m+1}\cdot\hat{\theta}^{\text{ridge}}_\Omega = \lambda\cdot(\hat{\theta}^{\text{ridge}}_\Omega)^\top\cdot I^{(,\{1\})}_{m+1-|\Omega|}\cdot I^{(\{1\},)}_{m+1-|\Omega|}\cdot\hat{\theta}^{\text{ridge}}_\Omega = \lambda\cdot||I_{m+1-|\Omega|}\cdot\hat{\theta}^{\text{ridge}}_\Omega||^2_2$$

and:

$$(\hat{\theta}^{\text{ridge}}_\Omega)^\top\cdot I^{(\Omega,)}_{m+1}\cdot R^\top_\lambda R_\lambda\cdot I^{(,\Omega)}_{m+1}\cdot\hat{\theta}^{\text{ridge}}_\Omega - 2\cdot(\hat{\theta}^{\text{ridge}}_\Omega)^\top\cdot I^{(\Omega,)}_{m+1}\cdot R^\top_\lambda R_\lambda\cdot\hat{\theta}_{\text{ridge}} = -(\hat{\theta}^{\text{ridge}}_\Omega)^\top\cdot I^{(\Omega,)}_{m+1}\cdot R^\top_\lambda R_\lambda\cdot I^{(,\Omega)}_{m+1}\cdot\hat{\theta}^{\text{ridge}}_\Omega$$

The latter follows since $R_\lambda \cdot \hat{\theta}_{\text{ridge}} = (R_\lambda \cdot I^{(,\Omega)}_{m+1}) \cdot \hat{\theta}^{\text{ridge}}_\Omega$ (rearranging equation 29). As a result:

$$\hat{\sigma}^2_{\epsilon,\lambda,\Omega} = \frac{1}{\nu_{\lambda,\Omega}} \cdot (y^\top y - (\hat{\theta}^{\text{ridge}}_\Omega)^\top \cdot I^{(\Omega,)}_{m+1} \cdot R^\top_\lambda R_\lambda \cdot I^{(,\Omega)}_{m+1} \cdot \hat{\theta}^{\text{ridge}}_\Omega - \lambda \cdot ||I_{m+1-|\Omega|} \cdot \hat{\theta}^{\text{ridge}}_\Omega||^2_2) \quad (32)$$

This expression is parallel to aps-lm variance of residuals (equation 19) but introducing a regularization term. Also, relative to this expression, the concept of degrees of freedom is less straightforward to define in ridge regression since the number of free parameters is undefined. We can generalize the concept of degrees of freedom (effective or equivalent degrees of freedom; Janson et al. (2015)) as the trace of the hat matrix $H$; $H \in \mathbb{R}^{n\times n}$ is a symmetric matrix that transforms $y$ into $\hat{y}$ as in $\hat{y} = H \cdot y$ (hence its name). For OLS, since $\hat{\theta}_{\text{ols}} = X^\dagger \cdot y$ and $\hat{y} = X \cdot \hat{\theta}_{\text{ols}} = X \cdot X^\dagger \cdot y$, the hat matrix is simply $H = X \cdot X^\dagger = I_{m+1}$. The degrees of freedom $\nu$ are then:

$$\nu = n - \text{Tr}(H) = n - \text{Tr}(I_{m+1}) = n - (m + 1)$$

which coincides with the standard definition. For aps-lm, $H_\Omega = (X \cdot I^{(,\Omega)}_{m+1}) \cdot (X \cdot I^{(,\Omega)}_{m+1})^\dagger = I_{m+1-|\Omega|}$ and thus:

$$\nu_\Omega = n - \text{Tr}(H_\Omega) = n - \text{Tr}(I_{m+1-|\Omega|}) = n - (m + 1) + \Omega$$

For ridge, since:

$$\hat{\theta}_{\text{ridge}} = (R^\top_\lambda R_\lambda)^{-1} \cdot R^\top R \cdot \hat{\theta}_{\text{ols}} = (R^\top_\lambda R_\lambda)^{-1} \cdot R^\top R \cdot X^\dagger \cdot y; \quad \hat{y}_\lambda = X \cdot \hat{\theta}_{\text{ridge}}$$

the hat matrix is simply $H_\lambda = X \cdot (R^\top_\lambda R_\lambda)^{-1} \cdot R^\top R \cdot X^\dagger$ and thus:

$$\nu_\lambda = n - \text{Tr}(H_\lambda) = n - \text{Tr}(X \cdot (R^\top_\lambda R_\lambda)^{-1} \cdot R^\top R \cdot X^\dagger) = n - \text{Tr}(X^\dagger \cdot X \cdot (R^\top_\lambda R_\lambda)^{-1} \cdot R^\top R) = n - \text{Tr}\left((R^\top_\lambda R_\lambda)^{-1} \cdot R^\top R\right)$$

where we use the cyclic property of the trace operator. As $\lambda$ tends to 0, $\nu_\lambda$ tends to $\nu$.

Similarly, for aps-ridge we get:

$$H_{\lambda,\Omega} = X \cdot I_{m+1}^{(,\Omega)} \cdot (R_\lambda \cdot I_{m+1}^{(,\Omega)})^\dagger \cdot R_\lambda \cdot (R_\lambda^\top R_\lambda)^{-1} \cdot R^\top R \cdot X^\dagger$$

$$\nu_{\lambda,\Omega} = n - \operatorname{Tr}\left(I_{m+1}^{(,\Omega)} \cdot (R_\lambda \cdot I_{m+1}^{(,\Omega)})^\dagger \cdot R_\lambda \cdot (R_\lambda^\top R_\lambda)^{-1} \cdot R^\top R\right)$$

As a sanity check, as $\lambda$ tends to 0, $R_\lambda$ should tend to $R$ and thus $\nu_{\lambda,\Omega}$ should tend to $\nu_\Omega$:

$$\nu_{\lambda\to 0,\Omega} = n - \operatorname{Tr}\left(I_{m+1}^{(,\Omega)} \cdot (R.I_{m+1}^{(,\Omega)})^\dagger \cdot R \cdot (R^\top R)^{-1} \cdot R^\top R\right) = n - \operatorname{Tr}\left((R \cdot I_{m+1}^{(,\Omega)}) \cdot (R.I_{m+1}^{(,\Omega)})^\dagger\right) = \nu_\Omega$$

Also, when $\Omega = \emptyset$, $\nu_{\lambda,\emptyset} = \nu_\lambda$:

$$\nu_{\lambda,\emptyset} = n - \operatorname{Tr}\left(I_{m+1}^{(,\emptyset)} \cdot (R_\lambda \cdot I_{m+1}^{(,\emptyset)})^\dagger \cdot R_\lambda \cdot (R_\lambda^\top R_\lambda)^{-1} \cdot R^\top R\right) = n - \operatorname{Tr}\left((R_\lambda^\top R_\lambda)^{-1} \cdot R^\top R\right) = \nu_\lambda$$

We can also derive confidence and prediction intervals for aps-ridge. Let us assume that we have an observation $k$ belonging to the application set $X^{\text{app}}_{k,\forall j \in \Omega_k^\complement}$. Then, the following approximation follows:

$$\hat{y}_k = X^{\text{app}}_{k,\forall j \in \Omega_k^\complement} \cdot \hat{\theta}^{\text{ridge}}_{\Omega_k}; \quad \hat{y}_k \pm t^{\alpha/2}_{\nu_{\lambda,\Omega_k}} \cdot \sqrt{\hat{\text{se}}^2(\hat{y}_k)} \tag{33}$$

$$X^{\text{app}}_{k,\forall j \in \Omega_k^\complement} \cdot \hat{\theta}^{\text{ridge}}_{\Omega_k} \pm t^{\alpha/2}_{\nu_{\lambda,\Omega_k}} \cdot \sqrt{X^{\text{app}}_{k,\forall j \in \Omega_k^\complement} \cdot \hat{\text{se}}^2(\hat{\theta}^{\text{ridge}}_{\Omega_k}) \cdot (X^{\text{app}}_{k,\forall j \in \Omega_k^\complement})^\top}$$

and prediction intervals as

$$y_k = X^{\text{app}}_{k,\forall j \in \Omega_k^\complement} \cdot \hat{\theta}^{\text{ridge}}_{\Omega_k} + \epsilon_{\Omega_k}; \quad y_k \pm t^{\alpha/2}_{\nu_{\lambda,\Omega_k}} \cdot \sqrt{\hat{\text{se}}^2(y_k)} \tag{34}$$

$$X^{\text{app}}_{k,\forall j \in \Omega_k^\complement} \cdot \hat{\theta}^{\text{ridge}}_{\Omega_k} \pm t^{\alpha/2}_{\nu_{\lambda,\Omega_k}} \cdot \sqrt{X^{\text{app}}_{k,\forall j \in \Omega_k^\complement} \cdot \hat{\text{se}}^2(\hat{\theta}^{\text{ridge}}_{\Omega_k}) \cdot (X^{\text{app}}_{k,\forall j \in \Omega_k^\complement})^\top + \hat{\sigma}^2_{\epsilon,\lambda,\Omega_k}}$$

This is an approximation since the distribution of the ridge estimator is unknown.

#### 4.1 aps-ridge: bias

The ridge estimator is biased (Hoerl and Kennard (2000)), and thus so is the aps-ridge estimator. Precisely:

$$\mathbb{E}[\hat{\theta}_{\text{ridge}}] - \theta = [(X^\top X + \lambda \cdot I_{m+1}^{(,\{1\})} \cdot I_{m+1}^{(\{1\},)})^{-1} - (X^\top X)^{-1}] \cdot X^\top X \cdot \theta$$

This, however, does not mean that the prediction on the dependent variable is biased:

$$\mathbb{E}[y - X \cdot \hat{\theta}_{\text{ridge}}] = \mathbb{E}[e_\lambda] = 0; \quad \mathbb{E}[X \cdot \hat{\theta}_{\text{ridge}}] = \bar{y}$$

where $e_\lambda$ denotes ridge residuals and $\bar{y}$ the mean of $y$. To prove this, we expand the expected value expression with the definition of $\hat{\theta}_{\text{ridge}}$ (equation 26) and $R^\top R = R_\lambda^\top R_\lambda - \lambda \cdot I_{m+1}^{(,\{1\})} \cdot I_{m+1}^{(\{1\},)}$ (equation 31):

$$\mathbb{E}[X \cdot \hat{\theta}_{\text{ridge}}] = \mathbb{E}[X \cdot (R_\lambda^\top R_\lambda)^{-1} \cdot R^\top R \cdot \hat{\theta}_{\text{ols}}] = \mathbb{E}[X \cdot \hat{\theta}_{\text{ols}}] - \lambda \cdot \mathbb{E}[X \cdot (R_\lambda^\top R_\lambda)^{-1} \cdot I_{m+1}^{(,\{1\})} \cdot I_{m+1}^{(\{1\},)} \cdot \hat{\theta}_{\text{ols}}]$$

The difference in means between ridge and OLS is:

$$\mathbb{E}[X \cdot \hat{\theta}_{\text{ridge}}] - \mathbb{E}[X \cdot \hat{\theta}_{\text{ols}}] = -\frac{\lambda}{n} \cdot \vec{1}_n^\top \cdot X \cdot (R_\lambda^\top R_\lambda)^{-1} \cdot I_{m+1}^{(,\{1\})} \cdot I_{m+1}^{(\{1\},)} \cdot \hat{\theta}_{\text{ols}} = -\lambda \cdot \hat{\mu}^\top \cdot (R_\lambda^\top R_\lambda)^{-1} \cdot I_{m+1}^{(,\{1\})} \cdot I_{m+1}^{(\{1\},)} \cdot \hat{\theta}_{\text{ols}}$$

To further simplify, we need to write $\hat{\mu}$ as a function of $R_\lambda^\top R_\lambda$ taking advantage of the intercept column of X:

$$\hat{\mu} = \frac{1}{n} \cdot X^\top \cdot \vec{1}_n = \frac{1}{n} \cdot X^\top X \cdot I_{m+1}^{(,2:(m+1))} = \frac{1}{n} \cdot R^\top R \cdot I_{m+1}^{(,2:(m+1))} = \frac{1}{n} \cdot R_\lambda^\top R_\lambda \cdot I_{m+1}^{(,2:(m+1))}$$

In the last step, we take advantage that $\lambda \cdot I_{m+1}^{(,\{1\})} \cdot I_{m+1}^{(\{1\},)}$ does not modify the first row of $R^\top R$ and thus, is equal to the first row of $R_\lambda^\top R_\lambda$. Plugging into the expression X gives rise to:

$$\mathbb{E}[X \cdot \hat{\theta}_{\text{ridge}}] - \mathbb{E}[X \cdot \hat{\theta}_{\text{ols}}] = -\frac{\lambda}{n} \cdot I_{m+1}^{(2:(m+1),)} \cdot I_{m+1}^{(,\{1\})} \cdot I_{m+1}^{(\{1\},)} \cdot \hat{\theta}_{\text{ols}} = -\frac{\lambda}{n} \cdot \begin{pmatrix} 1 \\ 0 \\ \vdots \\ 0 \end{pmatrix}^\top \cdot \begin{pmatrix} 0 \\ \hat{\theta}_2^{\text{ols}} \\ \vdots \\ \hat{\theta}_{m+1}^{\text{ols}} \end{pmatrix} = 0$$

Since we know that $\mathbb{E}[X \cdot \hat{\theta}_{\text{ols}}] = \bar{y}$, thus $\mathbb{E}[X \cdot \hat{\theta}_{\text{ridge}}] = \bar{y}$.

### 5 The sweep operator can replace the pseudoinverse

The sweep operator is a highly versatile matrix operation that has been used to optimize the computation of solutions to numerical linear algebra problems in statistics (Goodnight (1979)). The concept is similar to Gaussian elimination or forward Doolittle, which solves linear equations via diagonalization or triangularization-backward substitution, respectively. A sweep operation of the $k^{th}$-element in a square matrix $A$, $\mathbb{S}_k(A) = B$, is defined as

$$\begin{cases} B_{k,k} = \frac{1}{A_{k,k}} & A_{k,k} \neq 0 \\ B_{i,k} = -\frac{A_{i,k}}{A_{k,k}} & \forall i \neq k \\ B_{k,j} = \frac{A_{k,j}}{A_{k,k}} & \forall j \neq k \\ B_{i,k} = A_{i,j} - \frac{A_{i,k} \cdot A_{k,j}}{A_{k,k}} & \forall i, j \neq k \end{cases} \tag{35}$$

For example:

$$\mathbb{S}_1 \begin{pmatrix} 4 & -1 & 3 \\ -1 & 5 & 2 \\ 3 & 2 & 6 \end{pmatrix} = \begin{pmatrix} 1/4 & -1/4 & 3/4 \\ -(-1)/4 & 5 - \frac{(-1)\cdot(-1)}{4} & -\frac{(-1)\cdot 3}{4} \\ -3/4 & 2 - \frac{3\cdot(-1)}{4} & 6 - \frac{3\cdot 3}{4} \end{pmatrix} = \begin{pmatrix} 1/4 & -1/4 & 3/4 \\ 1/4 & 19/4 & 11/4 \\ -3/4 & 11/4 & 15/4 \end{pmatrix}$$

The sweep operator can be computed in the R-programming language with `fastmatrix::sweep.operator` (Osorio F. (2022)).

Replacing the pseudoinverse in our aps-lm model (main manuscript, section 3) by the sweep operator offers a strong reduction in computational costs. To do so, we define the extended matrix $\tilde{X}$ and its matrix of second moments as follows and expand using equations 7 and 9 from the main manuscript.

$$\tilde{X} := (X \quad y)\,; \quad \tilde{X}^\top \tilde{X} = \begin{pmatrix} X^\top X & X^\top y \\ y^\top X & y^\top y \end{pmatrix} = \begin{pmatrix} R^\top R & R^\top R \cdot \hat{\theta}_{\text{ols}} \\ \hat{\theta}_{\text{ols}}^\top \cdot R^\top R & y^\top y \end{pmatrix}$$

According to the theory of the sweep operator (Goodnight (1979)), $\hat{\theta}_{\text{ols}}$ can be retrieved from entries in the extended matrix after performing a sequence of sweeps. For example, let us assume that:

$$X = \begin{pmatrix} 1 & 1 & 4 & 3 \\ 1 & 5 & 3 & 6 \\ 1 & 7 & 8 & 3 \\ 1 & 2 & 1 & 4 \end{pmatrix}; \quad y = \begin{pmatrix} 1 \\ 4 \\ 3 \\ 1 \end{pmatrix}; \quad \hat{\theta}_{\text{ols}} = X^\dagger y = \begin{pmatrix} -39/11 \\ 1/11 \\ 4/11 \\ 1 \end{pmatrix}$$

Thus:

$$\tilde{X}^\top \tilde{X} = \begin{pmatrix} 4 & 15 & 16 & 16 & 9 \\ 15 & 79 & 77 & 62 & 44 \\ 16 & 77 & 90 & 58 & 41 \\ 16 & 62 & 58 & 70 & 40 \\ 9 & 44 & 41 & 40 & 27 \end{pmatrix}; \quad (X^\top X)^{-1} = \begin{pmatrix} 12369/968 & 311/242 & -1569/968 & -239/88 \\ 311/242 & 30/121 & -57/242 & -7/22 \\ -1569/968 & -57/242 & 265/968 & 31/88 \\ -239/88 & -7/22 & 31/88 & 5/8 \end{pmatrix}$$

In this case, by applying $\mathbb{S}_{\{1,\ldots,m+1\}}(\tilde{X}^\top \tilde{X}) = \mathbb{S}_{\{1,2,3,4\}}(\tilde{X}^\top \tilde{X}) = \mathbb{S}_4\mathbb{S}_3\mathbb{S}_2\mathbb{S}_1(\tilde{X}^\top \tilde{X})$ and extracting certain entries from the resulting matrix, we can retrieve $\hat{\theta}_{\text{ols}}$:

$$\mathbb{S}_{\{1,2,3,4\}}(\tilde{X}^\top \tilde{X}) = \begin{pmatrix} 12369/968 & 311/242 & -1569/968 & -239/88 & \mathbf{-39/11} \\ 311/242 & 30/121 & -57/242 & -7/22 & \mathbf{1/11} \\ -1569/968 & -57/242 & 265/968 & 31/88 & \mathbf{4/11} \\ -239/88 & -7/22 & 31/88 & 5/8 & \mathbf{1} \\ 39/11 & -1/11 & -4/11 & -1 & 0 \end{pmatrix} = \begin{pmatrix} (X^\top X)^{-1} & \boldsymbol{\hat{\theta}}_{\text{ols}} \\ -\hat{\theta}_{\text{ols}}^\top & (n-(m+1)) \cdot \hat{\sigma}_\epsilon^2 \end{pmatrix}$$

Please note that the sweep operator is commutative ($\mathbb{S}_l\mathbb{S}_m(A) = \mathbb{S}_m\mathbb{S}_l(A)$), so the order in which the sweeps are performed does not alter the result.

More interestingly for aps-lm, according to the theory of the sweep operator (Goodnight (1979)), we can also retrieve $\hat{\theta}_\Omega$ in terms of the sweep operator by sequentially sweeping $\tilde{X}^\top \tilde{X}$ $\forall k \in \Omega^\complement$ instead of $\{1, \ldots, m+1\}$ and extracting particular entries from the resulting matrix, where $\Omega^{\complement}$ is the set of variables that are not missing. Specifically:

$$\hat{\theta}_{\Omega} = \mathbb{S}_{\Omega^{\complement}}(\tilde{X}^{\top}\tilde{X})[\Omega^{\complement}; m+2] = I_{m+2}^{(\Omega \bigcup \{m+2\},)} \cdot \mathbb{S}_{\Omega^{\complement}}(\tilde{X}^{\top}\tilde{X}) \cdot I_{m+2}^{(,\forall j \neq (m+2))}$$

where $\mathbb{S}_{\Omega^{\complement}}(A)$ denotes the $|\Omega^{\complement}|$ sweep operations on elements $A_{kk} : \forall k \in \Omega^{\complement}$ and $I_{m+2}^{(,\forall j \neq (m+2))}$ is the (m+2)-identity matrix excluding all columns except for column $m+2$. Using again equations 7 and 9 from the main manuscript, we can write $\hat{\theta}_{\Omega}$ as a function of aps-lm parameters $R$, $\hat{\theta}_{\text{ols}}$ and $y^{\top}y$:

$$\hat{\theta}_{\Omega} = I_{m+2}^{(\Omega \bigcup \{m+2\},)} \cdot \mathbb{S}_{\Omega^{\complement}} \begin{pmatrix} R^{\top}R & R^{\top}R \cdot \hat{\theta}_{\text{ols}} \\ \hat{\theta}_{\text{ols}}^{\top} \cdot R^{\top}R & y^{\top}y \end{pmatrix} \cdot I_{m+2}^{(,\forall j \neq (m+2))} \tag{36}$$

Here, entry $(m+2, m+2)$ in $\tilde{X}^{\top}\tilde{X}$, equal to $y^{\top}y$, is included despite not influencing the sweep operation since this entry can be simultaneously used in the estimation of the variance of the errors:

$$\hat{\sigma}_{\epsilon,\Omega}^2 = \frac{1}{n - (m+1) + |\Omega|} \cdot \left[ I_{m+2}^{(\forall j \neq (m+2),)} \cdot \mathbb{S}_{\Omega^{\complement}} \begin{pmatrix} R^{\top}R & R^{\top}R \cdot \hat{\theta}_{\text{ols}} \\ \hat{\theta}_{\text{ols}}^{\top} \cdot R^{\top}R & y^{\top}y \end{pmatrix} \cdot I_{m+2}^{(,\forall j \neq (m+2))} \right] \tag{37}$$

where $I_{m+2}^{(\forall j \neq (m+2),)}$ is the (m+2)-identity matrix excluding all rows except for row $m+2$. Equations 36 and 37 offer more computationally-friendly alternatives to equations 10 and 16 from the main manuscript.

Parallelly for aps-ridge models, $\hat{\theta}_{\Omega}^{\text{ridge}}$ can also be written as a function of the sweep operator:

$$\hat{\theta}_{\Omega}^{\text{ridge}} = I_{m+2}^{(\Omega \bigcup \{m+2\},)} \cdot \mathbb{S}_{\Omega^{\complement}} \begin{pmatrix} R_{\lambda}^{\top}R_{\lambda} & R_{\lambda}^{\top}R_{\lambda} \cdot \hat{\theta}_{\text{ridge}} \\ \hat{\theta}_{\text{ridge}}^{\top} \cdot R_{\lambda}^{\top}R_{\lambda} & y^{\top}y \end{pmatrix} \cdot I_{m+2}^{(,\forall j \neq (m+2))} \tag{38}$$

Like for aps-lm models, $\hat{\sigma}_{\epsilon,\Omega,\lambda}^2$ can also be obtained via the sweep procedure:

$$\hat{\sigma}_{\epsilon,\Omega,\lambda}^2 = \frac{1}{\nu_{\lambda,\Omega}^{\text{ridge}}} \cdot \left[ I_{m+2}^{(\forall j \neq (m+2),)} \cdot \mathbb{S}_{\Omega^{\complement}} \begin{pmatrix} R_{\lambda}^{\top}R_{\lambda} & R_{\lambda}^{\top}R_{\lambda} \cdot \hat{\theta}_{\text{ridge}} \\ \hat{\theta}_{\text{ridge}}^{\top} \cdot R_{\lambda}^{\top}R_{\lambda} & y^{\top}y \end{pmatrix} \cdot I_{m+2}^{(,\forall j \neq (m+2))} - \lambda \cdot ||I_{m+1}^{(\{1\},)} \cdot \hat{\theta}_{\Omega}^{\text{ridge}}||_2^2 \right] \tag{39}$$

In this case, equations 38 and 39 offer reduced computational costs compared to equations 20 and 21 from the main manuscript.

### 6 Simulations

#### 6.1 Types of missing values

For $n \in \mathbb{N}$ number of observations and $m \in \mathbb{N}$ predictors, let $X^0 \in \mathbb{R}^{n \times m}$ be the unobserved design matrix containing the complete record of observations; note that we ignore the intercept column in this section since missing values cannot occur at the intercept variable by definition. Let $M \in \mathbb{R}^{n\times m}$ be a random binary matrix where $M_{i,j} = 1$ if element $X^0_{i,j}$ becomes missing, for $i \in \{1, \ldots, n\}$ and $j \in \{1, \ldots, m\}$. Let $X \in \mathbb{R}^{n\times m}$ be the resulting matrix of observations after suffering a random process that replaces observed entries by missing values according to $M$. Let $W \in \mathbb{R}^{n\times h}$ be the matrix of $h$ non-observed or hidden variables. In the most general case, we are interested in $P(M_{i,j} = 1|\phi_j, X^0_{i,\forall k\neq j}, X^0_{i,j}, W_{i,})$, where $\phi_j$ corresponds to unknown parameters associated to the variable $j$, where $X^0_{i,\forall k\neq j}$ corresponds to the values of the other observed variables $k$ different than $j$ for the same individual $i$ and where $W_{i,}$ corresponds to the vector $(W_{i,1}, \ldots, W_{i,h})^\top$.

The famous classification by D.B. Rubin defines 3 types of missing values (Rubin (1976)):

i) missing completely at random (MCAR) in which $P(M_{i,j} = 1|\phi_j, X^0_{i,\forall k\neq j}, X^0_{i,j}, W_{i,}) = P(M_{i,j} = 1|\phi_j)$,

ii) missing (conditionally) at random (MAR) where $P(M_{i,j} = 1|\phi_j, X^0_{i,\forall k\neq j}, X^0_{i,j}, W_{i,}) = P(M_{i,j} = 1|\phi_j, X^0_{i,\forall k\neq j})$ and

iii) missing not at random (MNAR) for all other cases.

In our simulation, though we use such general labels, we restrict to specific set-ups which are clearly defined below. For this purpose we define as a parameter the average probability of obtaining a missing value $p_{\mathrm{NA}}$ as

$$p_{\mathrm{NA}} = \frac{1}{n\cdot m}\sum_{i=1}^{n}\sum_{j=1}^{m} P(M_{i,j} = 1) \tag{40}$$

##### 6.1.1 MCAR

For our purpose, we use a MCAR model parameterized as follows:

$$P(M_{i,j} = 1|\phi_j, X^0_{i,\forall k\neq j}, X^0_{i,j}, W_{i,}) = P(M_{i,j} = 1|p_{\mathrm{NA}}) = p_{\mathrm{NA}} \tag{41}$$

which satisfies equation 40. Then, $M_{i,j} \sim \text{Binomial}(n = 1, p_{\mathrm{NA}})$ and thus the probability of a sample missing k number of variables follows:

$$P(X = k|p_{\mathrm{NA}}) = \binom{m}{k}\cdot p_{\mathrm{NA}}^k \cdot (1 - p_{\mathrm{NA}})^{m-k}$$

It is important to exclude the case where all variables are missing in a given individual as we would not be trying to perform prediction modelling on an individual if no variable was available. Excluding such cases, sets an upper limit for $p_{\mathrm{NA}}$ (one variable available per individual):

$$p_{\mathrm{NA}} = 1 - p_{\neq\mathrm{NA}} \leq 1 - \frac{n}{n\cdot m} = 1 - 1/m$$

Computing probabilities but excluding the situation with all missing variables can be achieved via renormalization:

$$P(X = k|p_{\text{NA}}) = \frac{1}{1 - p_{\text{NA}}^m} \cdot \binom{m}{k} \cdot p_{\text{NA}}^k \cdot (1 - p_{\text{NA}})^{m-k} \tag{42}$$

An efficient implementation of this missing value random process is a two-stage approach: for each individual, i) decide how many variables are lost and ii) decide which specific variables are lost. This can be implemented as: for each individual $i$, i) we draw from the set $\{1, \dots, (m-1)\}$ a single value $k_i$ by equation 42 and ii) we draw from the set $\{1, \dots, m\}$ a total of $k_i$ times without replacement to form the set of variables to become missing which we denote $\Omega_i$. The missing value indicator matrix will simply be determined as

$$M_{i,j} = \begin{cases} 0 & j \notin \Omega_i \\ 1 & j \in \Omega_i \end{cases}$$

##### 6.1.2 MNAR

For our purpose, we use an MNAR model satisfying:

$$P(M_{i,j} = 1|\phi_j, X^0_{i,\forall k \neq j}, X^0_{i,j}, W_{i,}) = P(M_{i,j} = 1|X^0_{i,j}, \lambda) \tag{43}$$

where $i \in \{1, \dots, n\}$, $j \in \{1, \dots, m\}$ and $\lambda \in \mathbb{R}$ is a parameter. We propose four link functions that relates the value of $X^0_{i,j}$ itself with the probability of becoming missing:

$$\text{Left (L)} \qquad P(M_{i,j} = 1|X^0_{i,j}, \lambda) = \frac{1}{1+e^{-(X^0_{i,j}+\lambda)}}$$

$$\text{Right (R)} \qquad P(M_{i,j} = 1|X^0_{i,j}, \lambda) = \frac{1}{1+e^{-(-X^0_{i,j}+\lambda)}}$$

$$\text{Left-Right (LR)} \qquad P(M_{i,j} = 1|X^0_{i,j}, \lambda) = \frac{1}{1+e^{-(|X^0_{i,j}|+\lambda)}}$$

$$\text{Center (C)} \qquad P(M_{i,j} = 1|X^0_{i,j}, \lambda) = 1 - \frac{1}{1+e^{-(|X^0_{i,j}|+\lambda)}}$$

The parameter $\lambda$ is included to tune the link function in order to control for $p_{\text{NA}}$. To find the value of $\lambda$, we set $p_{\text{NA}}$ and we define the following objective function:

$$g(\lambda) = \frac{1}{m \cdot n} \sum_{i=1}^{n} \sum_{j=1}^{m} P(M_{i,j} = 1|X^0_{i,j}, \lambda) - p_{\text{NA}}$$

We choose $\lambda$ so that $g(\lambda)$ becomes as close to zero as possible. This can be done with a one-dimensional root finding routine (e.g. `stats::uniroot` in R-base). Finally, for a total of $n \times m$ times and per position $i$,$j$ we draw a binomial variable with probability

$P(M_{i,j} = 1|X^0_{i,j}, \lambda)$ to fill the matrix $M$.

As a drawback to this methodology, it does not exclude the possibility of all variables becoming missing in a given sample at high $p_{\text{NA}}$ (unlike the two-stage algorithm proposed for MCAR). To solve this issue, we propose a so-called rescuing *a posteriori* algorithm that reassigns variables in samples that have lost all variables but respecting $P(M_{i,j} = 1|X^0_{i,j}, \lambda)$.

To do so, we define the binary indicator matrix $\xi \in \mathbb{R}^{(2^m-1)\times m}$, whose rows store all possible combinations of binary sequences of length $m$, with the exception of all-ones, where $\xi_{k,j} = 1$ denotes that the corresponding variable $j$ is missing in the combination $k$. For a given sample $i$, each variable has a probability of becoming missing equal to $p_{i,j} = P(M_{i,j} = 1|X^0_{i,j}, \lambda)$. We can define the pseudo-likelihood $\mathcal{L}$ of a pattern $\xi_{k,} := \{\xi_{k,1}, \ldots, \xi_{k,m}\}$ in sample $i$ as follows:

$$\mathcal{L}_i(\xi_{k,}) = \prod_{j=1}^{m} p_{i,j}^{\xi_{k,j}} (1 - p_{i,j})^{1-\xi_{k,j}}$$

We can normalize $\mathcal{L}_i(\xi_{k,})$ to obtain a vector of probabilities for each missing value pattern in sample $i$ (but excluding the case where all variables are missing):

$$P_i(\xi_{k,}) = \frac{\mathcal{L}_i(\xi_{k,})}{\sum_{k=1}^{2^m-1} \mathcal{L}_i(\xi_{k,})}$$

We can thus reassign samples with all missing values to another pattern by drawing one of the $2^m - 1$ patterns with probabilities $P_i(\xi_{k,})$. In any case, though we could have used this approach as the general procedure to generate missing values avoiding the case where all variables are missing, since there are exponential possibilities, this rescuing process is highly inefficient and is not recommended as a general generation approach. Therefore, we do not apply this procedure for the MCAR case. Strictly speaking, the all-variables-missing (AVM) issue is only a problem at high $p_{\text{NA}}$ (extreme missingness). The average number of expected samples suffering from AVM can be computed as

$$\mathbb{E}[\#\text{samples AVM}|p_{\text{NA}}] = n \cdot (p_{\text{NA}})^m$$

For example, see the table below:

Table S1: $\mathbb{E}[\#\text{samples AVM}|p_{\text{NA}}]$ as a function of $n$, $m$ and $p_{\text{NA}}$

| $n$ | $m$ | $p_{\text{NA}}$ | $\mathbb{E}[\#\text{samples AVM}|p_{\text{NA}}]$ |
|---|---|---|---|
| 100 | 20 | 0.5 | $9.53 \cdot 10^{-5}$ |
| 100 | 50 | 0.5 | $8.88 \cdot 10^{-14}$ |
| 100 | 20 | 0.9 | 12.16 |
| 100 | 50 | 0.9 | 0.52 |
| 100 | 20 | 0.95 | 35.85 |
| 100 | 50 | 0.95 | 7.69 |

##### 6.1.3 MAR

In our MAR model, for each variable $j \in \{1, \dots, m\}$, we draw a $t_j \neq j$ from the set $\{1, 2, \dots, m\} \setminus \{j\}$ with equal probabilities $\frac{1}{m-1}$. The probability of variable $j$ becoming missing in sample $i \in \{1, \dots, n\}$ is then defined as

$$P(M_{i,j} = 1|\phi_j, X^0_{i,\forall k \neq j}, X^0_{i,j}, W_{i,}) = P(M_{i,j} = 1|X^0_{i,t_j}, \lambda) \tag{44}$$

where $\lambda \in \mathbb{R}$ is a parameter. This means that $X^0_{i,j}$ may become missing depending on another variable of that same individual but not $X^0_{i,j}$ itself, unlike MNAR.

Like for MNAR, we now propose the following link functions:

$$\text{Left (L)} \qquad P(M_{i,j} = 1|X^0_{i,t_j}, \lambda) = \frac{1}{1+e^{-(X^0_{i,t_j}+\lambda)}}$$

$$\text{Right (R)} \qquad P(M_{i,j} = 1|X^0_{i,t_j}, \lambda) = \frac{1}{1+e^{-(-X^0_{i,t_j}+\lambda)}}$$

$$\text{Left-Right (LR)} \qquad P(M_{i,j} = 1|X^0_{i,t_j}, \lambda) = \frac{1}{1+e^{-(|X^0_{i,t_j}|+\lambda)}}$$

$$\text{Center (C)} \qquad P(M_{i,j} = 1|X^0_{i,t_j}, \lambda) = 1 - \frac{1}{1+e^{-(|X^0_{i,t_j}|+\lambda)}}$$

Again, we include a constant $\lambda$ to tune the probabilities so that $p_{\text{NA}}$ is respected. To tune $\lambda$, we define:

$$g(\lambda) = \frac{1}{m \cdot n} \sum_{i=1}^{n} \sum_{j=1}^{m} P(M_{i,j} = 1|X^0_{i,t_j}, \lambda) - p_{\text{NA}}$$

Like in MNAR, we choose $\lambda$ so that $g(\lambda)$ is as close to zero as possible. Finally, for a total of $n \times m$ times and per position $i, j$ we draw a binomial variable with probability $P(M_{i,j} = 1|X^0_{i,t_j}, \lambda)$ to fill matrix $M_{i,j}$. We also incorporate the same *a posteriori* rescuing mechanism as in MNAR but using the matrix $X^0_{i,t_j}$ instead of $X^0_{i,j}$.

#### 6.2 ADEMP simulation report

##### 6.2.1 Aims

The global objective is to benchmark aps-lm against unconditional mean imputation and MICE multiple imputation under the prediction paradigm with varying sample size, covariance structure of predictors, linear model goodness-of-fit (GOF) or missing value type, to compare aps-lm and aps-ridge and to derive the coverage of the calculated confidence and prediction intervals. There are four rounds of simulations: Round 1 aims to simply test aps-lm under a very wide range of conditions and round 2 focuses on how performance is impacted by different types of missing values. In round 3 the performance of aps-ridge and aps-lm is compared whereas round 4 investigates the coverage of derived confidence and prediction intervals of aps-lm.

###### 6.2.2 General parameters

The definition of the different parameters is listed in Table S2.

Table S2: General parameters - definitions

| **Param** | **Interpretation** | | |
|---|---|---|---|
| $m$ | Number of variables | | |
| $n$ | Number of samples | | |
| $\theta$ | Vector of linear coefficients | | |
| $R^2$ | Goodness-of-fit | | |
| $\mu$ | Vector of means of predictors | | |
| $\sigma^2$ | Vector of variances of predictors | | |
| $\Sigma$ | Covariance matrix of predictors | | |
| $p_H$ | Ref-App split proportion (hold-out) | | |
| $\text{type}_{\text{NA}}$ | Type of missing value | MCAR | |
| | | MNAR | subtypes = {L, R, LR, C} |
| | | MAR | subtypes = {L, R, LR, C} |
| $N_{\text{iter}}$ | Number of Monte-Carlo iterations | | |

To ensure that the simulations are systematic, we decided to fix certain parameters; this way, we avoid making parametric assumptions on potential generation mechanisms that could bias the conclusions of the outcomes at least for those parameters. However, as a result, making a specific choice may seem arbitrary. We selected the parameters to be of several scales and including positive and negative values when possible.

To begin with, we made a compromise in the number of variables; i.e. large enough to observe high-order interactions but small enough to achieve a relatively high number of Monte-Carlo iterations:

$$m = 10; \quad N_{\text{iter}} = 50$$

As linear coefficient vector, we chose:

$$\theta = (0.70, -0.40, 2.00, 0.10, 0.50, -2.00, 0.50, 3.00, 0.80, -0.90, 0.01) \tag{45}$$

For the generation of the design matrix $X$, we employed the following mean and variance vectors of the predictors:

$$\mu = (2.000, 1.000, 0.200, -2.000, 3.000, 0.100, 1.400, -2.000, 3.000, 0.001) \tag{46}$$

$$\sigma^2 = (1.42, 0.82, 1.22, 2.22, 1.67, 2, 0.71, 1.8, 0.73, 1.36)$$

As for covariance matrices of the predictors, we chose four matrices representing independence, weak, strong dependence and ultra-strong dependence.

$$\Sigma_{indep} = \begin{pmatrix} 1.42 & 0 & 0 & 0 & 0 & 0 & 0 & 0 & 0 & 0 \\ 0 & 0.82 & 0 & 0 & 0 & 0 & 0 & 0 & 0 & 0 \\ 0 & 0 & 1.22 & 0 & 0 & 0 & 0 & 0 & 0 & 0 \\ 0 & 0 & 0 & 2.22 & 0 & 0 & 0 & 0 & 0 & 0 \\ 0 & 0 & 0 & 0 & 1.67 & 0 & 0 & 0 & 0 & 0 \\ 0 & 0 & 0 & 0 & 0 & 2.00 & 0 & 0 & 0 & 0 \\ 0 & 0 & 0 & 0 & 0 & 0 & 0.71 & 0 & 0 & 0 \\ 0 & 0 & 0 & 0 & 0 & 0 & 0 & 1.80 & 0 & 0 \\ 0 & 0 & 0 & 0 & 0 & 0 & 0 & 0 & 0.73 & 0 \\ 0 & 0 & 0 & 0 & 0 & 0 & 0 & 0 & 0 & 1.36 \end{pmatrix}$$

$$\Sigma_{wd} = \begin{pmatrix} 1.42 & -0.16 & 0.32 & 0.49 & 0.3 & 0.11 & 0.24 & -0.64 & 0.02 & -0.8 \\ -0.16 & 0.82 & 0.28 & -0.82 & 0.3 & -0.39 & 0.23 & -0.09 & -0.24 & -0.14 \\ 0.32 & 0.28 & 1.22 & -0.36 & 0.78 & 0.19 & 0.46 & -0.34 & -0.2 & 0.28 \\ 0.49 & -0.82 & -0.36 & 2.22 & 0.27 & 0.75 & 0.33 & 0.03 & 0.31 & -0.26 \\ 0.3 & 0.3 & 0.78 & 0.27 & 1.67 & 0.84 & 0.44 & 0.42 & -0.17 & -0.16 \\ 0.11 & -0.39 & 0.19 & 0.75 & 0.84 & 2.00 & -0.03 & 1.14 & 0.03 & -0.48 \\ 0.24 & 0.23 & 0.46 & 0.33 & 0.44 & -0.03 & 0.71 & -0.14 & -0.06 & -0.05 \\ -0.64 & -0.09 & -0.34 & 0.03 & 0.42 & 1.14 & -0.14 & 1.80 & -0.030 & -0.31 \\ 0.02 & -0.24 & -0.2 & 0.31 & -0.17 & 0.03 & -0.06 & -0.03 & 0.73 & 0.09 \\ -0.8 & -0.14 & 0.28 & -0.26 & -0.16 & -0.48 & -0.05 & -0.31 & 0.09 & 1.36 \end{pmatrix}$$

$$\Sigma_{sd} = \begin{pmatrix} 1.42 & -0.06 & 0.58 & 0.36 & 0.92 & -0.12 & 0.51 & 0.61 & -0.02 & -0.02 \\ -0.06 & 0.82 & -0.15 & -0.19 & -0.09 & 0.58 & -0.13 & -0.14 & 0.01 & -0.09 \\ 0.58 & -0.15 & 1.22 & 0.84 & 1.03 & -0.58 & 0.79 & 1.10 & -0.05 & 0.11 \\ 0.36 & -0.19 & 0.84 & 2.22 & 0.99 & -0.31 & 0.73 & 1.02 & 0.01 & 0.28 \\ 0.92 & -0.09 & 1.03 & 0.99 & 1.67 & -0.21 & 0.96 & 1.17 & -0.15 & 0.15 \\ -0.12 & 0.58 & -0.58 & -0.31 & -0.21 & 2.00 & -0.41 & -0.42 & -0.03 & -0.19 \\ 0.51 & -0.13 & 0.79 & 0.73 & 0.96 & -0.41 & 0.71 & 0.91 & -0.11 & 0.13 \\ 0.61 & -0.14 & 1.10 & 1.02 & 1.17 & -0.42 & 0.91 & 1.80 & -0.17 & 0.06 \\ -0.02 & 0.01 & -0.05 & 0.01 & -0.15 & -0.03 & -0.11 & -0.17 & 0.73 & -0.13 \\ -0.02 & -0.09 & 0.11 & 0.28 & 0.15 & -0.19 & 0.13 & 0.06 & -0.13 & 1.36 \end{pmatrix}$$

$$\Sigma_{usd} = \begin{pmatrix} 1.42 & -0.8 & -1.05 & -0.85 & -1.08 & -0.9 & -0.78 & 1.43 & 0.75 & -1.18 \\ -0.8 & 0.82 & 0.51 & 0.72 & 0.65 & 0.52 & 0.54 & -0.94 & -0.49 & 0.81 \\ -1.05 & 0.51 & 1.22 & 1.19 & 1.02 & 0.86 & 0.82 & -1.28 & -0.78 & 0.91 \\ -0.85 & 0.72 & 1.19 & 2.22 & 1.21 & 0.7 & 1.06 & -1.31 & -0.86 & 0.94 \\ -1.08 & 0.65 & 1.02 & 1.21 & 1.67 & 1.05 & 0.74 & -1.41 & -0.61 & 1.3 \\ -0.9 & 0.52 & 0.86 & 0.7 & 1.05 & 2 & 0.56 & -1.04 & -0.6 & 1.04 \\ -0.78 & 0.54 & 0.82 & 1.06 & 0.74 & 0.56 & 0.71 & -1.03 & -0.67 & 0.76 \\ 1.43 & -0.94 & -1.28 & -1.31 & -1.41 & -1.04 & -1.03 & 1.8 & 1.01 & -1.44 \\ 0.75 & -0.49 & -0.78 & -0.86 & -0.61 & -0.6 & -0.67 & 1.01 & 0.73 & -0.69 \\ -1.18 & 0.81 & 0.91 & 0.94 & 1.3 & 1.04 & 0.76 & -1.44 & -0.69 & 1.36 \end{pmatrix}$$

where subindices *indep*, *wd*, *sd* and *usd* stand for independence, weak, strong and ultra-strong dependence, respectively. Looking at the entries in the covariance matrices alone, it is hard to interpret the corresponding degrees of dependence. To attempt to quantify it, we can first standardize $\Sigma$ as in the Pearson correlation matrix $C$ with entries $C_{i,j} = \frac{\Sigma_{i,j}}{\sqrt{\Sigma_{i,i}} \cdot \sqrt{\Sigma_{j,j}}}$ (displayed as Figure S1). Then, we can compute the eigenvalues $\lambda_j$ of $C$ and normalize them as in $\nu_j = \lambda_j / (\sum_{k=1}^{m} \lambda_k)$, where $\sum_{j=1}^{m} \nu_j = 1$; $\nu_j$ can be interpreted as the proportion of variance explained by the principal component $j$. Independence is reflected by equal $\nu_j$ (maximum entropy) whilst high levels of dependence deviate from equality; in other words, under independence, $\forall j : \nu_j = 1/m$. We can thus quantify dependence with either normalized entropy $S$ or normalized Gini-Simpson index $G$:

$$S = -\sum_{j=1}^{m} \frac{\log(\nu_j) \cdot \nu_j}{\log(m)}; \quad G = \frac{m}{m-1} \cdot \left[1 - \sum_{j=1}^{m} \nu_j^2\right]$$

For our chosen covariance matrices, the following $S$ and $G$ are obtained:

Table S3: Normalized Entropy and Gini-Simpson index for the chosen covariance matrices

| Param | Normalized Entropy | Normalized Gini-Simpson index |
|---|---|---|
| $\Sigma_{indep}$ | 1.000 | 1.000 |
| $\Sigma_{wd}$ | 0.828 | 0.920 |
| $\Sigma_{sd}$ | 0.787 | 0.853 |
| $\Sigma_{usd}$ | 0.449 | 0.501 |

Finally, we fixed the proportion of sample sizes of the reference-to-application datasets to the typical hold-out value of 80 %:

$$p_H = 0.8$$

###### 6.2.3 Generation mechanisms

The design matrix $X$ is drawn from a multivariate normal distribution (employing `MASS::mvrnorm`) with $\mu$ and $\Sigma$ as described in the previous section:

$$X \sim \mathcal{N}(\mu, \Sigma)$$

The dependent variable is a linear combination of the predictors (with intercept), plus Gaussian white noise:

$$y = (1|X) \cdot \theta + \epsilon; \quad \epsilon \sim \mathcal{N}(\vec{0}_n, \sigma_\epsilon^2 I_n) \tag{47}$$

The GOF of the model, $R^2$, is regulated by tuning the scale of the white noise, $\sigma_\epsilon^2$:

$$\tilde{y} = (1|X) \cdot \theta$$

with which we derive:

$$R^2 = \frac{\sigma_{\tilde{y}}^2}{\sigma_{\tilde{y}}^2 + \sigma_\epsilon^2}; \quad \sigma_\epsilon^2 = \left(\frac{1 - R^2}{R^2}\right) \cdot \sigma_{\tilde{y}}^2 \tag{48}$$

#### 6.3 Round 1: Testing aps-lm under a wide range of conditions

A total of 21 combinations of covariance matrices and missing value types were used for simulation (see Table 1, main manuscript), in which we explored the following range of parameters.

Table S4: Round 1 - range of parameters

| Param | Interpretation | Range | |
|---|---|---|---|
| $n$ | Number of samples | $n \in \{28, 56, 84, 112, 140, 168, 196, 224, 252, 280\}$ | |
| $R^2$ | Proportion of variance explained | $R^2 \in \{0.2, 0.45, 0.70, 0.95\}$ | |
| $p_{\text{NA}}$ | Proportion of missing values | $p_{\text{NA}} \in \{0.05, 0.10, 0.20, 0.30, 0.50, 0.70\}$ | |
| $\Sigma$ | Covariance matrix | $\Sigma = \{\Sigma_{indep}, \Sigma_{wd}, \Sigma_{sd}\}$ | |
| $\text{type}_{\text{NA}}$ | Type of missing value | MCAR | |
| | | MNAR | subtypes = {L, R, LR, C} |
| | | MAR | subtypes = {L, R, LR, C} |

All tabulated values for $n$, $R^2$ and $p_{\text{NA}}$ were used for simulation, resulting in $10 \times 4 \times 6 = 240$ scenarios for each of the 21 combinations. For a fixed covariance matrix and missing value type, for each $n$, $R^2$ and $p_{\text{NA}}$, the following steps were performed a total of $N_{\text{iter}}$ times: i) Generate $X$, $y$ as described above; ii) Split $X$ and $y$ into $X_{\text{ref}}$ (reference), $X_{\text{app}}$ (application) and $y_{\text{ref}}$, $y_{\text{app}}$ respecting proportion $p_H$; iii) Inject missing values in $X_{\text{app}}$ following a $\text{type}_{\text{NA}}$ scheme; iv). Perform benchmark:

a) Full: Train a linear model on $X_{\text{ref}}$ and test on $X_{\text{app}}$ using the original application dataset without missing values, $X^0_{\text{app}}$. b) Mean: Perform mean imputation on $X_{\text{app}}$: replacing missing values by means from the application dataset. Train a linear model on $X_{\text{ref}}$ and apply to $X_{\text{app,imputed}}$. It is to be noted that if a variable is missing for all samples, then the mean of the given variable in the application set is undefined and hence, mean imputation is not possible. c) MICE: Perform MICE imputation on $X_{\text{app}}$. Train a linear model on $X_{\text{ref}}$ and apply to $X_{\text{app,imputed}}$. It is to be noted that for non-convergence or if collinearity issues exist, MICE imputation procedure fails; the second behavior can be overridden by setting argument `remove.collinear` to `FALSE`, which we did throughout this study. d) aps-lm: Train an aps-lm model on $X_{\text{ref}}$ (find parameters $R$ and $\hat{\theta}_{\text{ols}}$) and apply to $X_{\text{app}}$. As for performance measure in round 1, we exclusively used the Pearson correlation squared for convenience. The bias measure is examined in round 2. For all conditions and for each Monte-Carlo iteration, indexed by the index $r$, we compute the Pearson correlation between the predicted, $\hat{y}^{\text{app}}_r \in \mathbb{R}^{n_{\text{app}}}$, and the real outcome vector $y^{\text{app}}_r \in \mathbb{R}^{n_{\text{app}}}$ as

$$\hat{\rho}_r(y^{\text{app}}_r, \hat{y}^{\text{app}}_r) = \frac{\widehat{\text{Cov}}(y^{\text{app}}_r, \hat{y}^{\text{app}}_r)}{\hat{\sigma}_{y^{\text{app}}_r} \cdot \hat{\sigma}_{\hat{y}^{\text{app}}_r}}$$

where $\widehat{\text{Cov}}(X, Y)$ is the covariance estimator. For visualization purposes, we plot $\hat{\rho}^2_r(y^{\text{app}}_r, \hat{y}^{\text{app}}_r)$

as a function of $n$, $R^2$, and $p_{\mathrm{NA}}$, the relationship of which is visualized as a locally weighted scatterplot smoothing (loess) curve presenting four differently colored curves for the model categories. This was possible with the help of the ggplot2 R-package (Wickham (2016)).

#### 6.4 Round 2: Simultaneously assessing all types of missing values

Having explored the effects of $R^2$ and $n$ in the previous round, we now fix these parameters (see Table S5).

Table S5: Round 2 - range of parameters

| **Param** | **Interpretation** | **Range** | |
|---|---|---|---|
| $n$ | Number of samples | $n = 200$ | |
| $R^2$ | Proportion of variance explained | $R^2 = 0.95$ | |
| $p_{\mathrm{NA}}$ | Proportion of missing values | $p_{\mathrm{NA}} \in \{0.05, 0.10, 0.20, 0.30, 0.50, 0.70\}$ | |
| $\Sigma$ | Covariance matrix | $\Sigma = \{\Sigma_{indep}, \Sigma_{wd}, \Sigma_{sd}, \Sigma_{usd}\}$ | |
| $\mathrm{type}_{\mathrm{NA}}$ | All missing value types in parallel | MCAR | |
| | | MNAR | subtypes = {L, R, LR, C} |
| | | MAR | subtypes = {L, R, LR, C} |

For these simulations, we used two values of $\theta$. In addition to the original $\theta$ we modified this from:

$$\theta_1^{\top} = (0.70, -0.40, 2.00, 0.10, 0.50, -2.00, 0.50, 3.00, 0.80, -0.90, 0.01)$$

to:

$$\theta_2^{\top} = (0.70, \mathbf{0.40}, \mathbf{10.0}, 0.10, 0.50, \mathbf{2.00}, 0.50, 3.00, 0.80, -0.90, 0.01)$$

We apply the different missing value schemes on the same datasets (unlike Round 1 in which different missing value types were evaluated independently). Given a covariance matrix, for each $p_{\mathrm{NA}}$, the following steps are performed a total of $N_{\mathrm{iter}}$ times: i) Generate $X$, $y$; ii) Split $X$ and $y$ into $X_{\mathrm{ref}}$ (reference), $X_{\mathrm{app}}$ (application) and $y_{\mathrm{ref}}$, $y_{\mathrm{app}}$ based on $p_H$; iii) Inject missing values in $X_{\mathrm{app}}$ following all types of missing value schemes on the same dataset in parallel; iv) Perform benchmark for each missing value scheme and for each imputation each method. For all conditions and for each Monte-Carlo iteration, indexed by an index $r$, we compute:

$$\hat{\rho}_r(y_r^{\mathrm{app}}, \hat{y}_r^{\mathrm{app}}) = \frac{\widehat{\mathrm{Cov}}(y_r^{\mathrm{app}}, \hat{y}_r^{\mathrm{app}})}{\hat{\sigma}_{y_r^{\mathrm{app}}} \cdot \hat{\sigma}_{\hat{y}_r^{\mathrm{app}}}}; \quad \mathrm{Bias}_r(y_r^{\mathrm{app}}, \hat{y}_r^{\mathrm{app}}) = \frac{1}{n_{\mathrm{app}}} \sum_{i=1}^{n_{\mathrm{app}}} (y_{r,i}^{\mathrm{app}} - \hat{y}_{r,i}^{\mathrm{app}})$$

For $\theta_1$ we also compute the mean squared error (MSE) as:

$$\text{MSE}_r(y_r^{\text{app}}, \hat{y}_r^{\text{app}}) = \frac{1}{n_{\text{app}}} \sum_{i=1}^{n_{\text{app}}} (y_{r,i}^{\text{app}} - \hat{y}_{r,i}^{\text{app}})^2$$

and the concordance correlation coefficient using the function `DescTools::CCC` (Signorell (2023)), which estimates it as follows:

$$\hat{\rho}_r^{\text{conc}}(y_r^{\text{app}}, \hat{y}_r^{\text{app}}) = \frac{2 \cdot \hat{\rho}_r(y_r^{\text{app}}, \hat{y}_r^{\text{app}}) \cdot \hat{\sigma}_{y_r^{\text{app}}} \cdot \hat{\sigma}_{\hat{y}_r^{\text{app}}}}{\hat{\sigma}^2_{y_r^{\text{app}}} + \hat{\sigma}^2_{\hat{y}_r^{\text{app}}} + \frac{n}{n-1} \cdot (\hat{\mu}_{y_r^{\text{app}}} - \hat{\mu}_{\hat{y}_r^{\text{app}}})^2}$$

#### 6.5 Round 3: aps-lm vs aps-ridge

The set-up in round 3 is mostly equivalent to round 2 (see section 6.4), only that instead of benchmarking aps-lm against imputation approaches (mean imputation and MICE imputation), aps-lm is benchmarked against of aps-ridge. Only $\theta_1$ is used in this round of simulation. aps-ridge is equivalent to the version described in section 4, where $\lambda$ is estimated (once per iteration) by 5-fold cross-validation in the complete reference dataset. To evaluate performance throughout this round of simulations, we employed the MSE metric instead of squared Pearson correlation since the R-package employed for cross-validation (`glmnet`, Friedman et al. (2010)), by default, uses this same metric to optimize the regularization parameter $\lambda$. Thus a fair comparison to ridge-based approaches is only possible if we compare the MSE.

Table S6: Round 4 - range of parameters

| Param | Interpretation | Range |
|---|---|---|
| $n$ | Number of samples | $n = \{50, 100, 200\}$ |
| $R^2$ | Proportion of variance explained | $R^2 = \{0.25, 0.5, 0.95\}$ |
| $\Omega_0$ | Missing variable set 0 | $\Omega_0 = \emptyset$ |
| $\Omega_1$ | Missing variable set 1 | $\Omega_1 = \{2\}$ |
| $\Omega_2$ | Missing variable set 2 | $\Omega_2 = \{2, 4, 7\}$ |
| $\Omega_3$ | Missing variable set 3 | $\Omega_3 = \{1, 2, 3, 4, 8, 9\}$ |
| $\Sigma$ | Covariance matrix | $\Sigma = \{\Sigma_{indep}, \Sigma_{wd}, \Sigma_{sd}\}$ |
| $N_{\text{cov}}$ | Number of samples used to estimate coverage | $N_{\text{cov}} = 1000$ |
| $N_{\text{rep}}$ | Number of times coverage estimation is repeated | $N_{\text{rep}} = 20$ |

#### 6.6 Round 4: Coverage of prediction and confidence intervals

In this round of simulations, we want to investigate whether the prediction and confidence levels of the estimated prediction and confidence intervals of the aps-submodel are valid, i.e., the predefined coverage is obtained in the simulations. The scenarios we investigated are shown in Table S6. For all scenarios $\mu$ and $\theta$ will be chosen as before in 46 and 45. In contrast to the other rounds of simulations, we have here to fix $\Omega$ and not to simulate it by a missing value mechanism because we have to simulate from the same submodel several times to determine the coverage. The simulations are performed in the following steps:

(i) Simulation of a reference vector of predictors $x^0$
We want to calculate the prediction interval for a specific vector of predictors which has to be simulated from the full model without missing values. We thus simulate $x^0$ from $\mathcal{N}(\mu, \Sigma)$.

(ii) Simulation of the full model
We then have to simulate the full model, see equation 47

$$y = (1|X) \cdot \theta + \epsilon; \quad \epsilon \sim \mathcal{N}(\vec{0}_n, \sigma_\epsilon^2 I_n)$$

We do this by simulating $x_1$, ..., $x_n$ independently from $X_i \sim \mathcal{N}(\mu, \Sigma)$ as before. $x_1$, ..., $x_n$ then render the design matrix $X$.
Again, $\sigma_\epsilon^2$ is determined by $R^2$ (see equation 48) such that we can simulate $y_1$, ..., $y_n$ from the full model equation.

(iii) Estimation of parameters
Now we estimate the parameters $R$ and $\hat{\theta}_{ols}$ from the simulated full model and compute $y^\top y$.

(iv) Calculation of the confidence and prediction intervals of the submodel
The submodel for which we want to calculate the prediction interval is

$$y = (1|X) \cdot I_{m+1}^{(,\Omega)} \cdot \theta_\Omega + \epsilon_\Omega; \quad \epsilon_\Omega \sim \mathcal{N}(\vec{0}_n, \sigma_{\epsilon,\Omega}^2 I_n)\ .$$

The estimated expected outcome $\hat{y}_\Omega^0$ of the submodel given $x^0$ is (compare equation (12) in the main manuscript):

$$\hat{y}_\Omega^0 = x_\Omega^0 \cdot \hat{\theta}_\Omega$$

with $\hat{\theta}_\Omega$ calculated with the parameters from (iii) (see equation (10) in the main manuscript). Here, $x_\Omega^0$ is the same as $x^0$ but the values corresponding to $\Omega$ are omitted. We can now compute the confidence and prediction intervals of the submodel for the given $x^0$ from equations (18) and (19) in the main manuscript.

(v) Simulation from the submodel
We have now to simulate from the submodel given $x^0$ to see if the predefined coverage is reached. We thus have to simulate from the equation

$$y = x_\Omega^0 \cdot \theta_\Omega + \epsilon_\Omega; \quad \epsilon_\Omega \sim \mathcal{N}(0, \sigma_{\epsilon,\Omega}^2)\ . \tag{49}$$

Note that here $\theta_\Omega$ and $\sigma^2_{\epsilon,\Omega}$ are the true (not estimated) values. $\theta_\Omega$ can be derived from equation (10) in the main manuscript for the true values:

$$\theta_\Omega = \left(R_{\text{true}} \cdot I^{(,\Omega)}_{m+1}\right)^\dagger \cdot R_{\text{true}} \cdot \theta$$

Again, we have here $R_{\text{true}}$ instead of $R$. Whereas $R$ is estimated, $R_{\text{true}}$ is the true value which can be determined from

$$R_{\text{true}} = \text{chol}(n \cdot \Sigma + n \cdot \mu \cdot \mu^\top) \ .$$

The final parameter needed for the simulation from equation 49, is $\sigma^2_{\epsilon,\Omega}$. For this, we first define

$$\tilde{y_\Omega} = (1|X) \cdot I^{(,\Omega)}_{m+1} \cdot \theta_\Omega \ .$$

Then similar to equation 48 we have

$$\sigma^2_{\epsilon,\Omega} = \left(\frac{1 - R^2_\Omega}{R^2_\Omega}\right) \cdot \sigma^2_{\tilde{y}_\Omega} \ .$$

$\sigma^2_{\tilde{y}_\Omega}$ can be obtained from

$$\sigma^2_{\tilde{y}_\Omega} = \theta^\top_\Omega \cdot I^{(\Omega,)}_{m+1} \cdot \Sigma \cdot I^{(,\Omega)}_{m+1} \cdot \theta_\Omega \ .$$

Finally, the following equation is used for the determination of $R^2_\Omega$

$$R^2_\Omega = \frac{\sigma^2_{\tilde{y}_\Omega}}{\sigma^2_y} = \frac{\sigma^2_{\tilde{y}_\Omega}}{\theta^\top \cdot \Sigma \cdot \theta + \sigma^2_\epsilon} \ .$$

We can now simulate from equation 49 and will obtain a value $y^0$.

For each $x^0$, steps (ii) to (v) are repeated $N_{\text{cov}} = 1000$ times. Then it is calculated how often $y^0$ is in the calculated prediction interval to determine the coverage. Similarly, the coverage of the calculated confidence interval can be determined by noting how often the mean $\tilde{y}^0_\Omega := x^0_\Omega \cdot \theta_\Omega$ is in the calculated confidence interval. The generation of $x^0$ in step (i) is repeated $N_{\text{rep}} = 20$ times such that we finally have 20 values of the coverage of prediction and confidence interval each.

### 7 Epigenetic aging clocks

We tested the aps-lm method on the application of epigenetic aging clocks. To do so, we employed the DNA methylation microarray data freely available at EWAS Data Hub (Xiong et al. (2020)), compiled and curated from Gene Expression Omnibus (GEO) (Barrett et al. (2012)). This dataset targets 85,512 cytosines and consists of 8,374 samples from a wide range of tissues and ages. The EWAS datahub epigenetic aging dataset (11.73 GB) can be downloaded with the following bash command:

```
wget https://download.cncb.ac.cn/ewas/datahub/download/age_methylation_v1.zip
```

We obtained the methylation sites of Horvath's (Horvath (2013)) epigenetic aging clock via the wateRmelon R-package (Pidsley et al. (2013)), using the command `data(coef)`. To extract the EWAS datahub data on the sites, we ran the following bash command:

```
grep -f horvath_CpGs.txt age_methylation_v1.txt > horvath_age_methylation_v1.txt
```

The data was then split in a 9:1 ratio into $X_{\text{ref}}, y_{\text{ref}}$ (n = 7,536) and $X_{\text{app}}, y_{\text{app}}$ (n = 838) where $X$ refers to the methylation values and $y$ to the chronological age. Based on $X_{\text{ref}}, y_{\text{ref}}$, we had four different set-ups:

- (*Scenario 1*): mean imputation on $X_{\text{ref}}$ followed by a linear model,
- (*Scenario 2*): MICE imputation on $X_{\text{ref}}$ followed by a linear model,
- (*Scenario 3*): mean imputation on $X_{\text{ref}}$ followed by a ridge model, and
- (*Scenario 4*): MICE imputation on $X_{\text{ref}}$ followed by a ridge model.

For MICE imputation on $X_{\text{ref}}$, we generated five different imputed versions, denoted as $X_{\text{ref}}^{\text{MICE},r}$, $r \in \{1, \ldots, 5\}$ (generating several imputed datasets is standard practice when applying MICE). For implementing MICE on the large dataset of the EWAS Datahub, we used the parallelized version `micemd::mice`.

We then evaluated the performance of aps-lm and ridge-lm in comparison to mean imputation on $X_{\text{app}}$ by assuming $y_{\text{app}}$ to be unknown. For the MICE imputation in scenarios 2 and 4, we estimated linear coefficients $\hat{\theta}_{\text{ols}}^{\text{MICE},r}$, mean $\hat{\mu}^{\text{MICE},r}$ and covariance matrix $\hat{\Sigma}^{\text{MICE},r}$ for each imputed version $r \in \{1, \ldots, 5\}$ (see equations 2 and 13 in the main text). These estimates are pooled to obtain $\hat{\mu}^{\text{MICE}}$, $\hat{\Sigma}^{\text{MICE}}$ and $\hat{\theta}_{\text{ols}}^{\text{MICE}}$. For pooling means, given that all imputed datasets were of the same sample size, we simply averaged the means across imputed datasets; for pooling covariance matrices, we used `miceadds::micombine.cov`; for pooling linear coefficients we used `mice::pool`. We obtained $R^{\text{MICE}}$ from equation 15 (main manuscript):

$$R^{\text{MICE}} = \text{chol}((n-1) \cdot \hat{\Sigma}^{\text{MICE}} + n \cdot \hat{\mu}^{\text{MICE}} \cdot (\hat{\mu}^{\text{MICE}})^\top)$$

For the ridge regression in scenarios 3 and 4, parameters were derived as follows (MI refers to mean imputation):

$$R_\lambda^{\text{MI}} = \text{chol}\left((R^{\text{MI}})^\top \cdot R^{\text{MI}} + \lambda \cdot I_{m+1}^{(,\{1\})} \cdot I_{m+1}^{(\{1\},)}\right); \quad \hat{\theta}_{\text{ridge}}^{\text{MI}} = ((R_\lambda^{\text{MI}})^\top R_\lambda^{\text{MI}})^{-1} \cdot (R^{\text{MI}})^\top R^{\text{MI}} \cdot \hat{\theta}_{\text{ols}}^{\text{MI}}$$

and

$$R_\lambda^{\text{MICE}} = \text{chol}\left((R^{\text{MICE}})^\top \cdot R^{\text{MICE}} + \lambda \cdot I_{m+1}^{(,\{1\})} \cdot I_{m+1}^{(\{1\},)}\right); \quad \hat{\theta}_{\text{ridge}}^{\text{MICE}} = ((R_\lambda^{\text{MICE}})^\top R_\lambda^{\text{MICE}})^{-1} \cdot (R^{\text{MICE}})^\top R^{\text{MICE}} \cdot \hat{\theta}_{\text{ols}}^{\text{MICE}}$$

To find the regularization parameter $\lambda$, we employed 10-fold cross-validation via the efficient routines implemented in the glmnet R-package (Friedman et al. (2010)) on mean-imputed $X_{\text{ref}}$. In order to translate $\lambda$ from glmnet's (Friedman et al. (2010)) objective function to the standard ridge's cost function we applied:

$$\lambda = \frac{n \cdot \lambda_{\text{glmnet}}}{\sqrt{\left(\frac{(n-1)\cdot\hat{\sigma}_y^2}{n}\right)}}$$

This transformation is required since glmnet modifies the cost function, normalizes $\lambda$ by the sample size and scales the dependent variable. For all scenarios, the response variable $y$ (i.e. age) was transformed with the piecewise Horvath transformation (Horvath (2013)):

$$F(y,k) = t = \begin{cases} \log\left(\frac{y+1}{k+1}\right) & y \leq k \\ \\ \frac{y-k}{k+1} & y > k \end{cases}$$

The constant $k$ is interpreted as adult age ($k = 20$, according to Horvath (2013)) and is used to distinguish two types of dynamics: logarithmic at childhood and linear at adulthood. An inverse transform can return predictions to the original age scale:

$$y = F^{-1}(t,k) = \begin{cases} e^t \cdot (k+1) - 1 & t \leq 0 \\ \\ t \cdot (k+1) + k & t > 0 \end{cases}$$

To test how performance decreased by increasing levels of missing values beyond its background levels, we artificially injected missing values in $X_{\text{app}}$. For simplicity, we solely implemented the MCAR criterion. We employed the following parameters:

Table S7: Application: epigenetic clocks - range of parameters

| **Param** | **Interpretation** | **Range** |
|---|---|---|
| $p_{\text{NA}}$ | Proportion of missing values | $p_{\text{NA}} = p_0 + 0.05 \cdot e; \quad e \in [[0:18]]$ |
| Scenario | Scenario | {TrainMeanImp, TrainMICE, TrainMeanImp_L2, TrainMICE_L2} |
| Testing | Testing Configuration | {MeanImp, aps-lm} |
| $N_{\text{iter}}$ | Number of iterations per $p_{\text{NA}}$ | 10 |

where $p_0 = 0.04190432$ is the background level of missing values in the application dataset. We measured performance by MSE and the bias between real and predicted age (on the original rather than the transformed scale).
