## Supplementary Figures for "Adaptive predictor-set linear model: an imputation-free method for linear regression prediction on datasets with missing values"

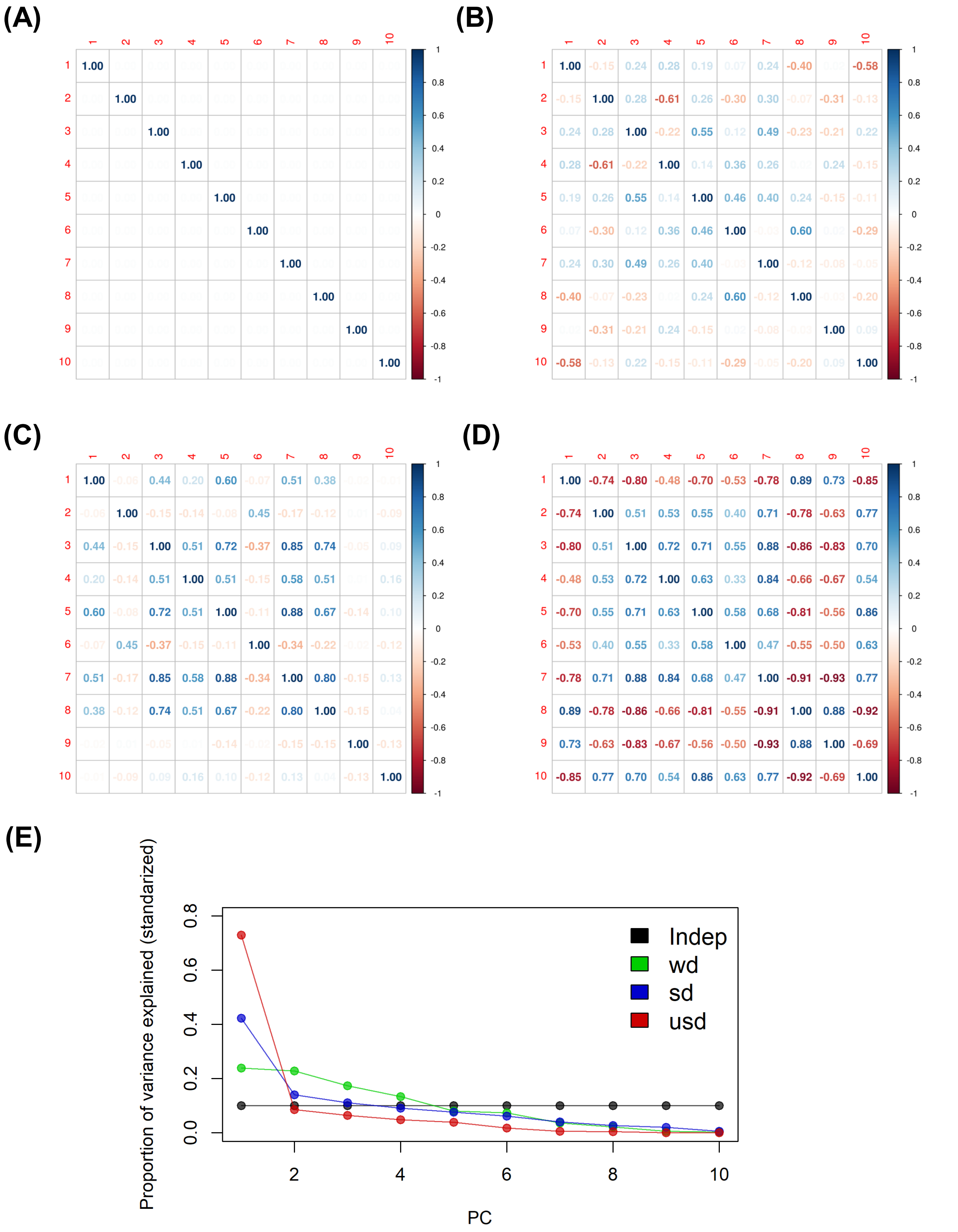


**Figure S1**. Correlation matrices corresponding to the covariances matrices employed: (A) independence, (B) weak, (C) strong and (D) ultra-strong dependence. (E) Proportion of variance explained computed from the eigenvalues of the correlation matrices.

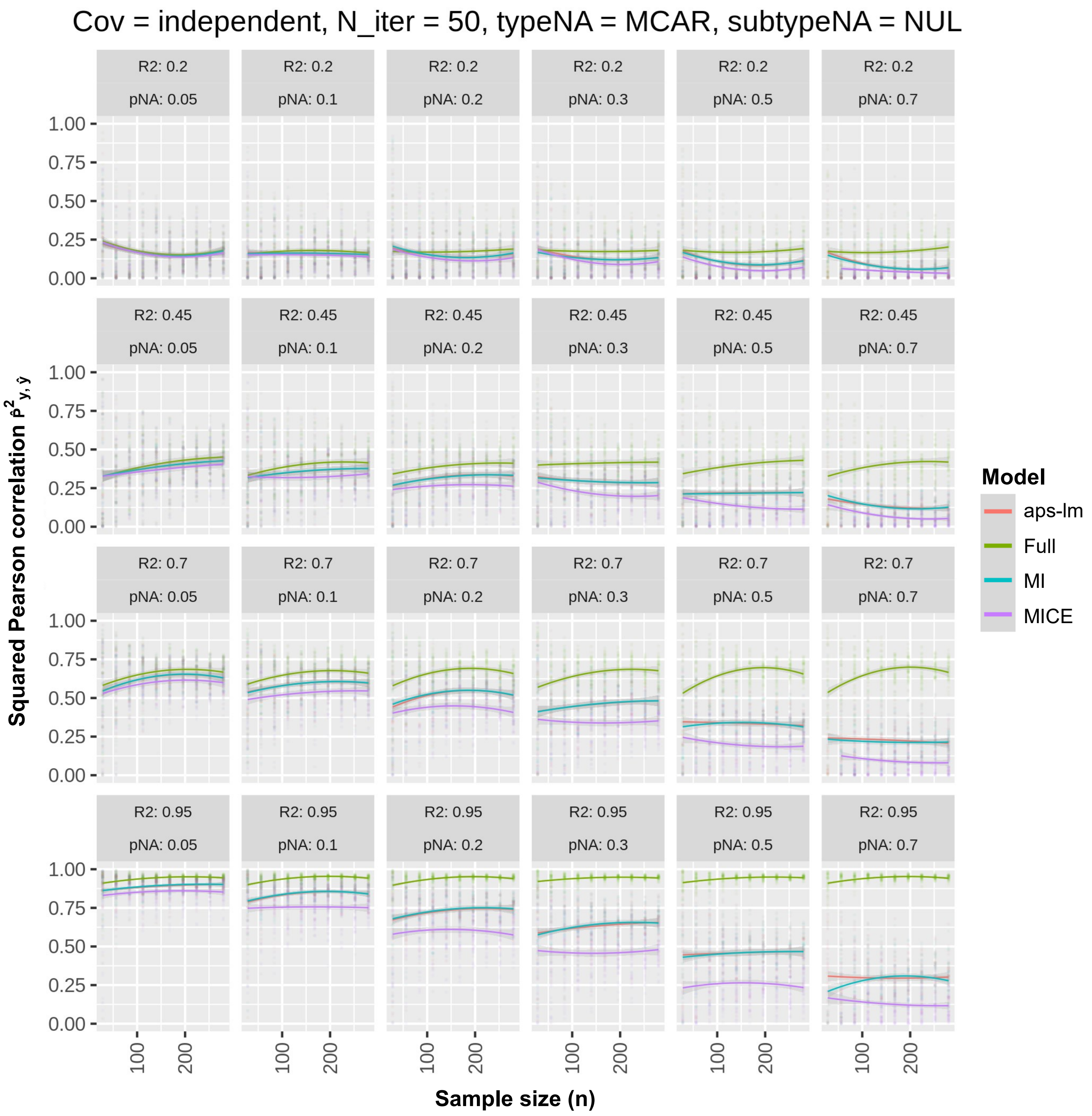


**Figure S2**. Round 1-Simulation 1: MCAR under independence. Missing values were injected on the application dataset following an MCAR scheme with variable proportion of missing values ($p_{NA}$). Squared Pearson correlation between real and predicted dependent variable is plotted at varying sample size (n), $R^2$ and $p_{NA}$ for: i) the estimated linear regression model of stage (i) applied on the application dataset $X_{app}$ without missing data (full), applied on $X_{app}$ with missing data after performing ii) unconditional mean (MI) or iii) MICE imputation (MICE) and iv) aps-lm without applying any imputation. Simulation results are visualized as jitter plots while overall tendencies in the mean are represented as LOESS curves.

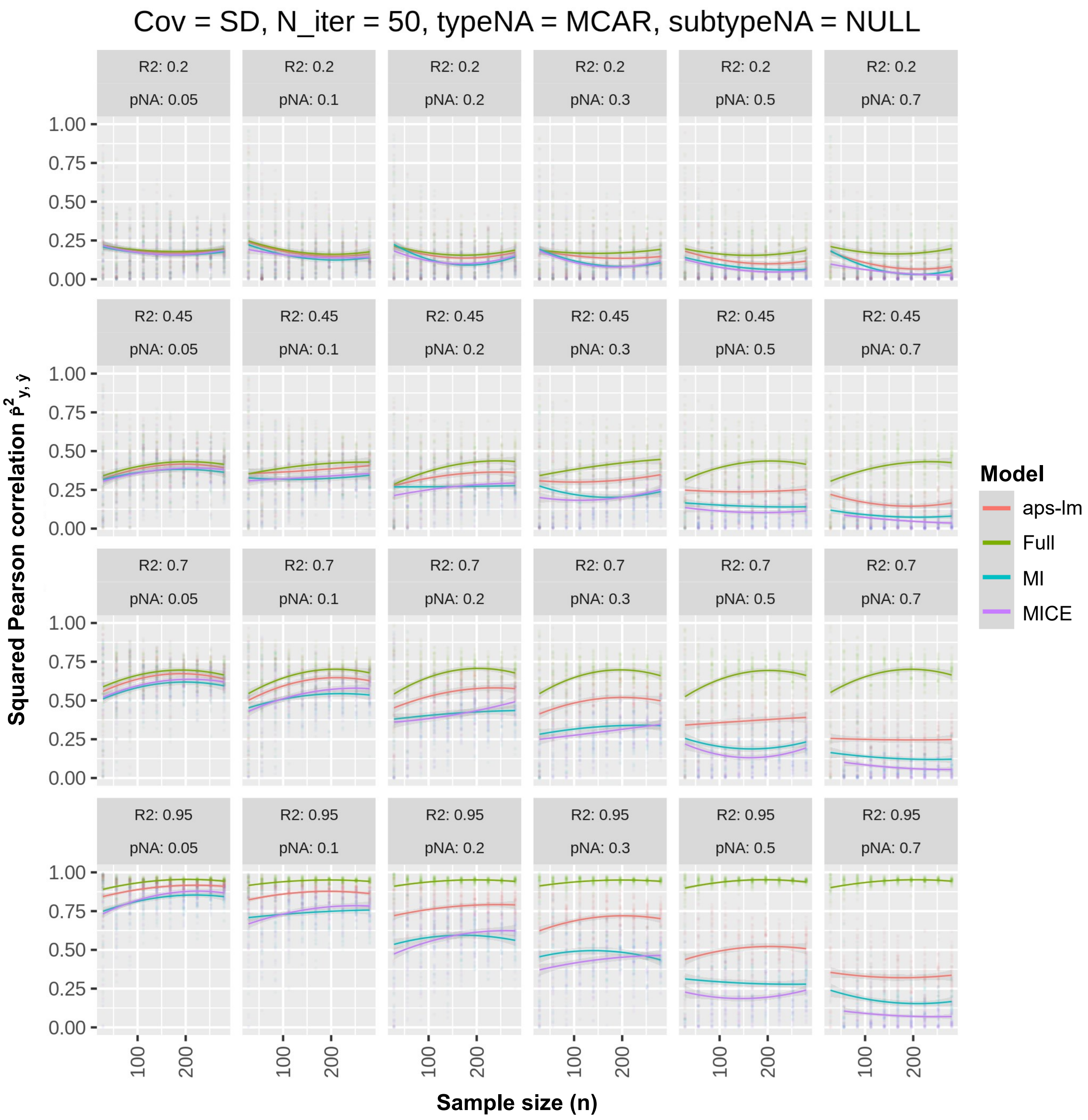


**Figure S3**. Round 1-Simulation 3: MCAR under strong dependence. For further details see legend to Fig. S2.

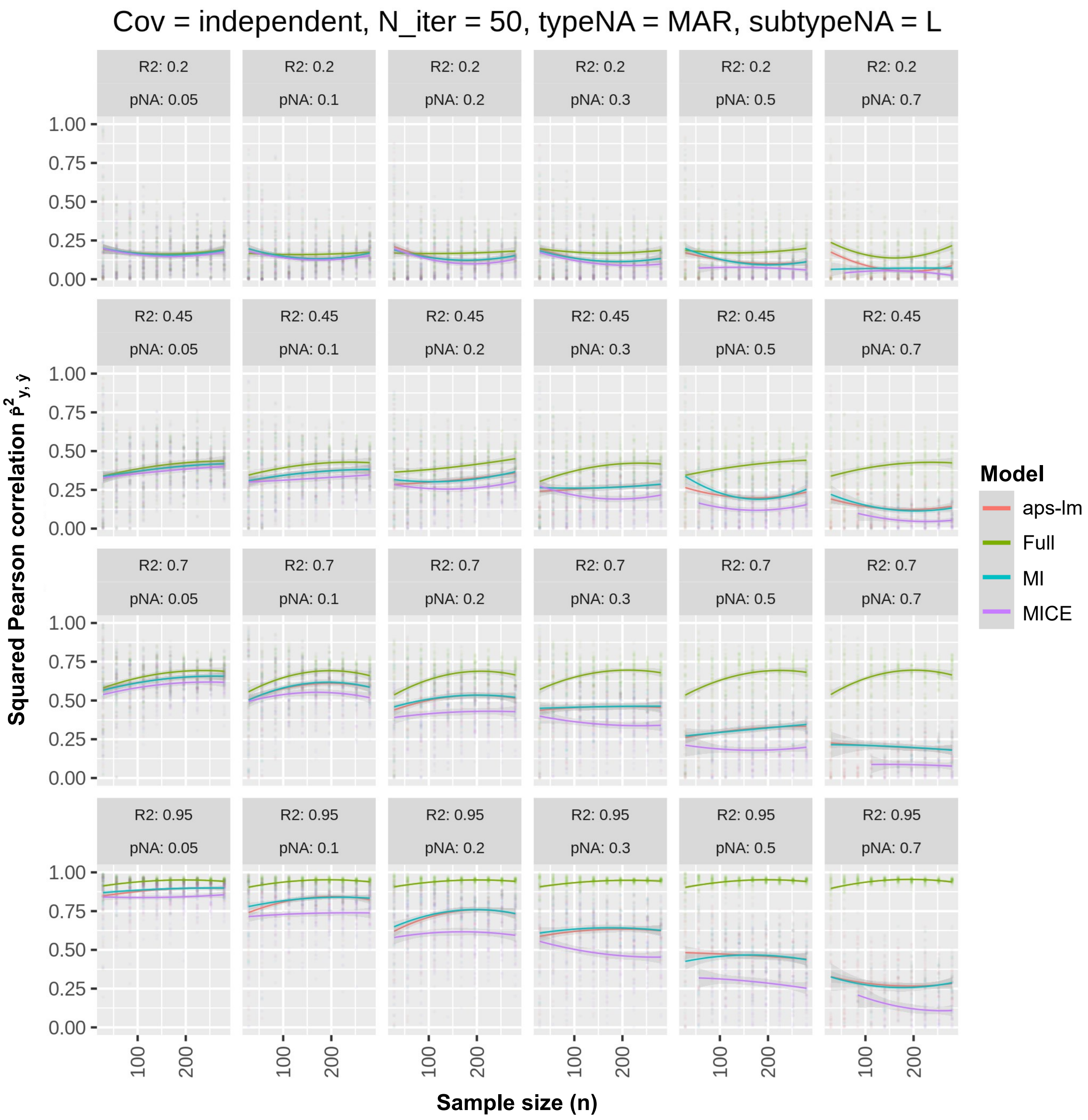


**Figure S4**. Round 1-Simulation 4: MAR-L under independence. For further details see legend to Fig. S2.

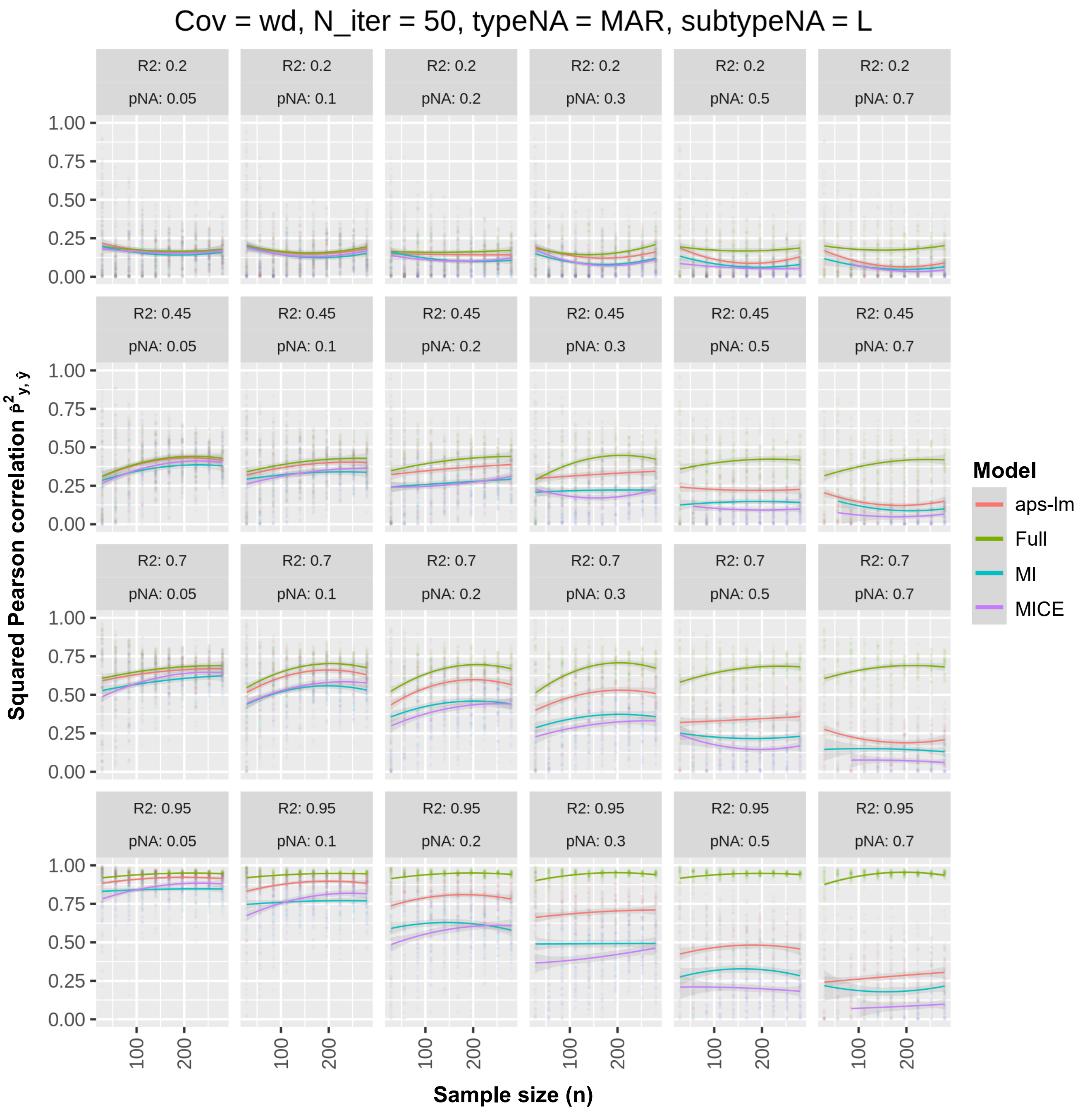


**Figure S5**. Round 1-Simulation 5: MAR-L under weak dependence. For further details see legend to Fig. S2.

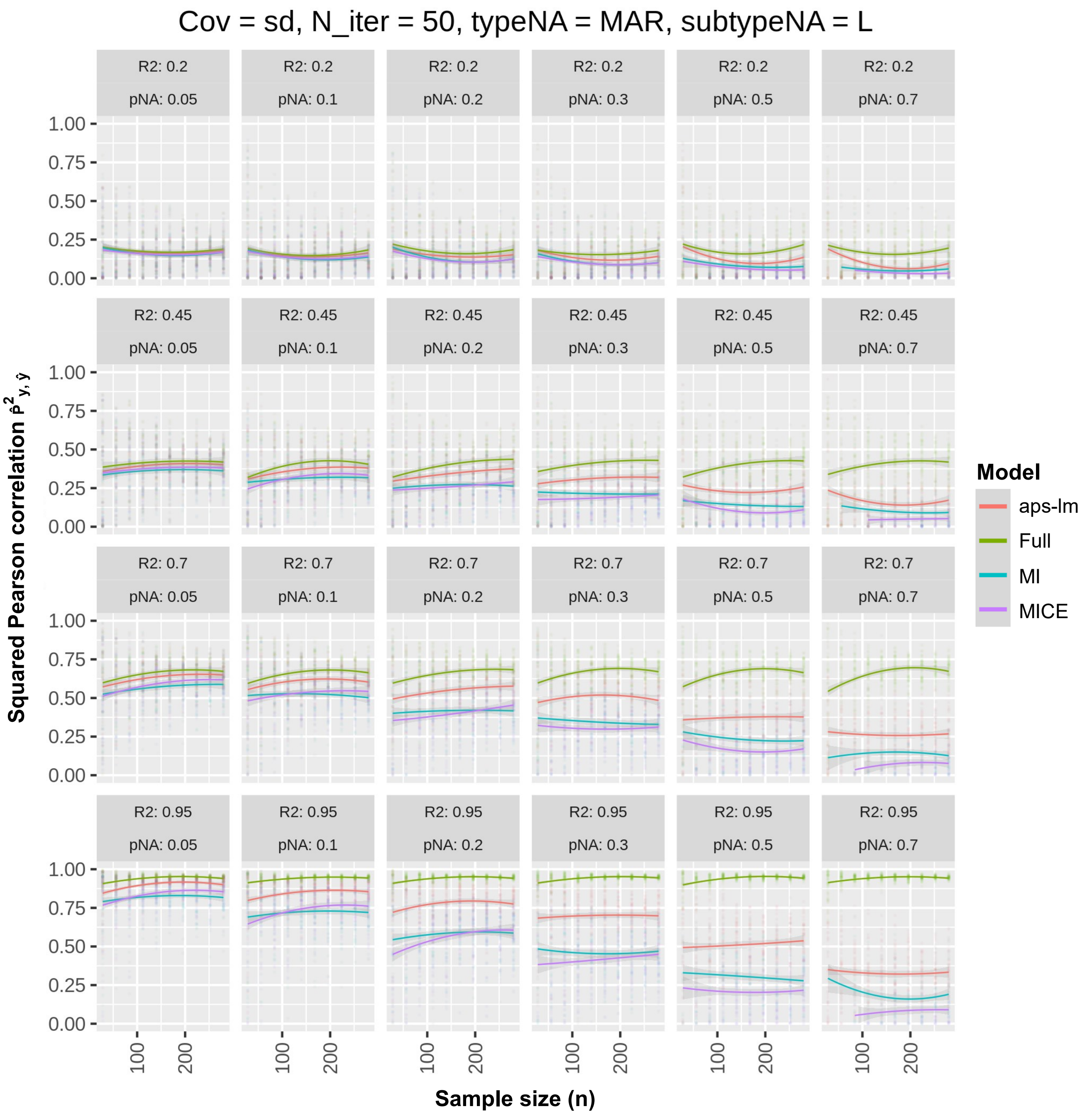


**Figure S6**. Round 1-Simulation 6: MAR-L under strong dependence. For further details see legend to Fig. S2.

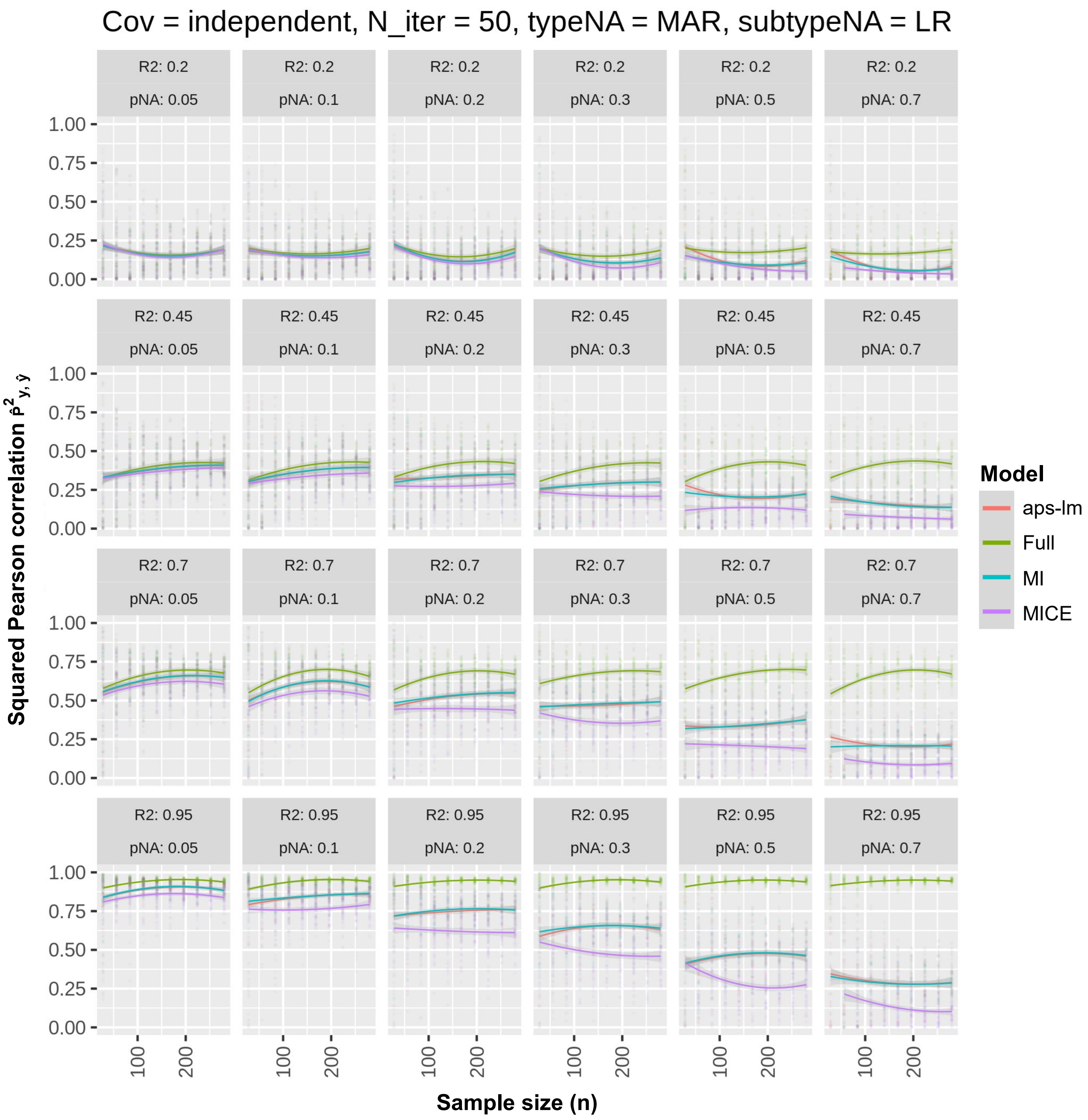


**Figure S7**. Round 1-Simulation 7: MAR-LR under independence. For further details see legend to Fig. S2.

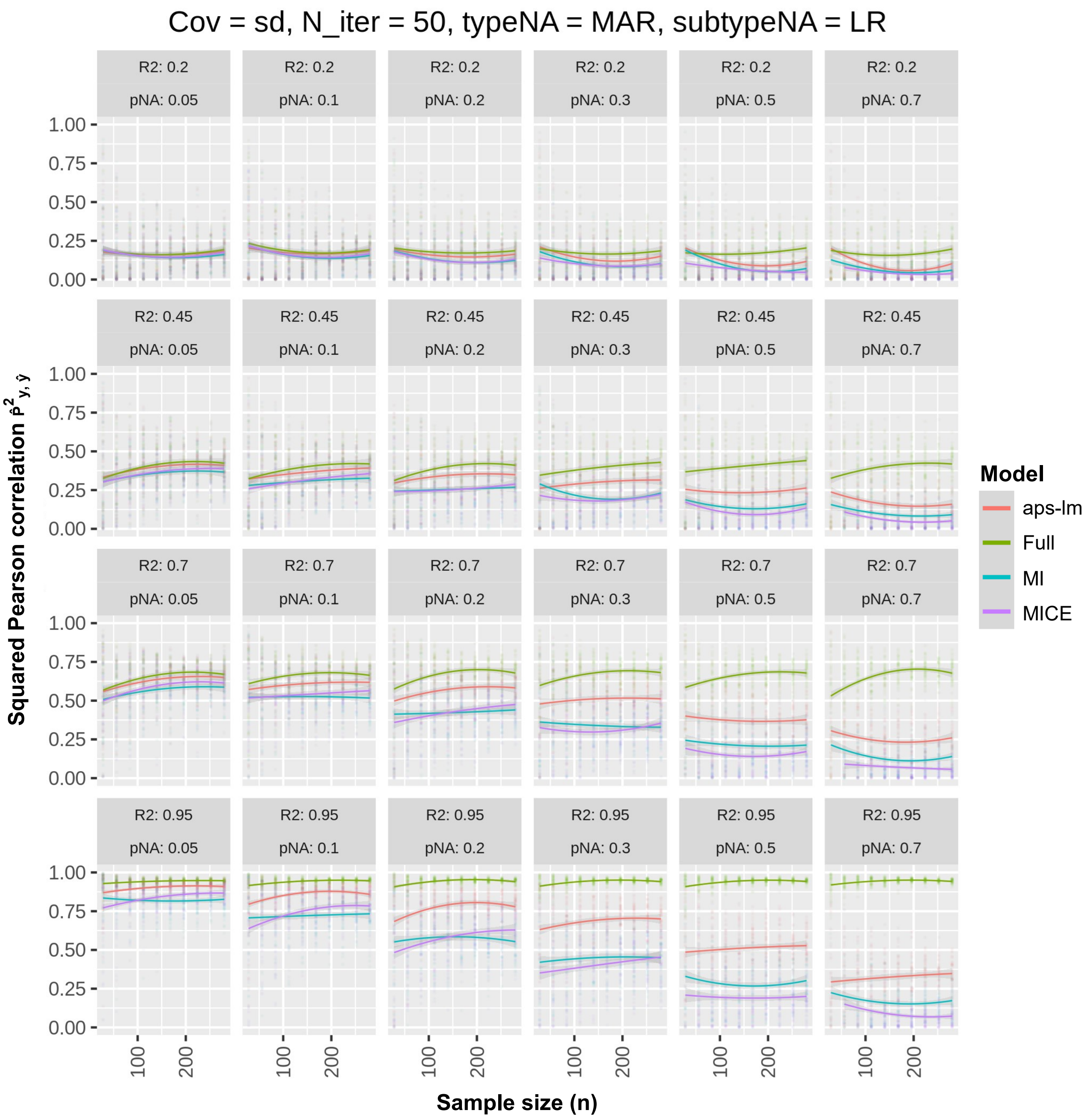


**Figure S8**. Round 1-Simulation 9: MAR-LR under strong dependence. For further details see legend to Fig. S2.

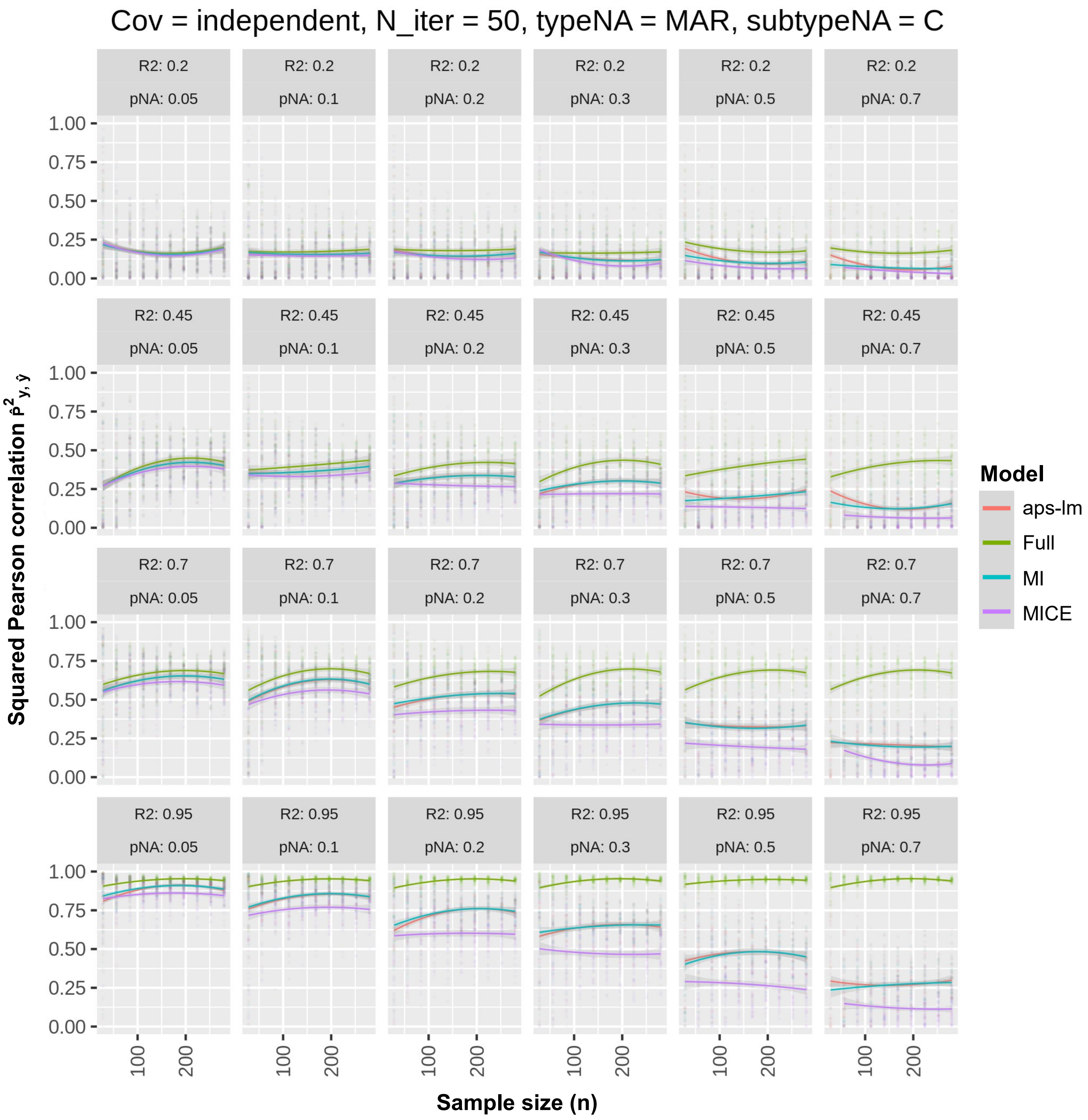


**Figure S9**. Round 1-Simulation 10: MAR-C under independence. For further details see legend to Fig. S2.

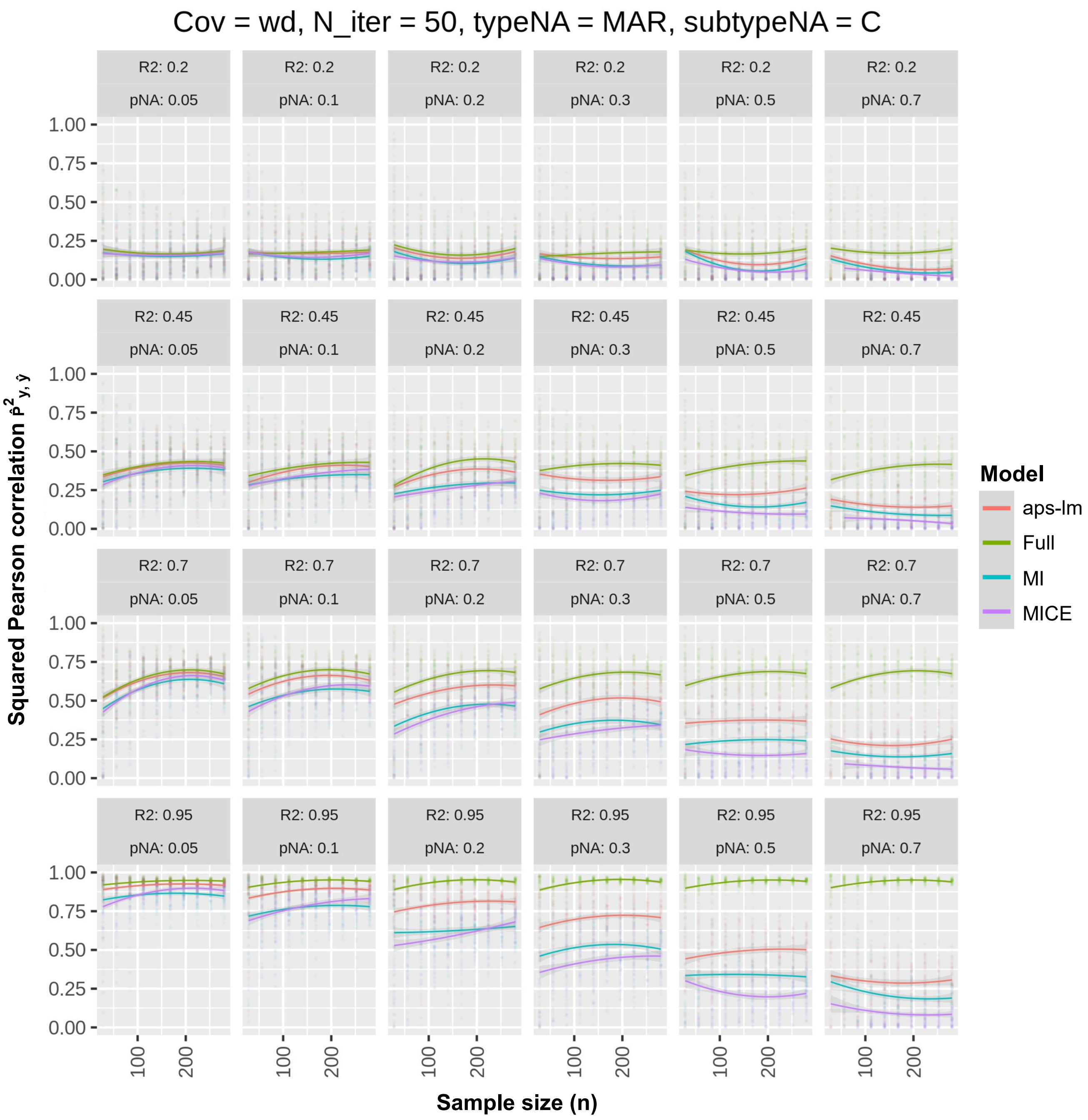


**Figure S10**. Round 1-Simulation 11: MAR-C under weak dependence. For further details see legend to Fig. S2.

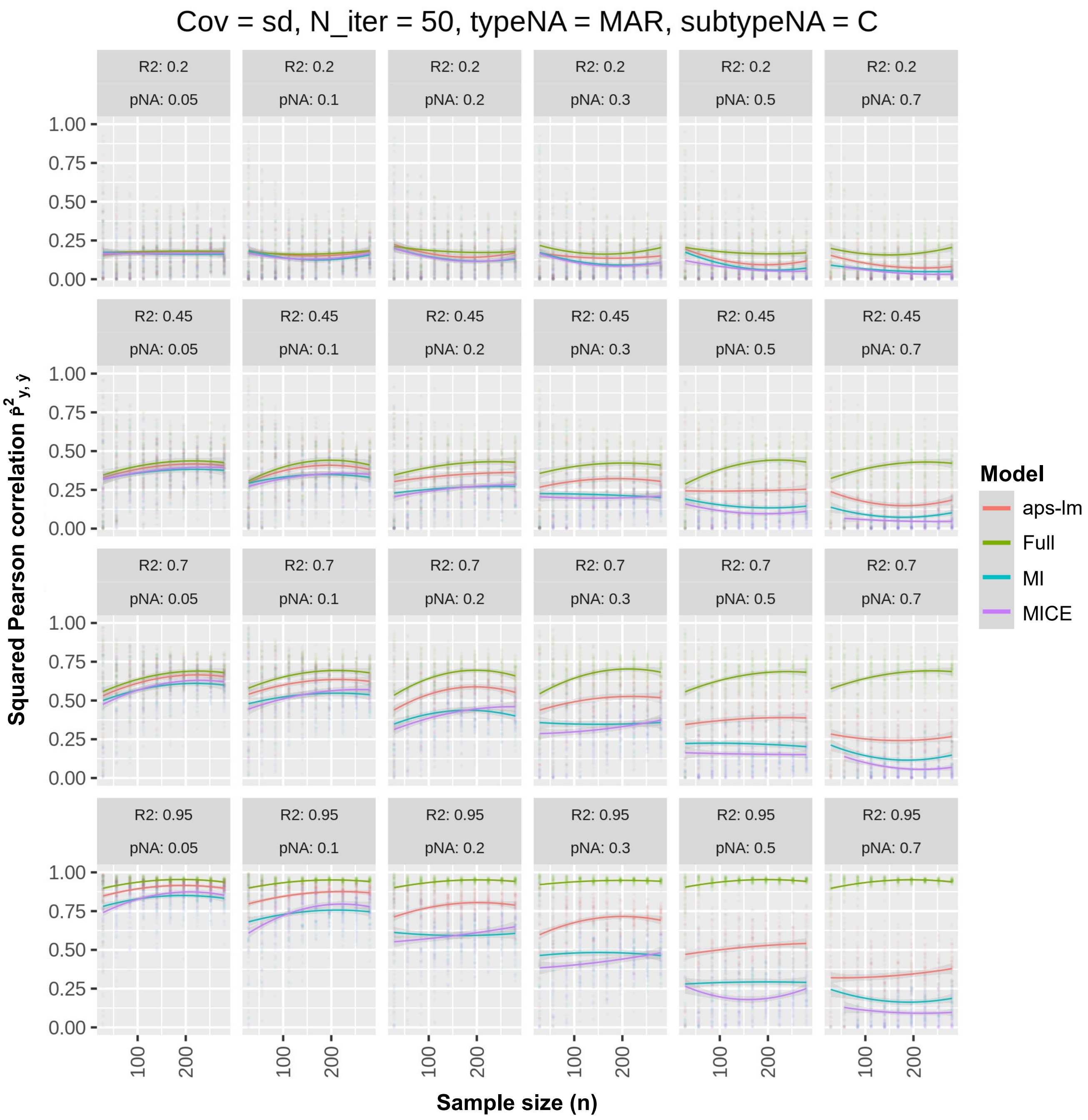


**Figure S11**. Round 1-Simulation 12: MAR-C under strong dependence. For further details see legend to Fig. S2.

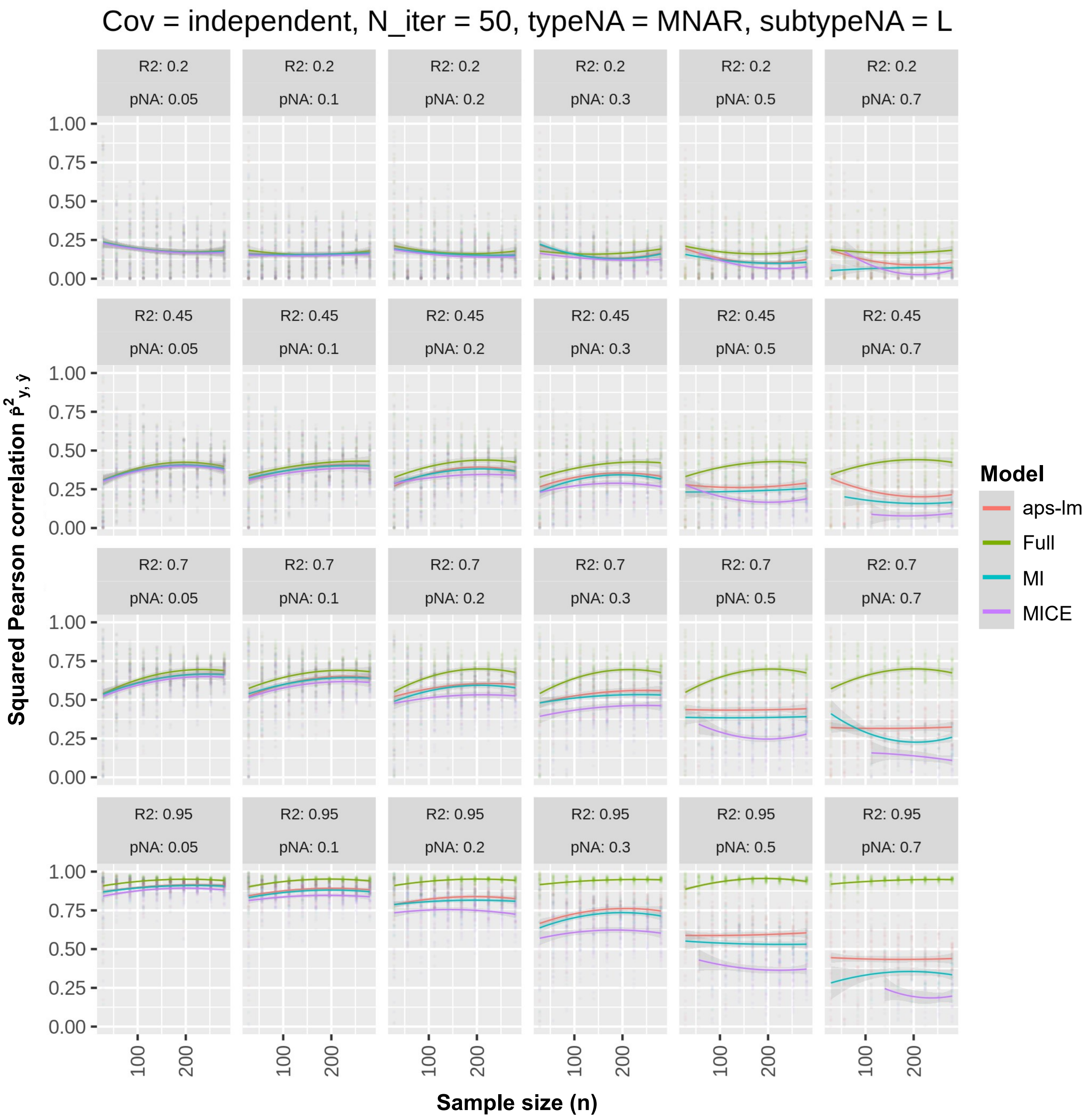


**Figure S12**. Round 1-Simulation 13: MNAR-L under independence. For further details see legend to Fig. S2.

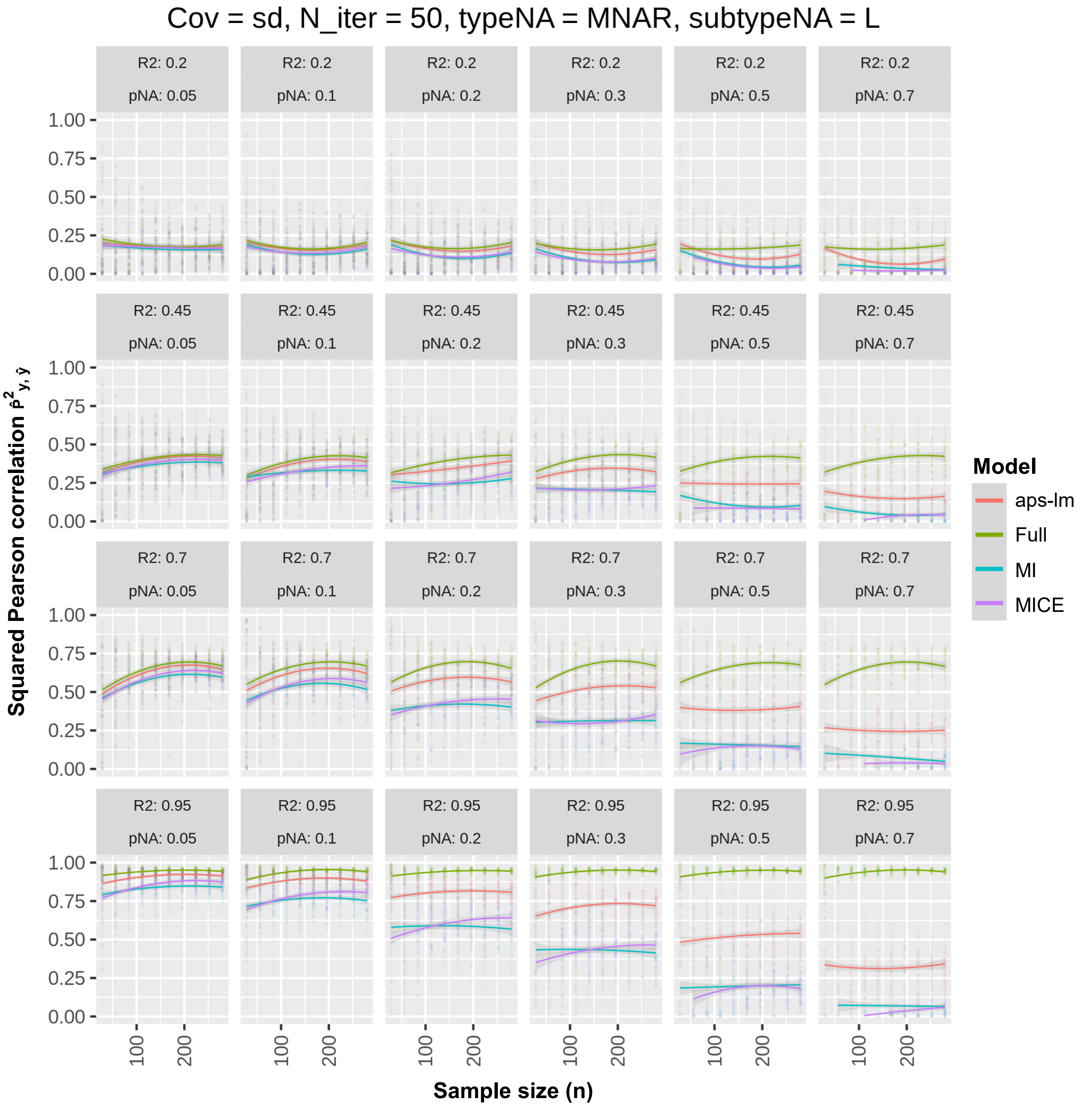


**Figure S13**. Round 1-Simulation 15: MNAR-L under strong dependence. For further details see legend to Fig. S2.

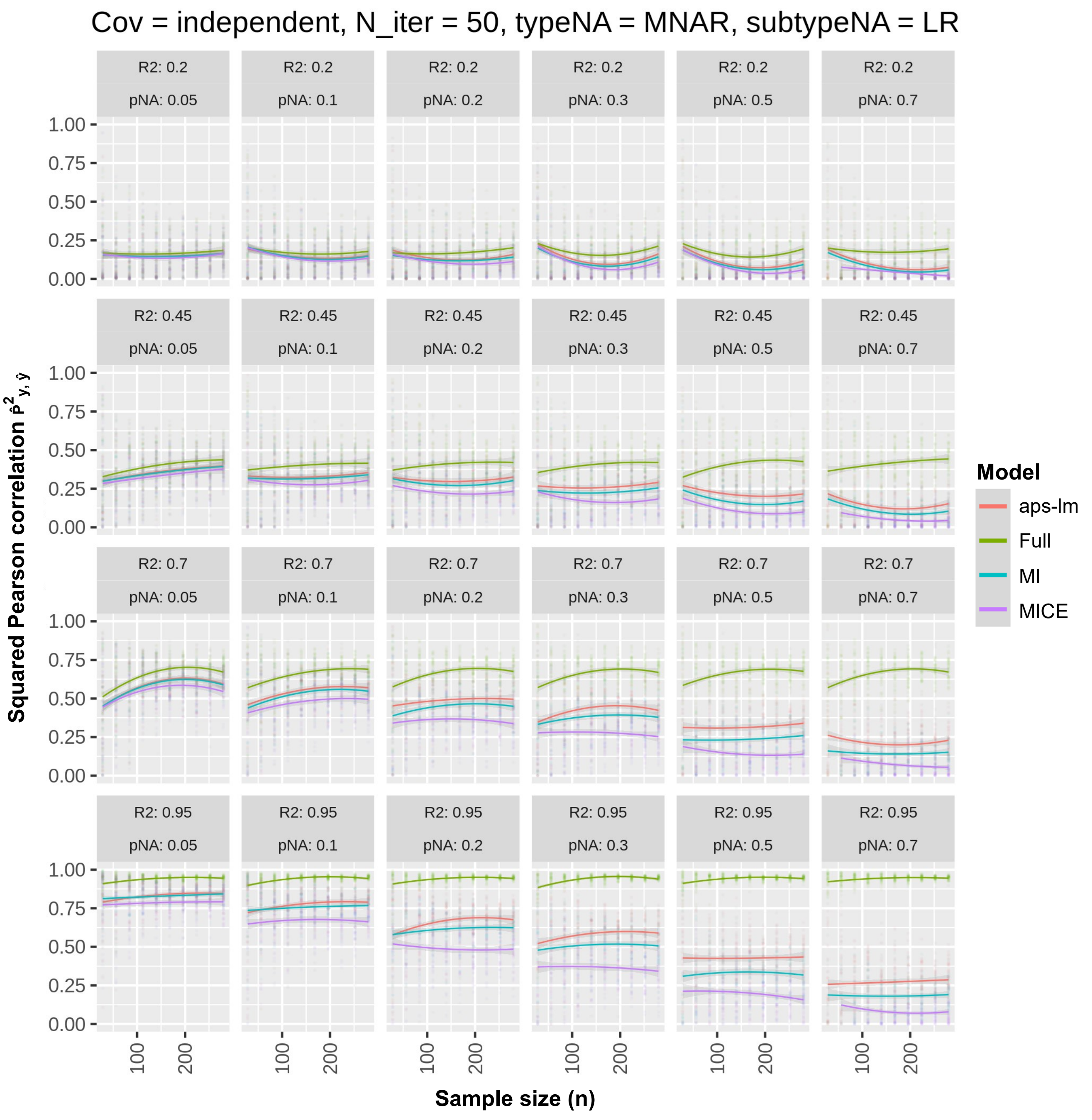


**Figure S14**. Round 1-Simulation 16: MNAR-LR under independence. For further details see legend to Fig. S2.

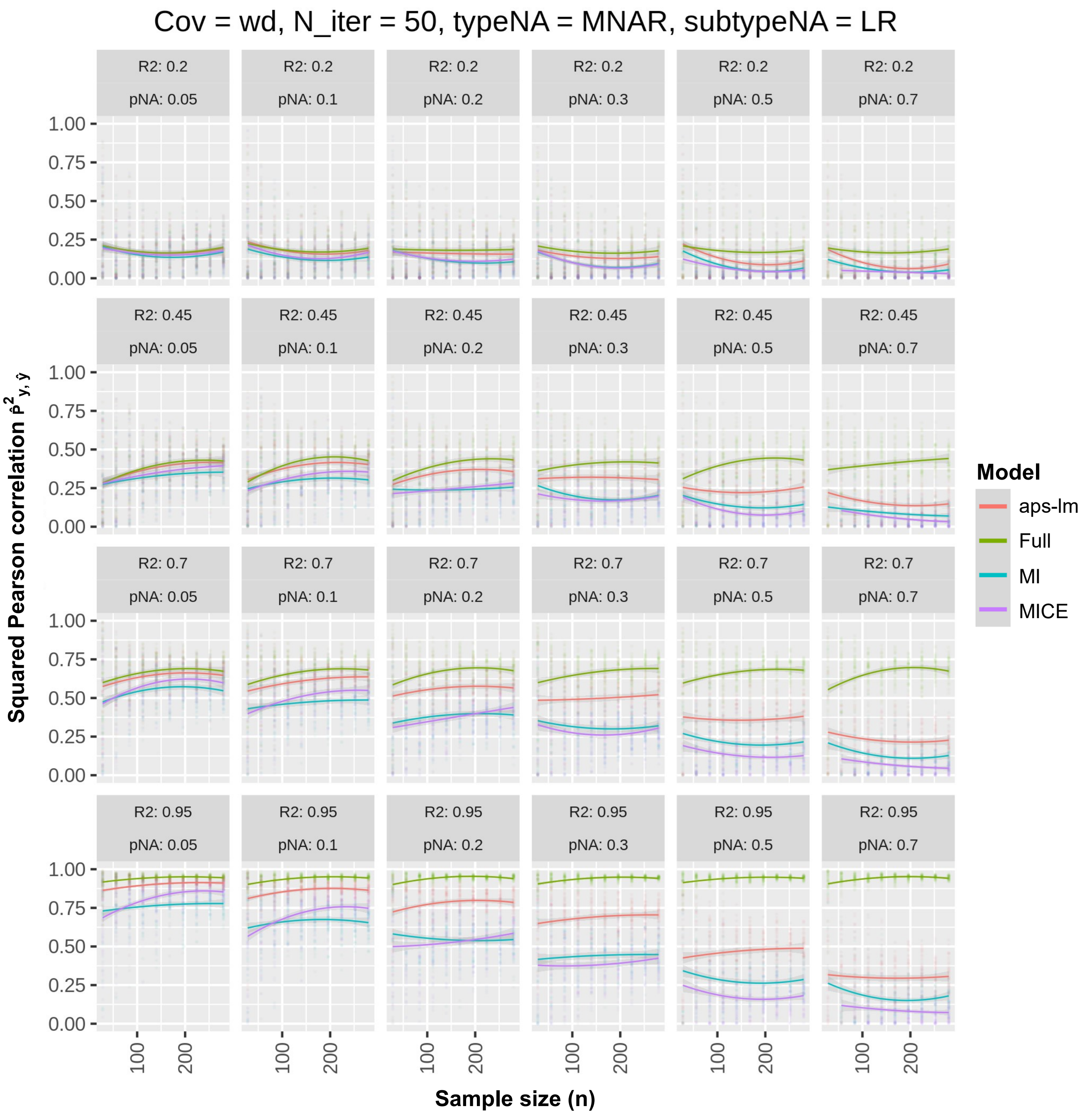


**Figure S15**. Round 1-Simulation 17: MNAR-LR under weak dependence. For further details see legend to Fig. S2.

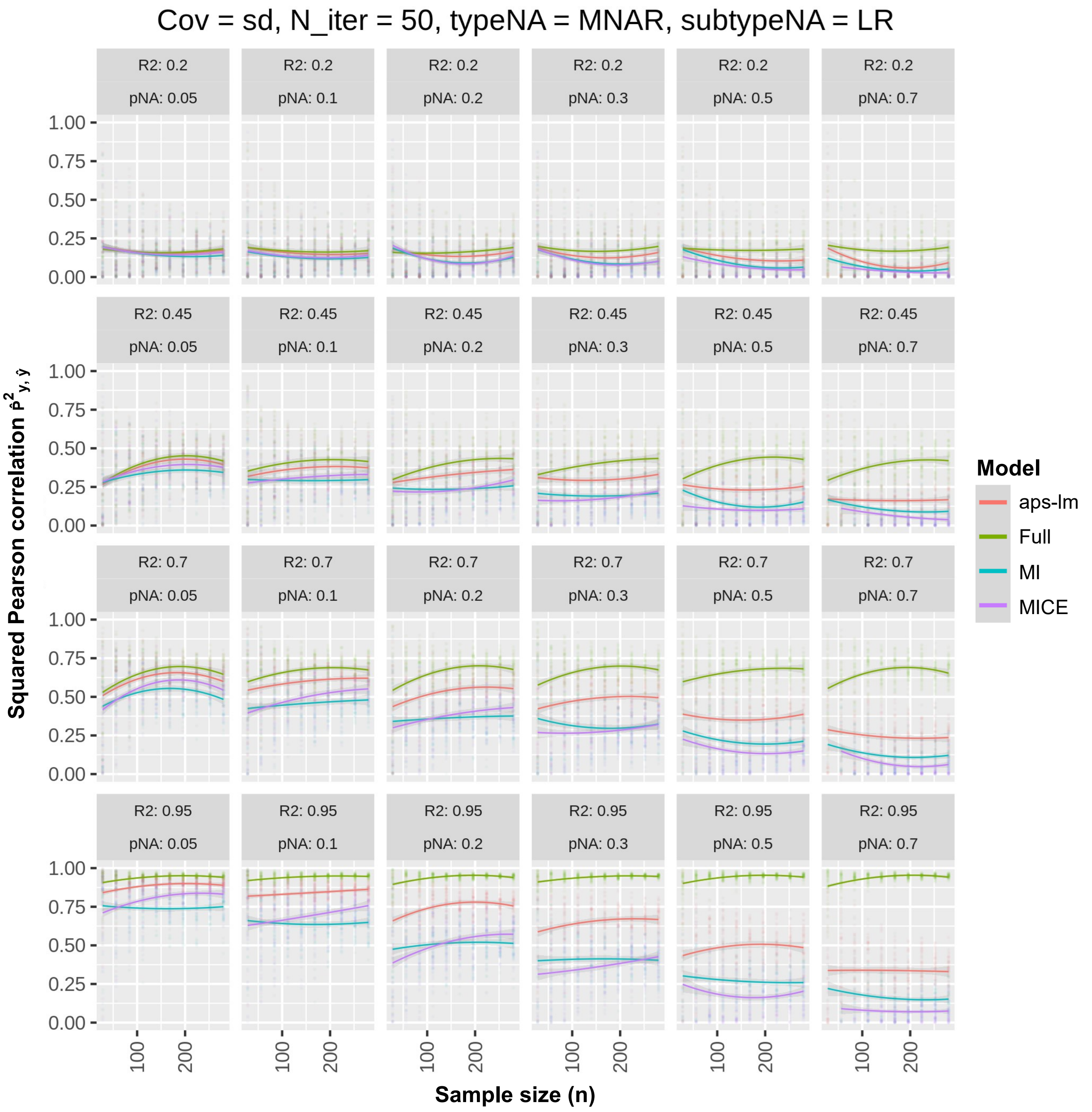


**Figure S16**. Round 1-Simulation 18: MNAR-LR under strong dependence. For further details see legend to Fig. S2.

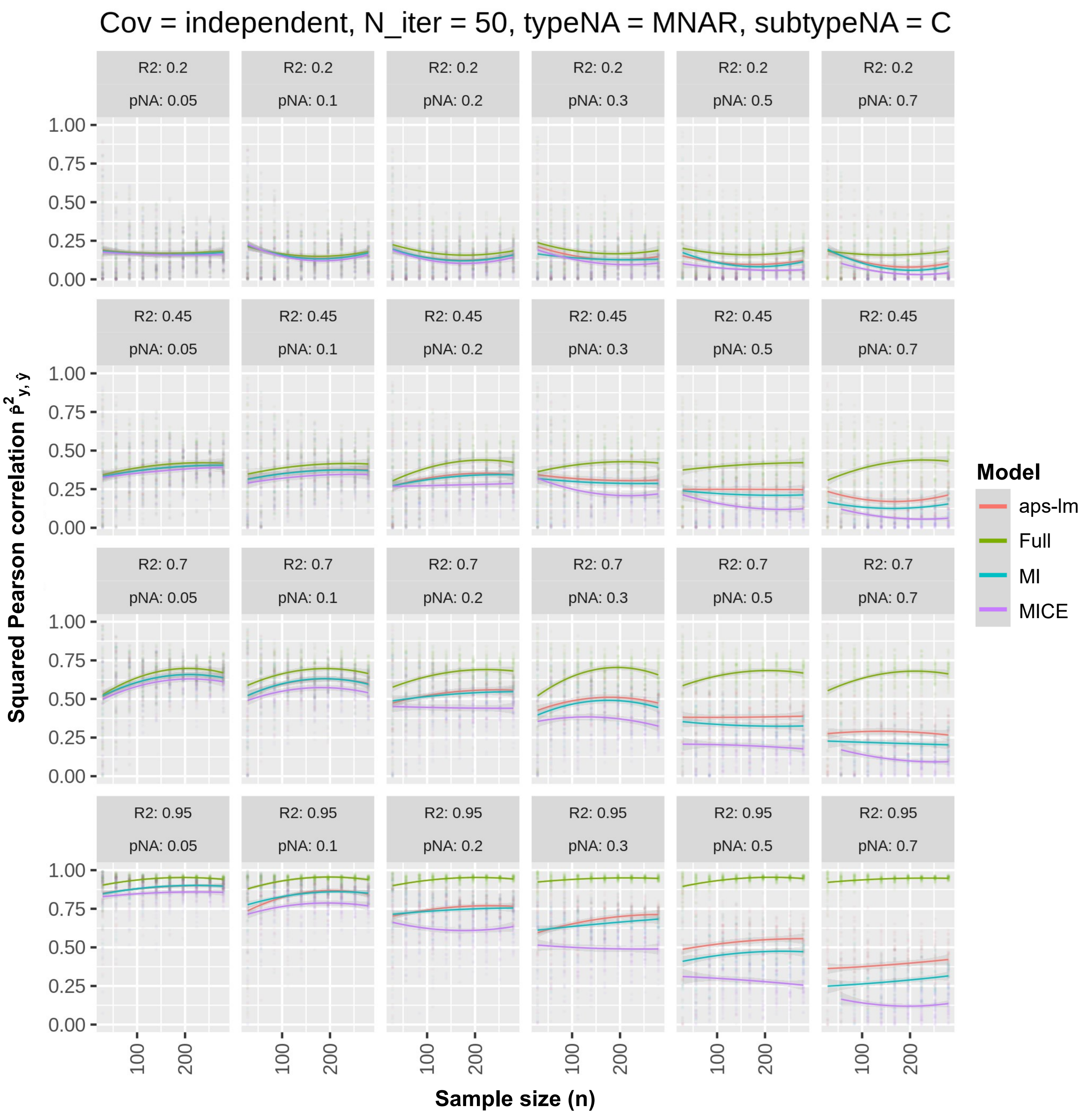


**Figure S17**. Round 1-Simulation 19: MNAR-C under independence. For further details see legend to Fig. S2.

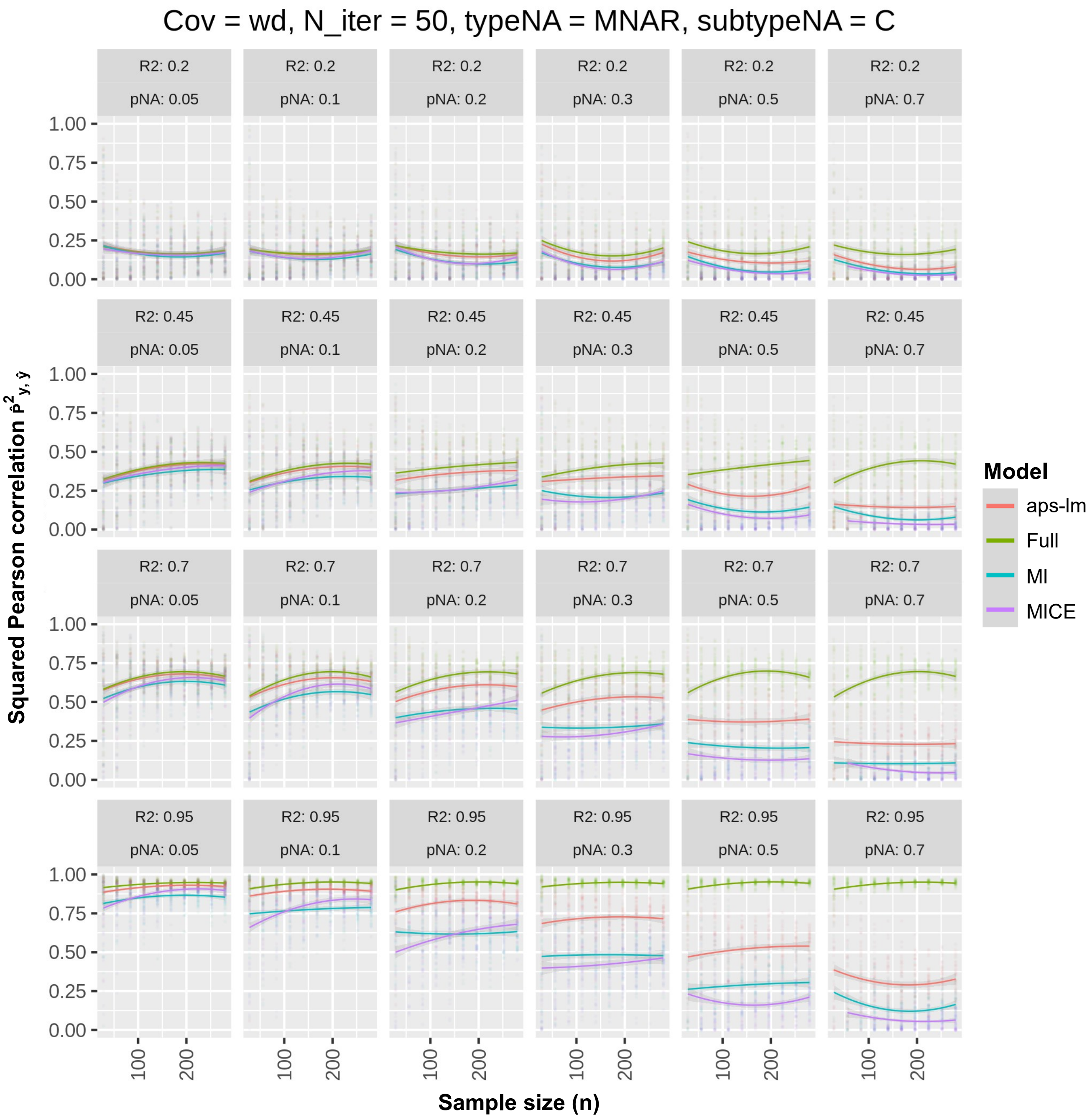


**Figure S18**. Round 1-Simulation 20: MNAR-C under weak dependence. For further details see legend to Fig. S2.

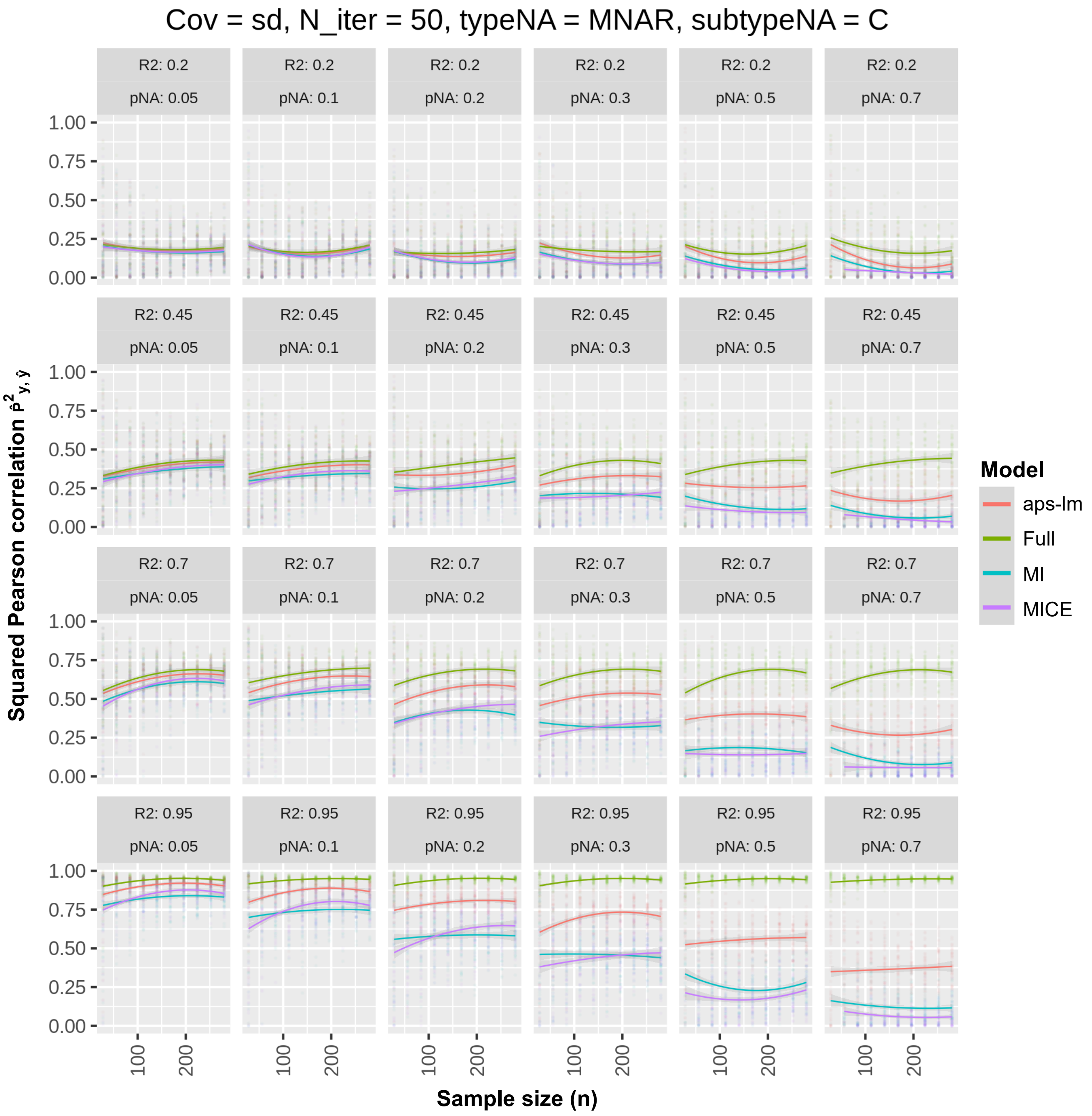


**Figure S19**. Round 1-Simulation 21: MNAR-C under strong dependence. For further details see legend to Fig. S2.

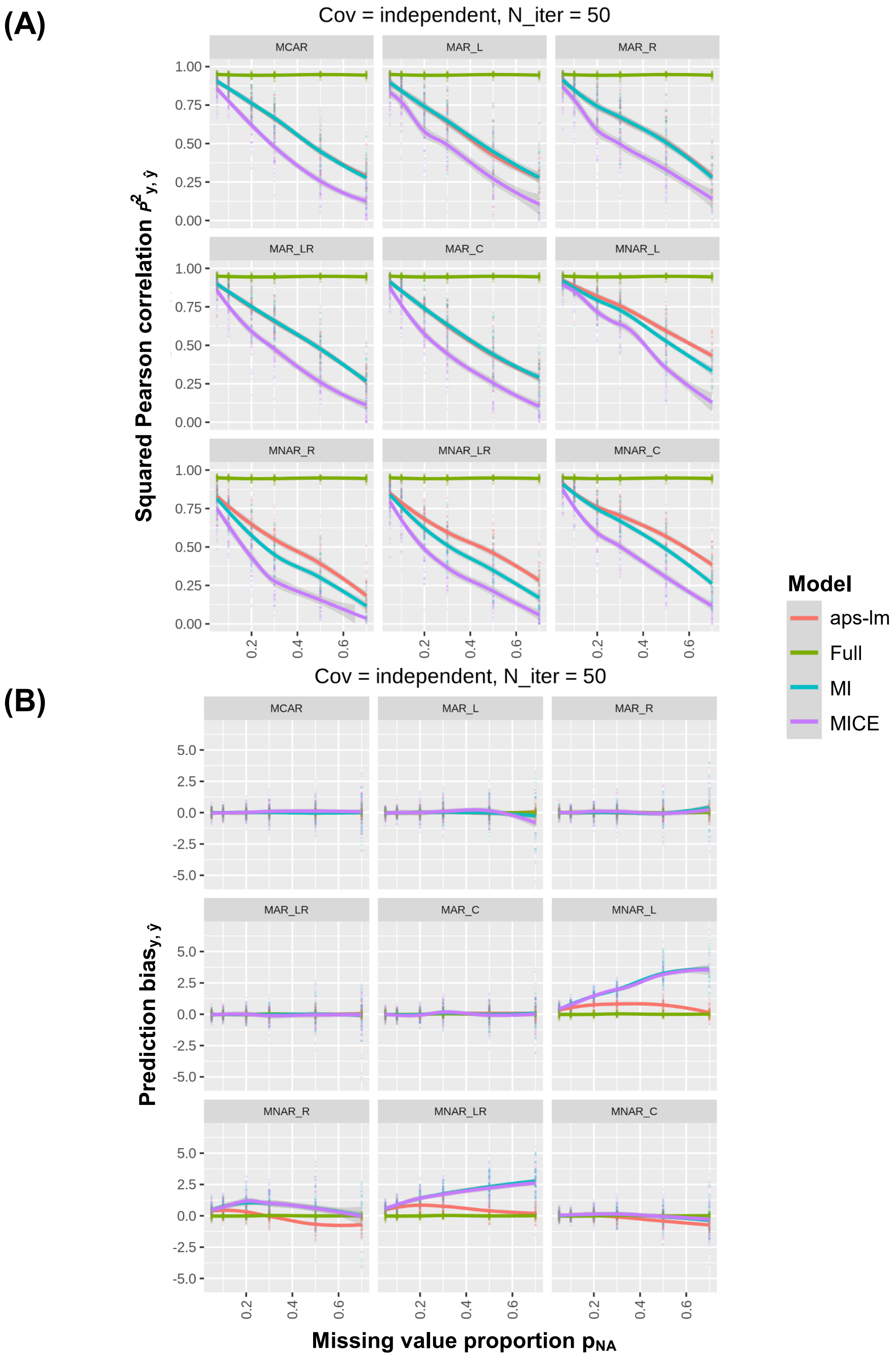


**Figure S20**. Round 2-Simulation 1: all missing value types under independence employing θ$_1$. Missing values were injected on the application dataset following all implemented missing value schemes (MCAR, MAR-L,R, LR, C and MNAR-L, R, LR, C). (A) Squared Pearson correlation between real and predicted dependent variable or (B) prediction bias is plotted at varying missing value type, $R^2$ and $p_{NA}$. For further details see legend to Fig. S2.

(A)

(B)

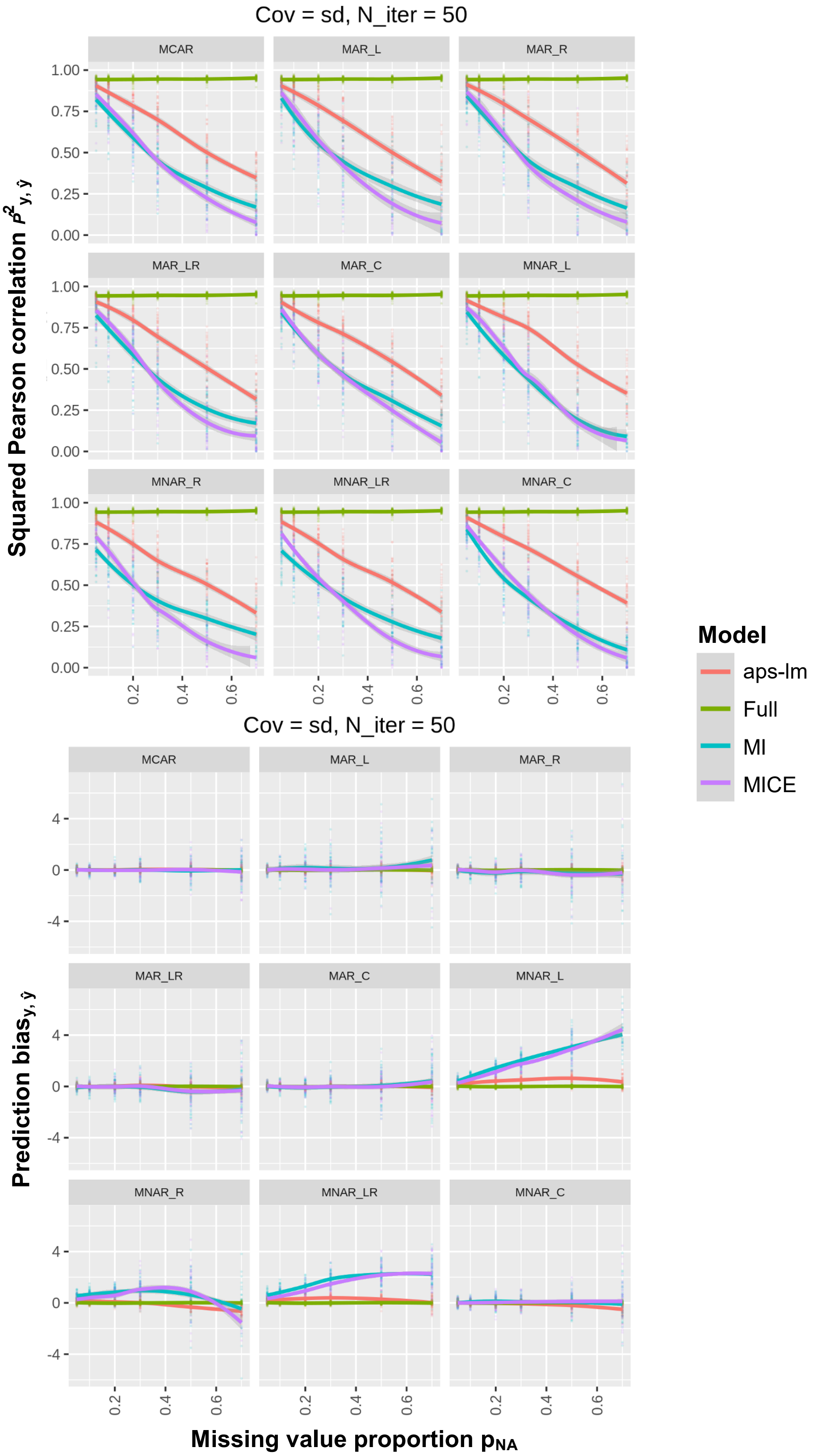


**Figure S21**. Round 2-Simulation 3: all missing value types under strong dependence employing $\theta_1$. For further details see legend to Fig. S20.

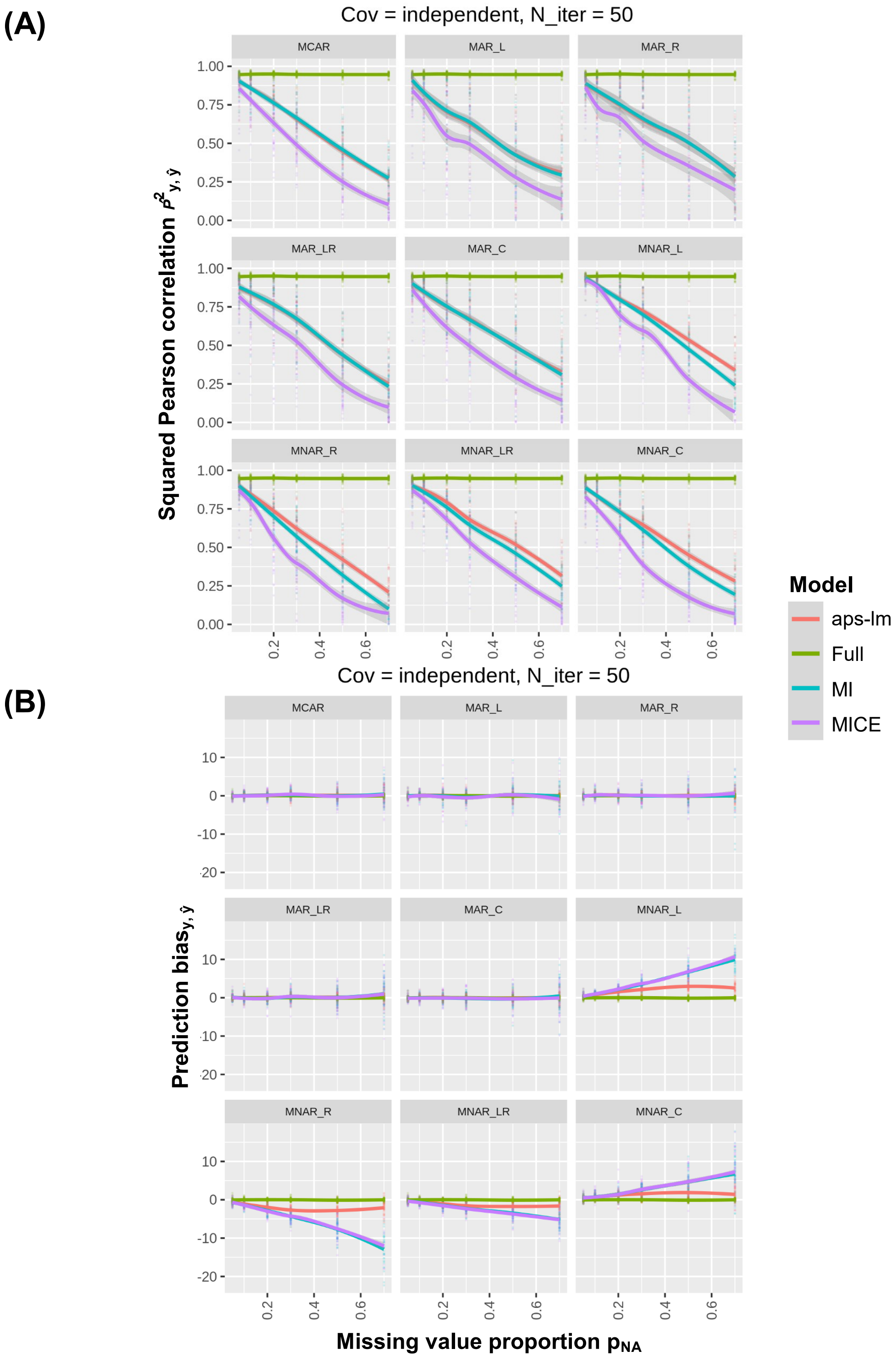


**Figure S22**. Round 2-Simulation 5: all missing value types under independence employing $\theta_2$. For further details see legend to Fig. S20.

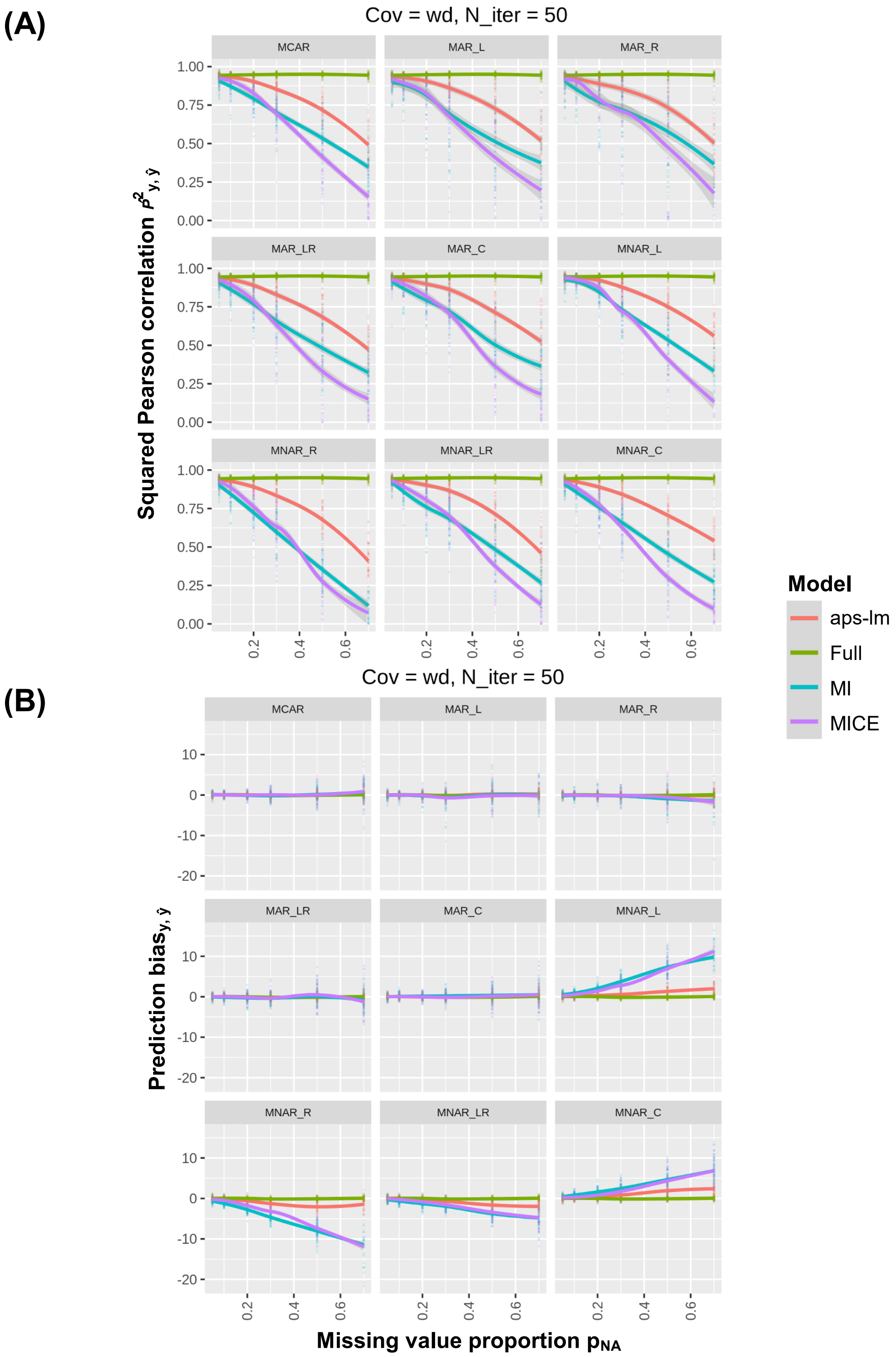


**Figure S23**. Round 2-Simulation 6: all missing value types under weak dependence employing $\theta_2$. For further details see legend to Fig. S20.

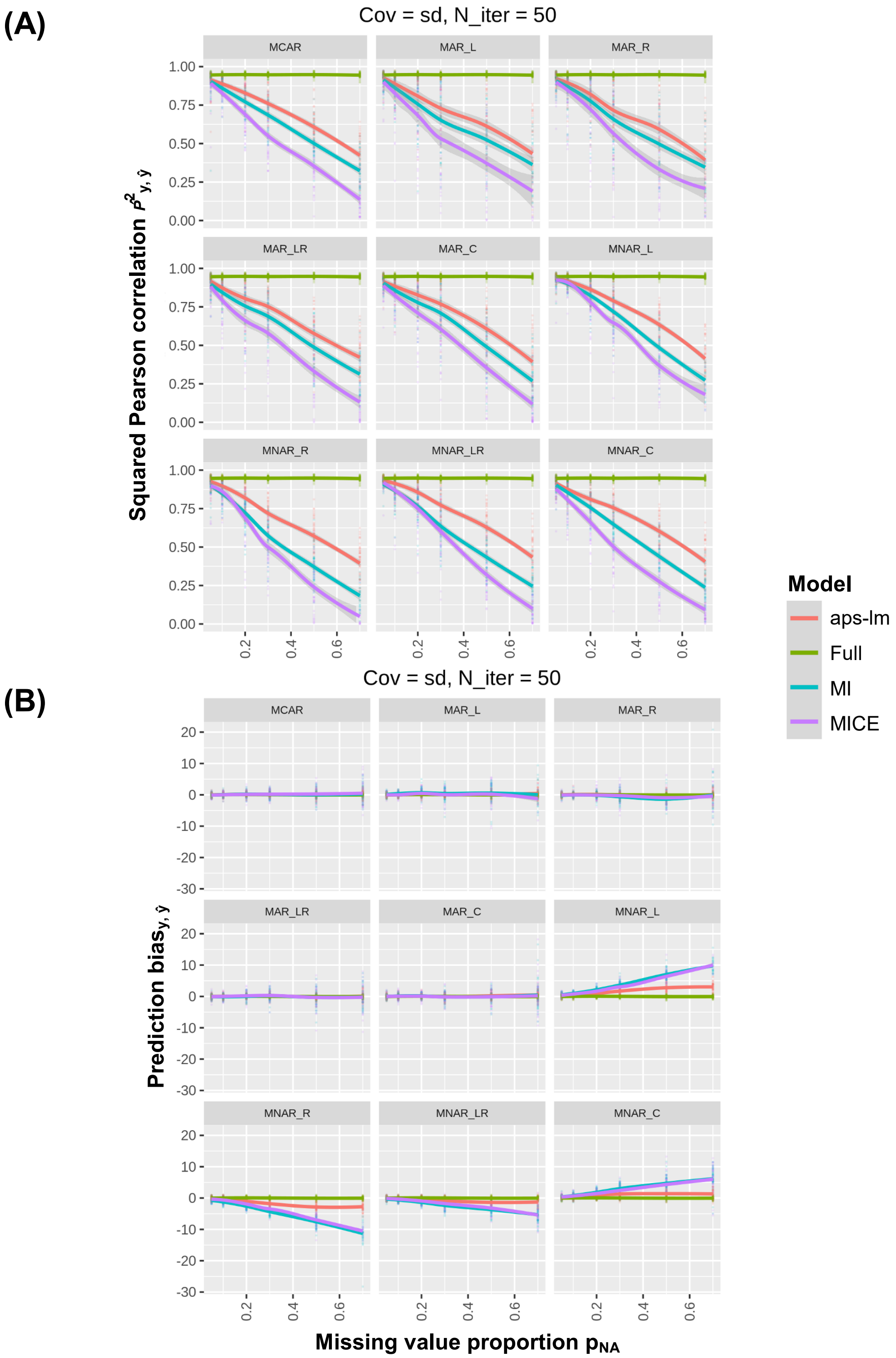


**Figure S24**. Round 2-Simulation 7: all missing value types under strong dependence employing $\theta_2$. For further details see legend to Fig. S20.

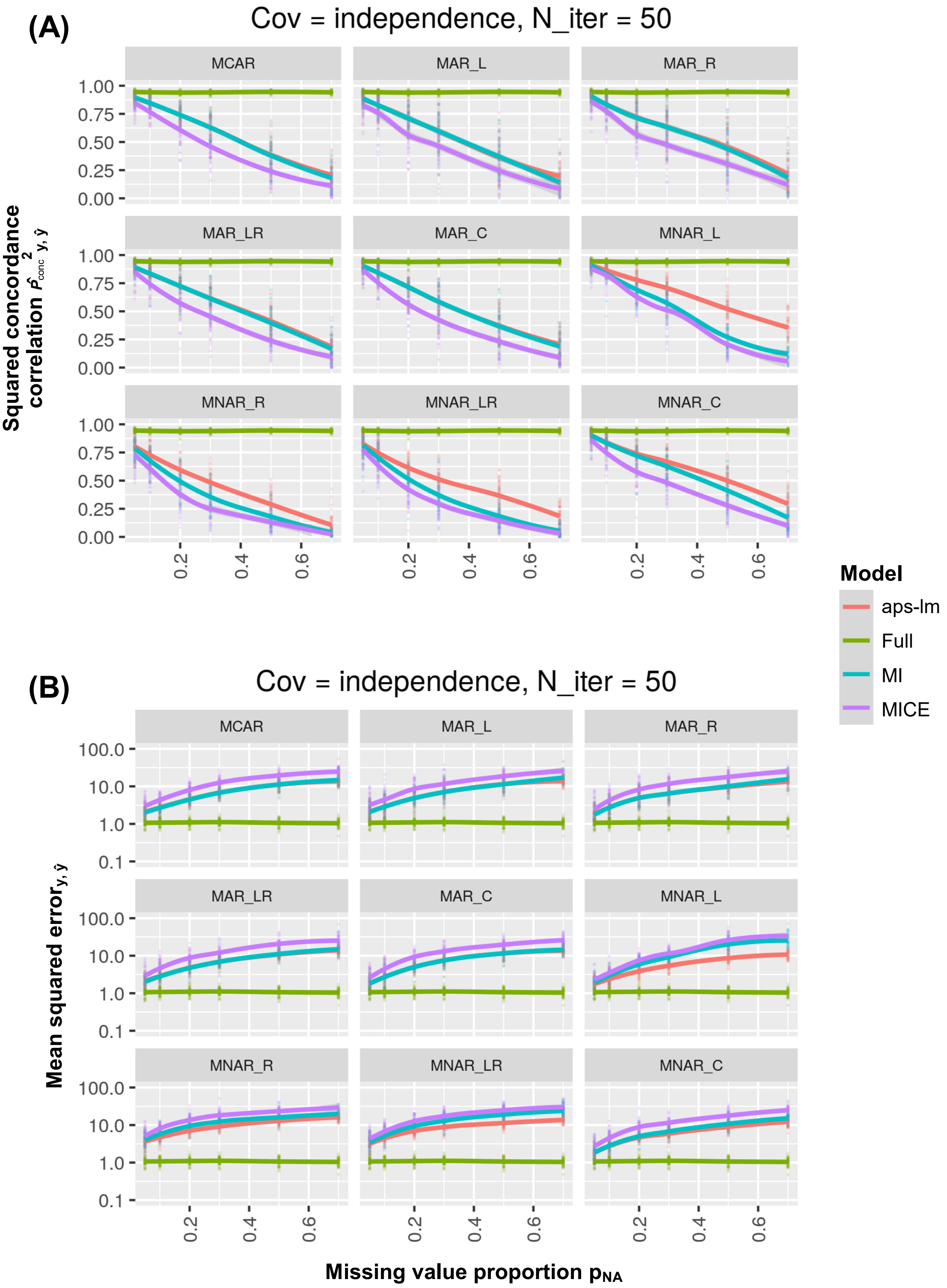


**Figure S25**. Round 2-Simulation 1: all missing value types under independence employing $\theta_1$. Missing values were injected on the application dataset following all implemented missing value schemes (MCAR, MAR-L,R, LR, C and MNAR-L, R, LR, C). (A) Squared concordance correlation between real and predicted dependent variable or (B) prediction mean squared error is plotted at varying missing value type, $R^2$ and $p_{NA}$. Note that the mean squared error axis is in log-scale.
For further details see legend to Fig. S2.

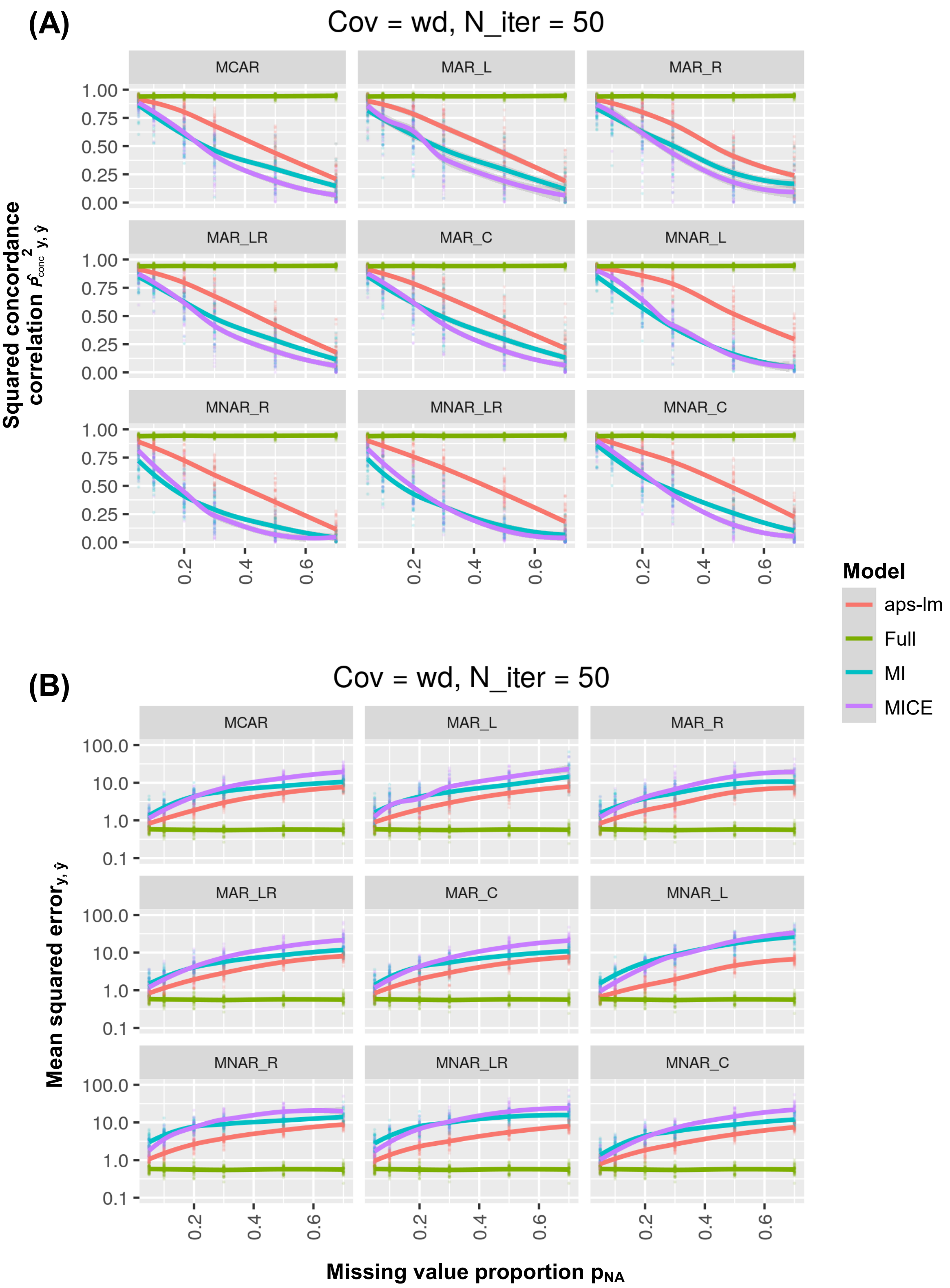


**Figure S26**. Round 2-Simulation 2: all missing value types under weak dependence employing $\theta_1$. For further details see legend to Fig. S25.

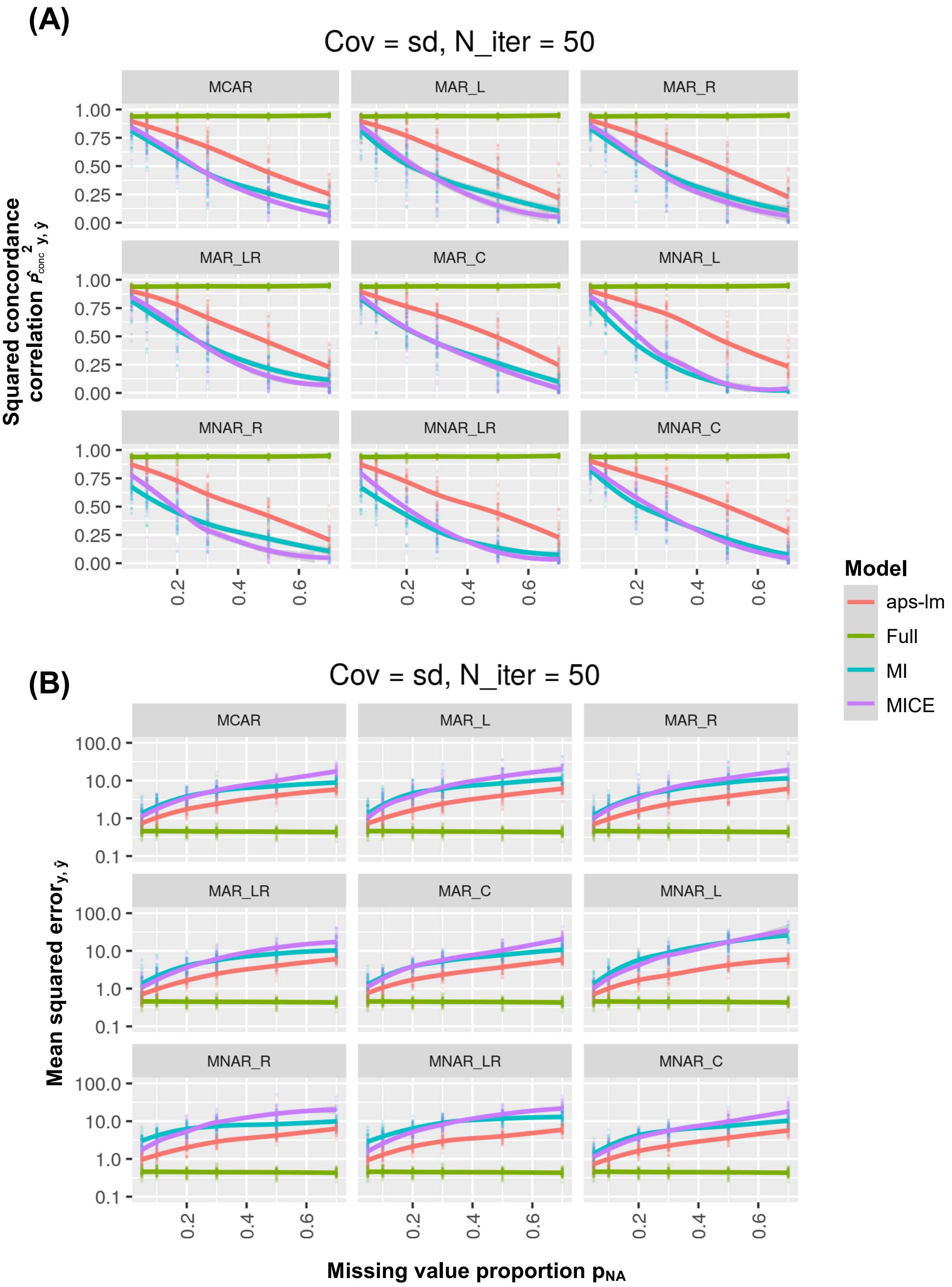


**Figure S27**. Round 2-Simulation 3: all missing value types under strong dependence employing $\theta_1$. For further details see legend to Fig. S25.

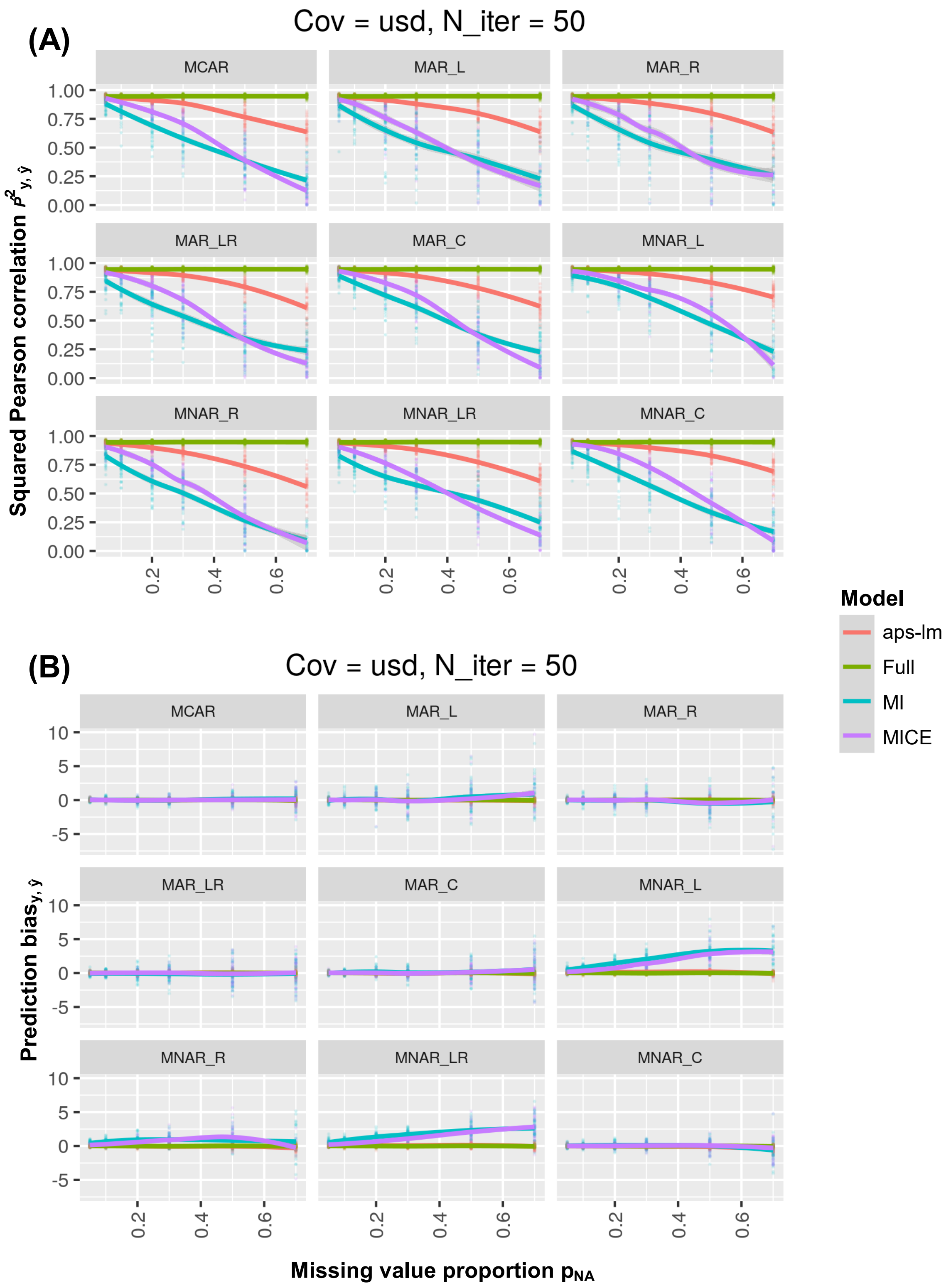


**Figure S28.** Round 2-Simulation 4: all missing value types under ultra-strong dependence employing $\theta_1$. For further details see legend to Fig. S20.

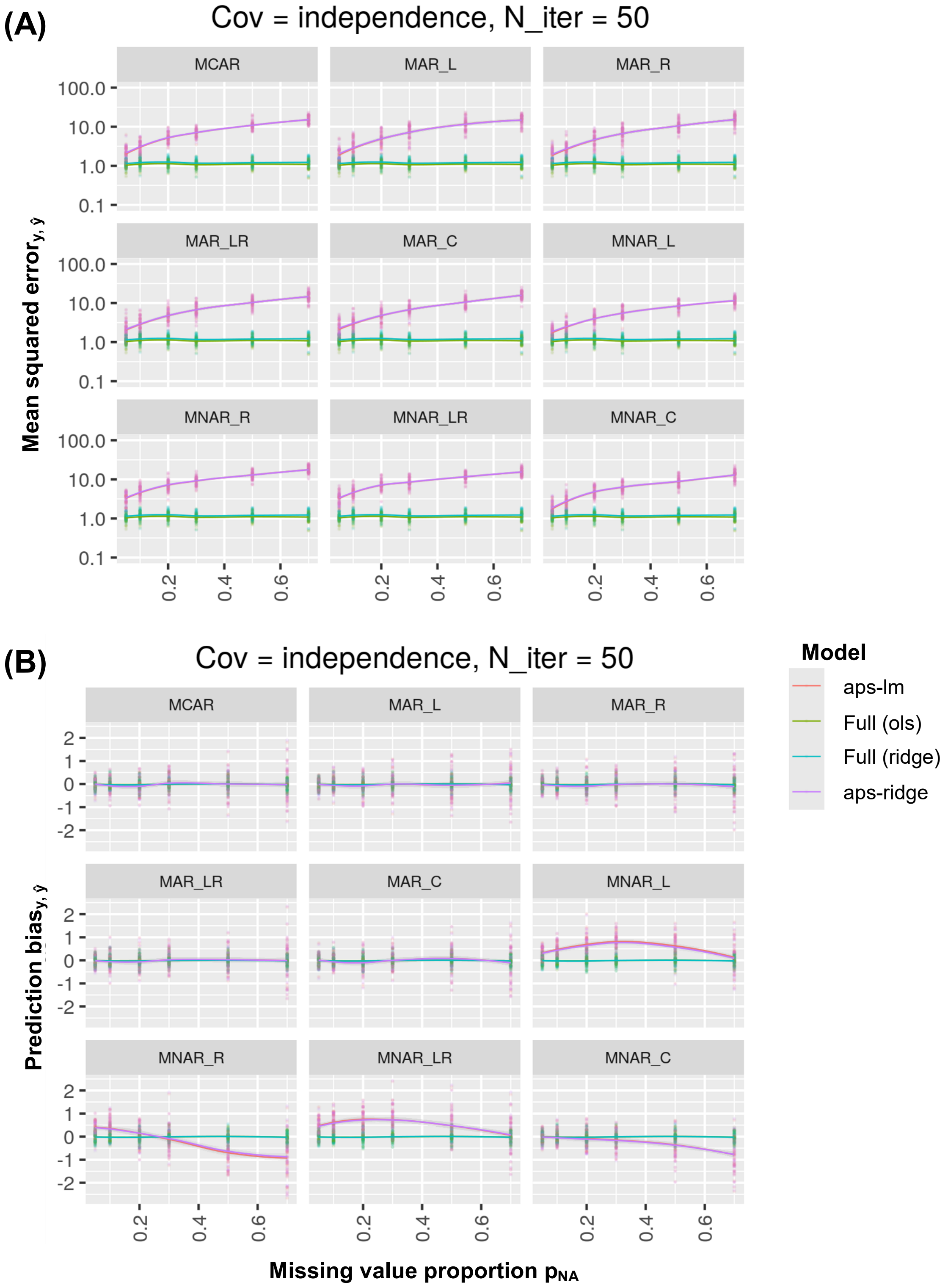


**Figure S29**. Round 3-Simulation 1: all missing value types under independence employing $\theta_1$. Missing values were injected on the application dataset following all implemented missing value schemes (MCAR, MAR-L,R, LR, C and MNAR-L, R, LR, C). (A) Prediction mean squared error between real and predicted dependent variable or (B) prediction bias is plotted at varying missing value type, $R^2$ and $p_{NA}$. This is shown for the estimated ordinary least squares and ridge linear regression model of stage 1 applied on the application dataset $X_{app}$ without missing data (i) full (ols), ii) full (ridge), respectively) and iii) aps-lm and iv) aps-ridge without applying any imputation. The regularization parameter for full (ridge) and aps-ridge was set equally and obtained via cross-validation on the reference dataset based on the prediction mean-squared error metric. Note that the mean squared error axis is in log-scale. Both curves aps-ridge and aps-lm, and curves full (ols) and full (ridge) are completely overlaying in panel A and B.

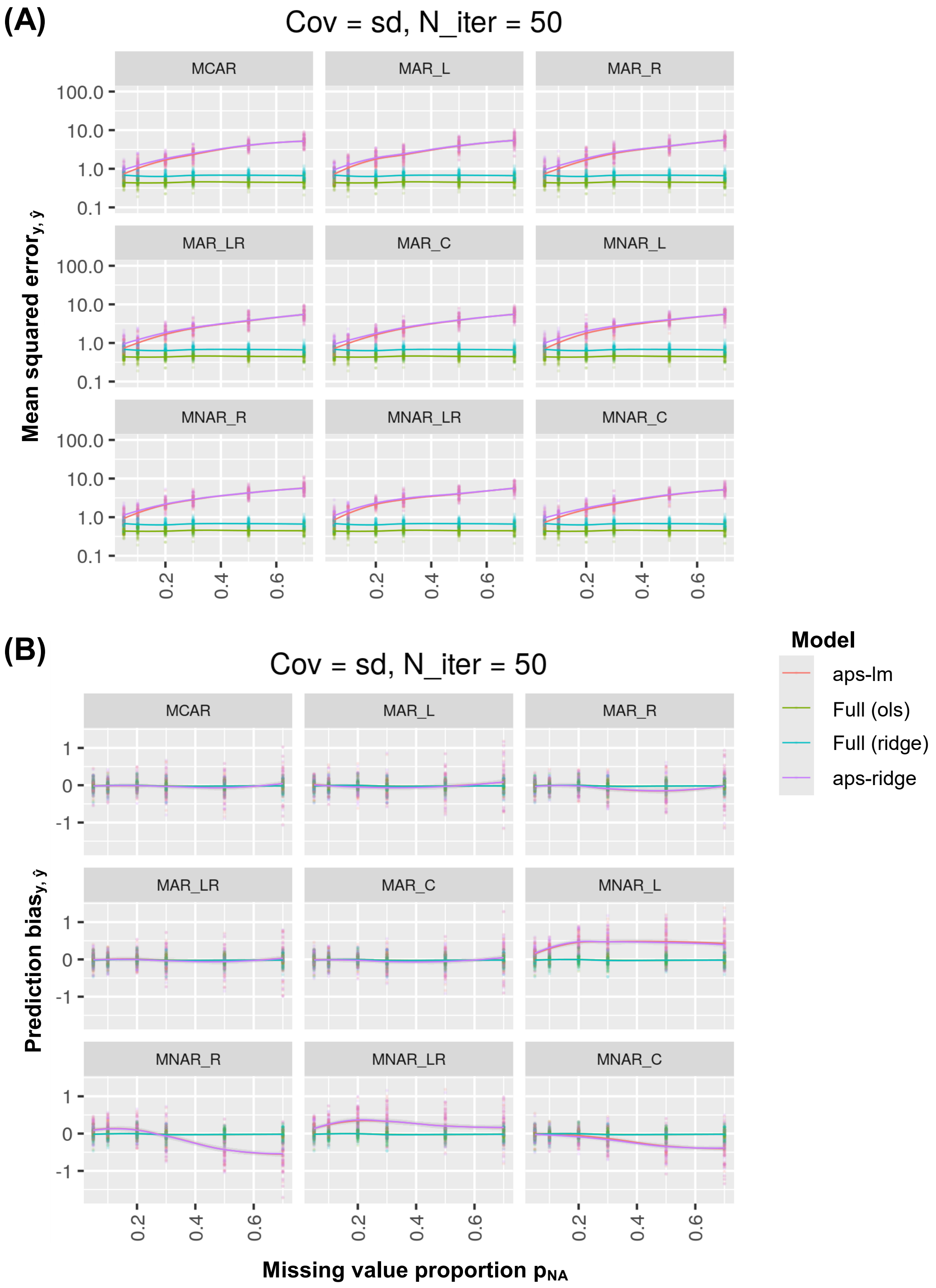


**Figure S30.** Round 3-Simulation 3: aps-lm vs aps-ridge for all missing value types under strong dependence. Curves aps-ridge and aps-lm are partially overlaying in panel A and completely overlaying in panel B. Curves full (ols) and full (ridge) are completely overlaying in panel B.
For further details, see Fig. S29.

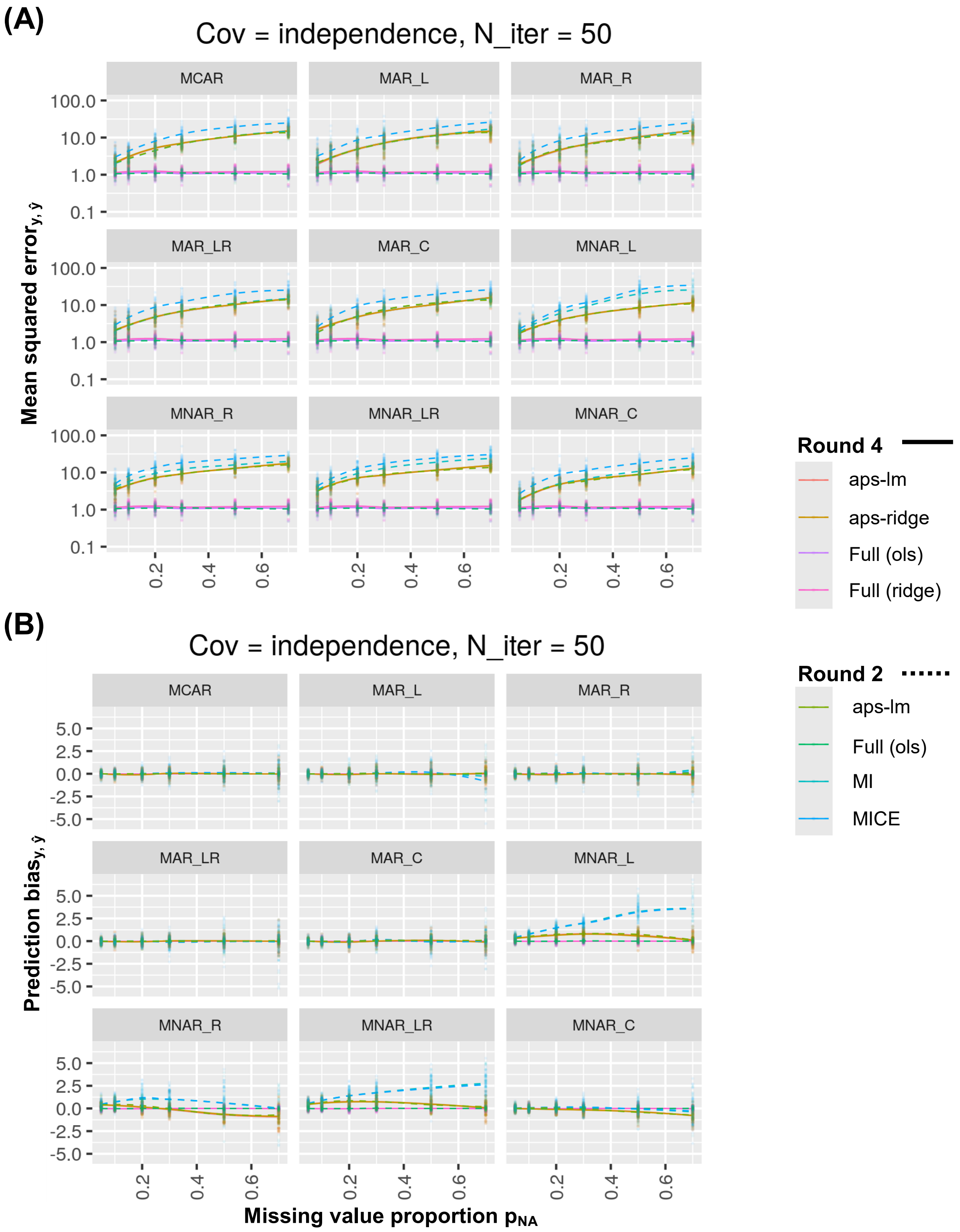


**Figure S31.** Aps-lm, aps-ridge and all imputation approaches for all missing value types under independence. (A) Prediction mean squared error for round 2 and 3-Simulation 1 (overlay between Fig. S25B (dotted) and Fig. S29A (solid)). (B) Prediction bias for round 2 and 3-Simulation 1 (overlay between Fig. S20B (dotted) and Fig. S29B (solid)). For further details, see Fig. S20, S25 and S29.

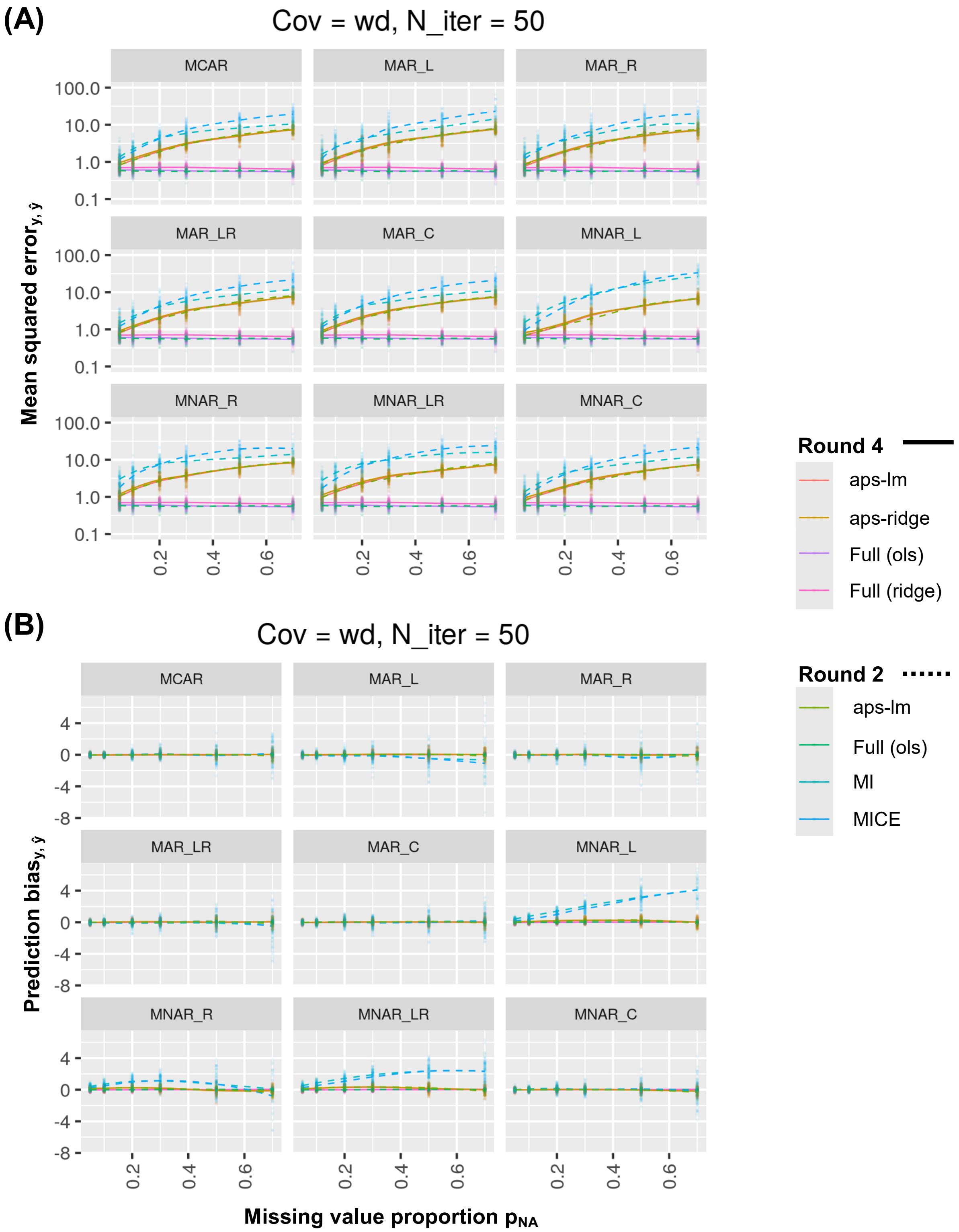


**Figure S32.** Aps-lm, aps-ridge and all imputation approaches for all missing value types under weak dependence. (A) Prediction mean squared error for round 2 and 3-Simulation 2 (overlay between Fig. S26B (dotted) and Fig. 5A (solid)). (B) Prediction bias for round 2 and 3-Simulation 2 (overlay between Fig. 4B (dotted) and Fig. 5B (solid)). For further details, see Fig. 4-5 and S26.

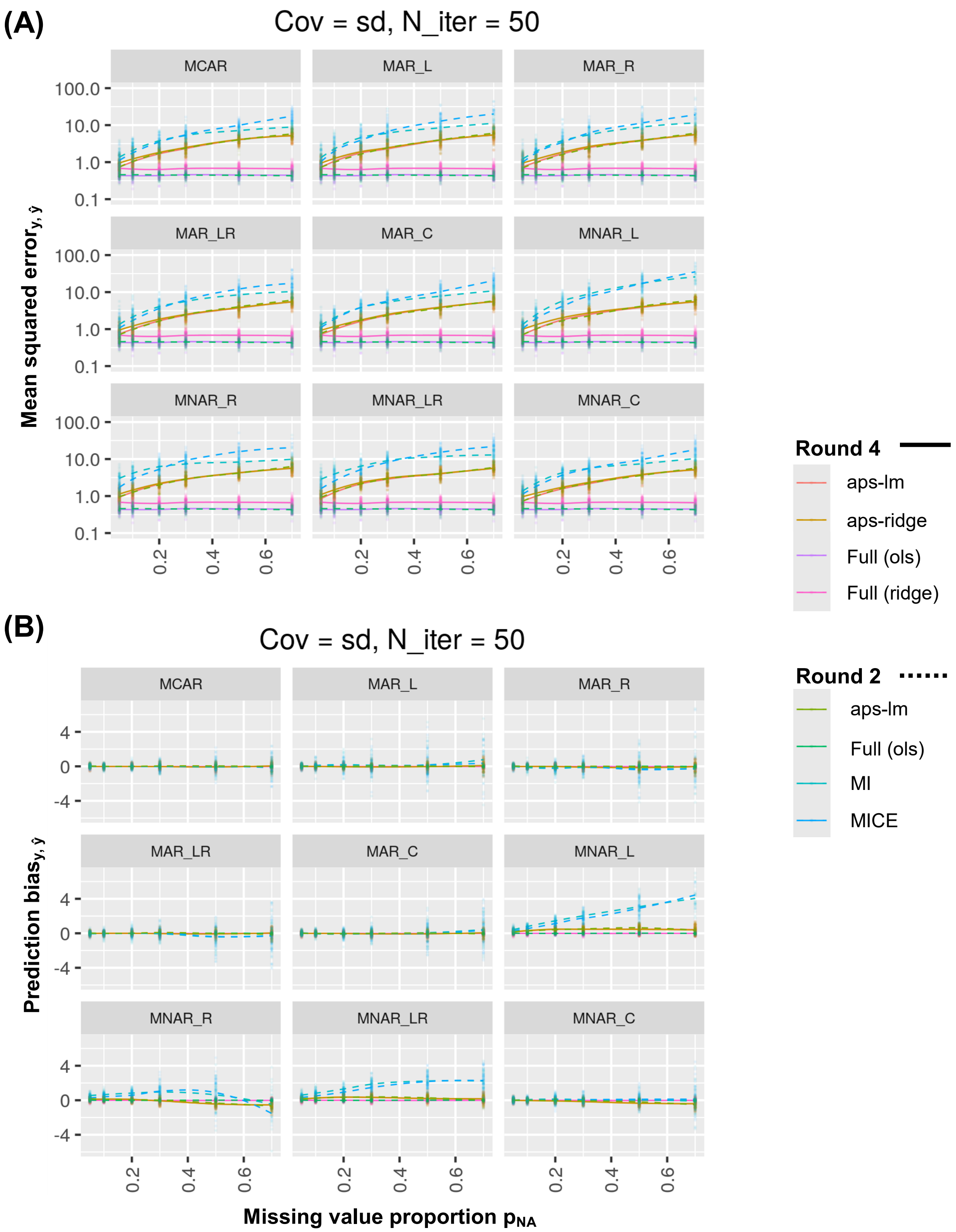


**Figure S33.** Aps-lm, aps-ridge and all imputation approaches for all missing value types under strong dependence. (A) Prediction mean squared error for round 2 and 3-Simulation 3 (overlay between Fig. S27B (dotted) and Fig. S30A (solid)). (B) Prediction bias for round 2 and 3-Simulation 3 (overlay between Fig. 21B (dotted) and Fig. 30B (solid)). For further details, see Fig. S21, S27 and S30.

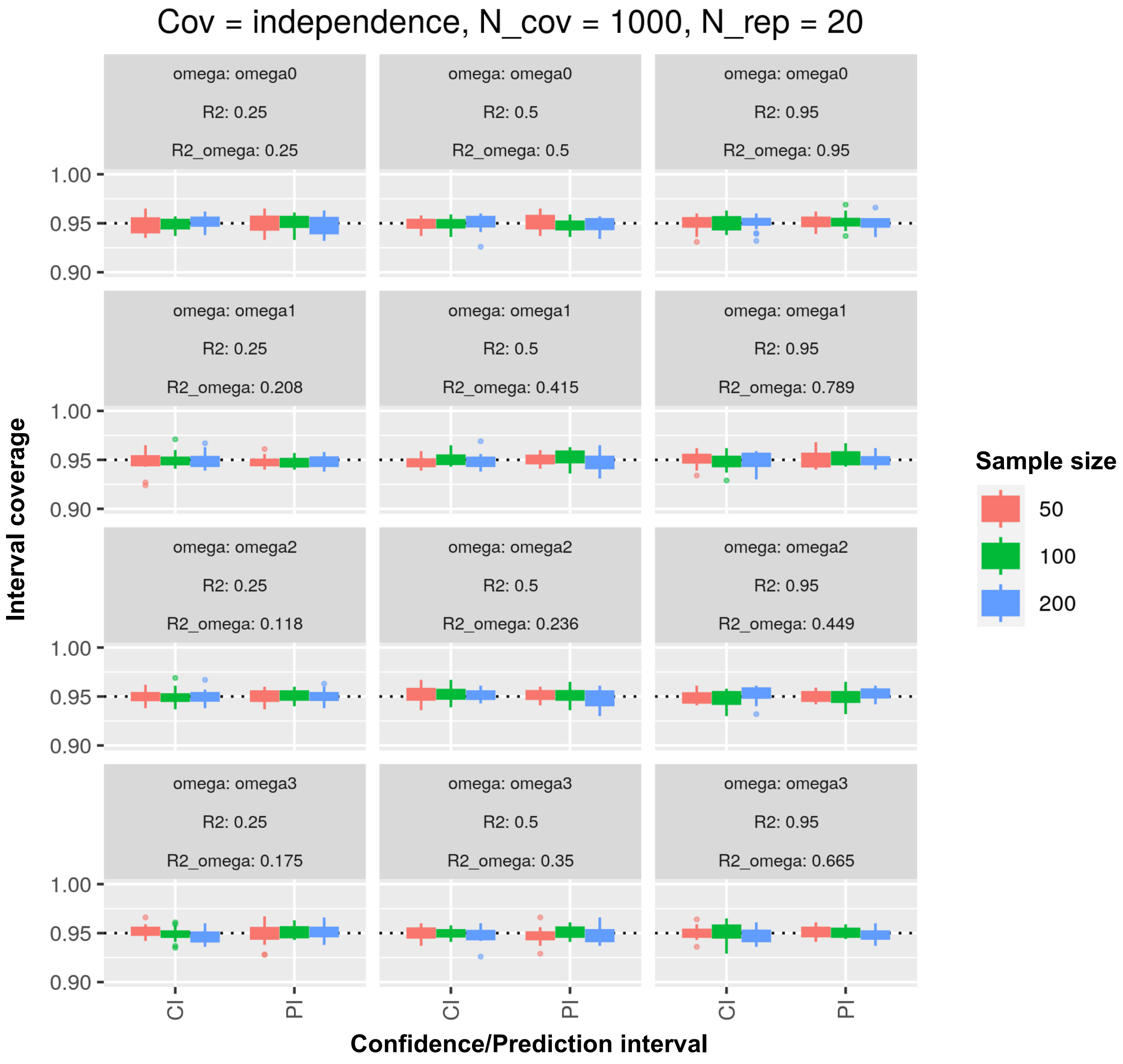


**Figure S34**. Round 4-Simulation 1: aps-lm coverage confidence and prediction intervals under independence for different $R^2$ and sample sizes, n. Four sets of missing value sets are tested: $\Omega_0$={} (omega0), $\Omega_1$={2} (omega1), $\Omega_2$={2,4,7} (omega2) and $\Omega_3$={1,2,3,4,8,9} (omega3). $R^2_\Omega$ (R2_omega) denotes the corresponding $R^2$ of the submodel after excluding covariates in Ω. Boxplots represent the distribution of the interval coverage across replicates ($N_{rep}$ = 20), estimated as the number of iterations at which E[Y|X] (confidence interval) or Y|X (prediction interval) lies within the bounds of the estimated intervals; the total number of iterations employed for each iteration was $N_{cov}$ = 1000. Note that the y-axis begins at 0.90. CI: confidence interval; PI: prediction interval.

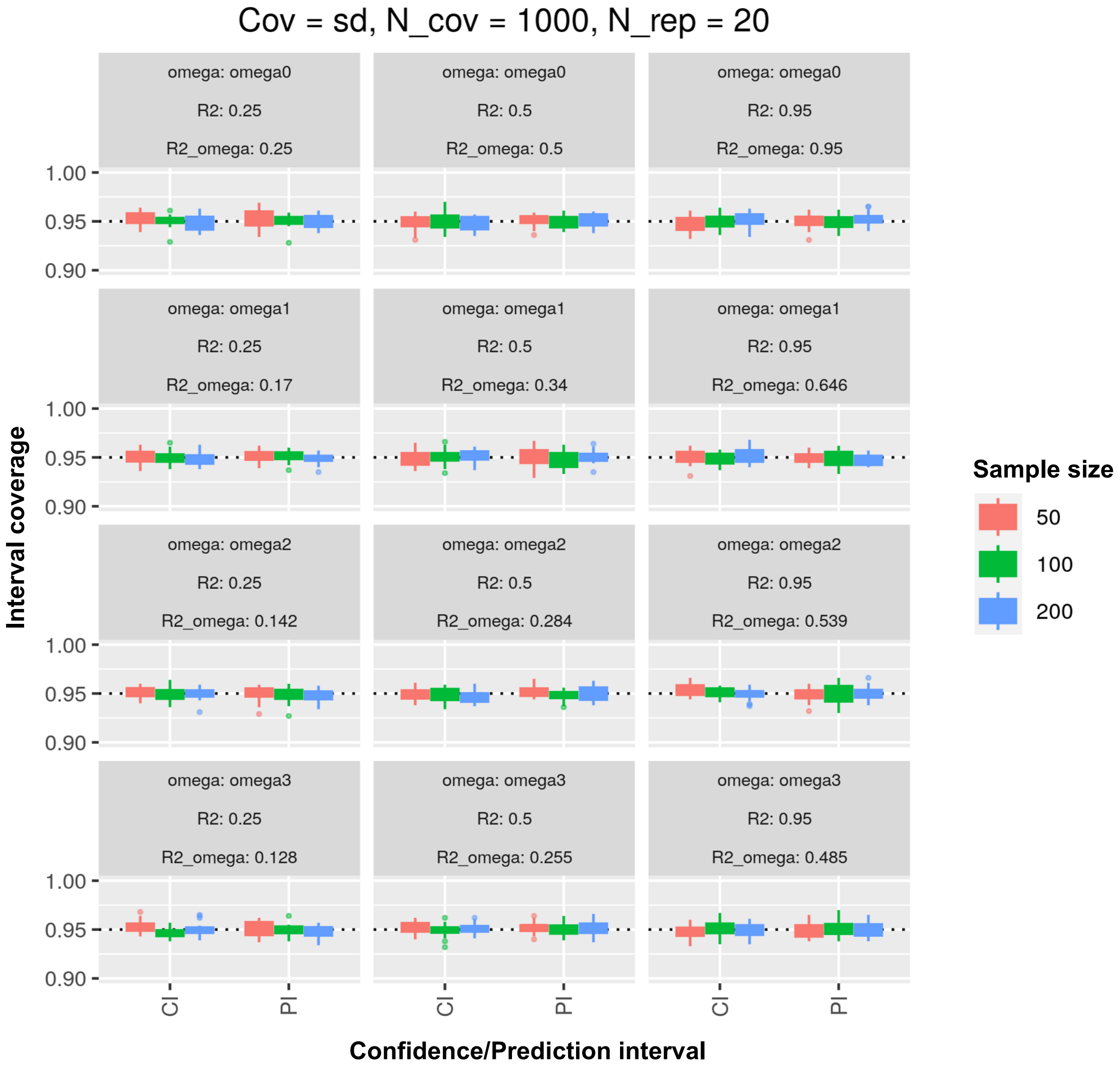


**Figure S35.** Round 4-Simulation 3: aps-lm coverage confidence and prediction intervals under strong dependence for different $R^2$ and sample sizes, n. For further details, see Fig. S34.
